# High-resolution mapping and digital atlas of subcortical regions in the macaque monkey based on matched MAP-MRI and histology

**DOI:** 10.1101/2021.11.23.469706

**Authors:** Kadharbatcha S Saleem, Alexandru V Avram, Daniel Glen, Cecil Chern-Chyi Yen, Frank Q Ye, Michal Komlosh, Peter J Basser

## Abstract

Subcortical nuclei and other deep brain structures are known to play an important role in the regulation of the central and peripheral nervous systems. It can be difficult to identify and delineate many of these nuclei and their finer subdivisions in conventional MRI due to their small size, buried location, and often subtle contrast compared to neighboring tissue. To address this problem, we applied a multi-modal approach in *ex vivo* non-human primate (NHP) brain that includes high-resolution mean apparent propagator (MAP)-MRI and five different histological stains imaged with high-resolution microscopy in the brain of the same subject. By registering these high-dimensional MRI data to high-resolution histology data, we can map the location, boundaries, subdivisions, and micro-architectural features of subcortical gray matter regions in the macaque monkey brain. At high spatial resolution, diffusion MRI in general, and MAP-MRI in particular, can distinguish a large number of deep brain structures, including the larger and smaller white matter fiber tracts as well as architectonic features within various nuclei. Correlation with histology from the same brain enables a thorough validation of the structures identified with MAP-MRI. Moreover, anatomical details that are evident in images of MAP-MRI parameters are not visible in conventional T1-weighted images. We also derived subcortical template “SC21” from segmented MRI slices in three-dimensions and registered this volume to a previously published anatomical template with cortical parcellation (Reveley et al., 2017; Saleem and Logothetis, 2012), thereby integrating the 3D segmentation of both cortical and subcortical regions into the same volume. This newly updated three-dimensional D99 digital brain atlas (V2.0) is intended for use as a reference standard for macaque neuroanatomical, functional, and connectional imaging studies, involving both cortical and subcortical targets. The SC21 and D99 digital templates are available as volumes and surfaces in standard NIFTI and GIFTI formats.

## Introduction

Deep brain structures, such as the basal ganglia, thalamus, hypothalamus, amygdala, and brainstem, are known to play important roles in the regulation of autonomic (sympathetic and parasympathetic), sensorimotor, cognitive, and limbic functions. The anatomical mapping of subcortical regions in macaques and marmosets has enabled the study of the functional activity of these regions within larger brain circuits using fMRI (Baker et al., 2006; Hung et al., 2015; Logothetis et al., 2012; Matsui et al., 2012; Murris et al., 2020; Ortiz-Rios et al., 2015; Schaeffer et al., 2020; Turchi et al., 2018), transcranial focused ultrasound stimulation (Folloni et al., 2019), and optogenetics (Galvan et al., 2012; Stauffer et al., 2016). Continuing advances in neuroimaging have yielded a growing number of useful anatomical contrasts, in addition to the conventional T1- and T2-weighted MRIs, enabling multiparametric mapping of white matter fiber tracts and neuroanatomical regions in the deep brain structures; for review see (Plantinga et al., 2014). In particular, diffusion MRI (dMRI) is a noninvasive preclinical and clinical neuroimaging method that probes tissue microstructure and brain connectivity. dMRI is sensitive to the microscopic motions of water molecules diffusing in tissue and can therefore reflect features of cellularity or myelin content (Pierpaoli and Basser, 1996; Pierpaoli et al., 1996). Signal models of dMRI, such as diffusion tensor imaging (DTI), can measure diffusion in both isotropic and anisotropic tissue, such as white matter, and characterize the underlying microstructure with widely used scalar parameters such as the fractional anisotropy (*FA*) and the mean diffusivity (*MD*) (Basser, 1995; Basser et al., 1994).

Mean apparent propagator (MAP)-MRI (Ozarslan et al., 2013) is now a clinically feasible (Avram et al., 2016) advanced diffusion MRI (dMRI) method that explicitly measures the diffusion propagators (i.e., the probability density function of 3D net displacements of diffusing water molecules) using an efficient analytical series approximation. The measured propagators provide a complete description of the diffusion processes in tissues and can be used to derive microstructural parameters (Avram et al., 2017) obtained with other methods such as DTI or diffusion kurtosis imaging (DKI). Thus, MAP-MRI subsumes other dMRI methods, such as DTI, providing parameters like the fractional anisotropy (*FA*), the mean, axial, and radial diffusivities (*MD*, *AD*, and *RD*, respectively), or the DEC maps (Pajevic and Pierpaoli, 1999). Most importantly, MAP-MRI yields a family of new microstructural parameters that quantify important properties of the diffusion propagators, such as zero-displacement probabilities, non-gaussianity (*NG*), and propagator anisotropy (*PA*) (Avram et al., 2014a; Avram et al., 2018a; Avram et al., 2017). These scalar parameters describe tissue water mobility more efficiently and comprehensively than conventional dMRI methods (Hutchinson et al., 2018) providing excellent contrast in both gray and white matter. For example, the return-to-origin probability (*RTOP*) that quantifies the probability of zero net displacements, is affected by the presence of microscopic barriers and hindrances (e.g., cell membranes and filaments, respectively). The return-to-axis and return-to-plane probabilities (*RTAP* and *RTPP*, respectively) decompose the *RTOP* with respect to the principal diffusion directions providing useful information about pore size and shape in anisotropic tissues. The Non-Gaussianity index (*NG*) quantifies the deviation of the propagator from a homogeneous Gaussian (free) diffusion process reflecting diffusion heterogeneity (i.e., the presence of microscopic water pools with different diffusivities). The propagator anisotropy (*PA*) generalizes the *FA* and quantifies variations in water diffusion along different orientations and is modulated by the shapes and orientations of underlying tissue components. We can also visualize diffusion anisotropy using MAP-derived 3D fiber orientation distribution functions (*fODF*s), and the corresponding direction encoded color [DEC] map (Pajevic and Pierpaoli, 1999). Together the DTI and MAP parameters provide a richer description of tissue microstructure compared to conventional T1 or T2-weighted MRIs and are therefore well-suited for detailed anatomical mapping of cortical and subcortical structures (Avram et al., 2020a; Avram et al., 2020b; Saleem et al., 2020). Recent studies suggest that dMRI is an important imaging contrast for delineating deep brain anatomy and mapping structural connectivity in humans (Calabrese et al., 2015b; Oishi et al., 2020).

The comprehensive high-resolution 3D MRI-histology based atlas of subcortical regions in the macaque monkey is of great use in its application to a project that involves clinical, functional, or anatomical studies. In particular, it is of immediate value to register the 3D atlas to a given macaque brain MRI scan in order to determine the potential target for deep brain stimulation (DBS) in the macaque model of neurological disorders, the areal location of fMRI responses, or the regions-of-interest for anatomical tracer injections (connectome studies). Several studies provide MRI-based atlases in humans, including *in vivo* or *ex vivo* anatomical delineation of deep brain structures with ultrahigh-resolution MRI (Abosch et al., 2010; Deistung et al., 2013a; Deistung et al., 2013b; Ewert et al., 2018; Hoch et al., 2019a; Hoch et al., 2019b; Keuken et al., 2014; Lenglet et al., 2012; Pauli et al., 2018; Rijkers et al., 2007). In contrast, a limited number of studies have done the detailed mapping of subcortical regions or the creation of a subcortical atlas using MRI in NHP, macaque: 1) INIA19, a cortical and subcortical template atlas was created from high-resolution T1-weighted MRI of 19 macaques for imaging-based studies of NHPs (Rohlfing et al., 2012). The authors of that study pointed out that the segmentation of the hypothalamus, amygdala, and basal forebrain in their maps is incomplete. Regions in the brainstem (midbrain and hindbrain) were also not fully described. 2) Calabrese and colleagues (Calabrese et al., 2015a) presented an MRI-DTI based atlas of rhesus macaque brain based on 10 postmortem brain specimens. It provided detailed three-dimensional segmentation of major cortical areas and white matter pathways, but a limited number of subcortical regions (basal ganglia, thalamus, amygdala but no brainstem parcellation). 3) D99 (Reveley et al., 2017), a high-resolution 3D digital template atlas of the rhesus macaque brain based on the Saleem and Logothetis atlas, was mainly focused on a complete parcellation of cortical areas, and also provided a template for a limited number of subcortical targets. 4) A recent study created a subcortical atlas of the rhesus macaque or SARM (Hartig et al., 2021) based on an *ex vivo* structural scan of a single subject, but it relied on histological sections obtained from a different subject (Paxinos et al., 2009) to delineate subcortical regions on *ex vivo* MR images.

Because subcortical structures are usually concentrated in a small region of the brain and have limited contrast on conventional MRI, imaging detailed neuroanatomy with T1- and T2-weighted scans alone without the corresponding matched histological information from the same animal is difficult and unreliable. As shown in the results section below, the combination of different MRI parameters with high-spatial resolution (200 μm), aided by histological information is pivotal in delineating nuclei and fiber tracts in deep brain structures, including sub-structures and laminae, e.g., in the thalamus. A complete 3D brain segmentation based on MRI-histology correlations is critical to assist neuroanatomical and neuroimaging studies in macaques, in particular, and primates in general.

Here, we combine high-resolution MRI data, including conventional MRIs and DTI/MAP parameters, obtained at 7T with 9 microstructural parameters, and five histological stains visualized with high-resolution microscopy in the same specimen to map the location, boundaries, and microarchitectural features of subcortical regions, and associated white matter pathways of the macaque monkey *ex vivo*. This integrated multi-modal approach has produced a more objective and reproducible delineation of gray and white matter regions and their boundaries in subcortical targets, including the basal ganglia, thalamus, hypothalamus, limbic region (amygdala), basal forebrain, and the rostrocaudal extent of the brainstem (midbrain, pons, and medulla). In addition, the 3D information for subcortical regions generated within the DTI/MAP parameters were registered to a standard D99 digital atlas (Reveley et al., 2017). This new atlas is designed to provide a practical standard template for region definition of both cortical and subcortical targets in the NHP brain.

## Materials and methods

### Perfusion fixation

An adult male rhesus monkey (*Macaca mulatta*), weighing 13.55 kg., was perfused for the *ex vivo* MRI and histological studies. All procedures adhered to the Guide for the Care and Use of Laboratory Animals (National Research Council) and were carried out under a protocol approved by the Institutional Animal Care and Use Committee of the National Institute of Mental Health (NIMH) and the National Institute of Health (NIH). The animal was deeply anesthetized with sodium pentobarbital and perfused transcardially with 0.5 liters of heparinized saline, followed by 4 liters of 4% paraformaldehyde, both in 0.1 M phosphate buffer (pH 7.4). After perfusion, the brain was removed from the cranium, photographed, and post-fixed for 8h in the same buffered paraformaldehyde solution. Following the post-fixation, the brain was transferred into 0.1 M phosphate-buffered saline (PBS) with sodium azide before the MRI data acquisition.

### Ex vivo MRI

#### Data acquisition

A 3D structural MRI scan was obtained from the fixed *ex vivo* brain specimen. The MRI volume was centered inside a virtual cylindrical enclosure (68 mm in diameter) with the anterior posterior direction aligned along the axis of the cylinder and manually oriented to match the stereotaxic plane of the D99 atlas, defined by the standard anatomical interaural axis and orbital ridges (Reveley et al., 2017; Saleem and Logothetis, 2012). The oriented MRI volume and cylindrical structure were used to 3-D print a custom-made cylindrical brain mold for accurately positioning the fixed brain specimen in the MRI scanner. The 3D mold containing the fixed brain specimen (Fig. 1, inset) was placed inside a custom 70 mm diameter cylindrical container which was filled with fomblin and gently stirred under vacuum for 4 h to remove air bubbles around the brain. Subsequently, the container was sealed and prepared for high-resolution DTI/MAP-MRI, MTR, and T2-weighted imaging using a Bruker 7T/300 mm horizontal MRI scanner with a 72 mm quadrature RF coil (Bruker, Billerica, USA). All MRI volumes were eventually registered to D99 atlas using rigid-body and non-linear registration (see below).

**Fig. 1.**
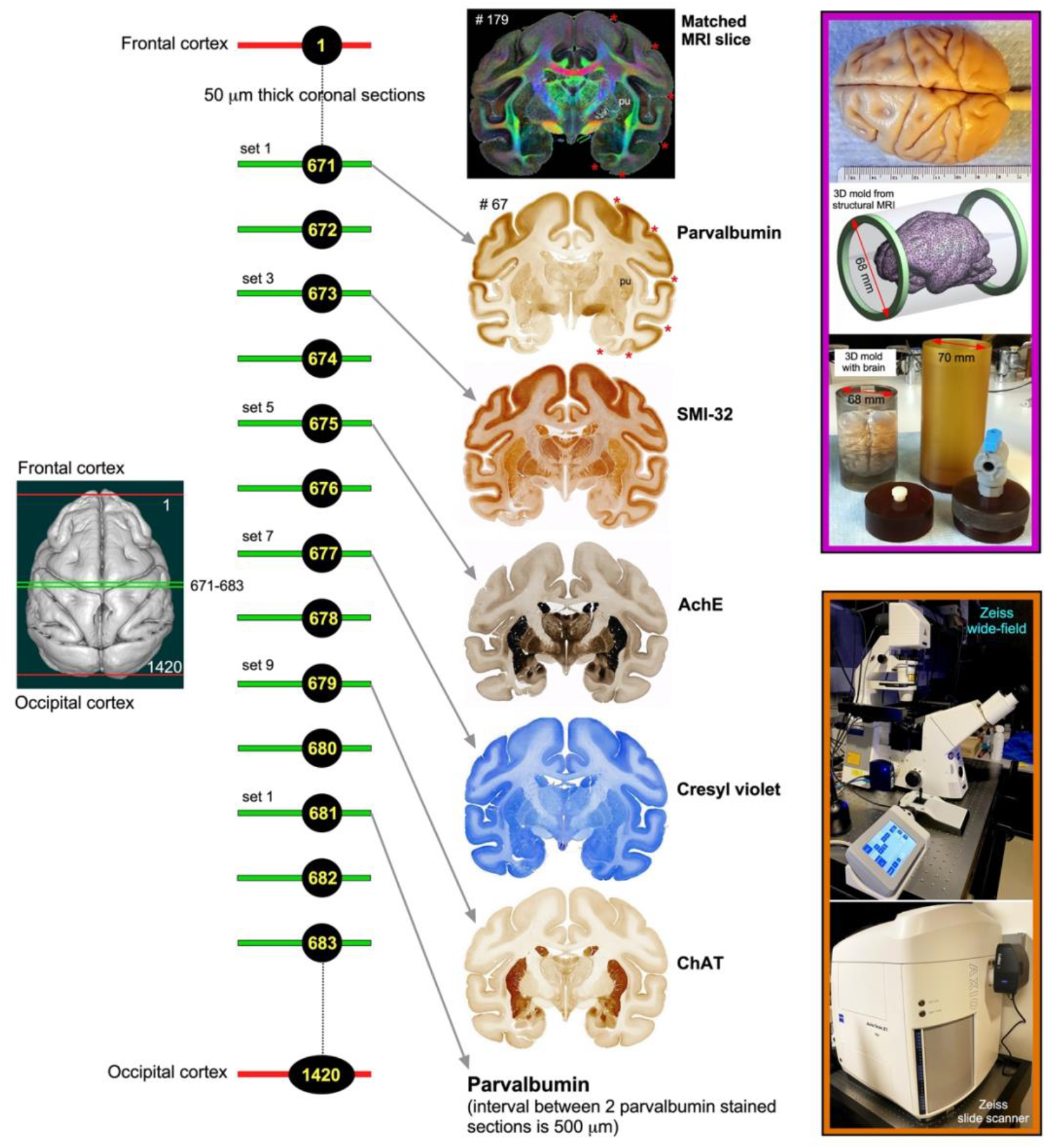
Histological processing, staining, and high-resolution imaging. Frozen sections were cut coronally from the frontal cortex to the occipital cortex at 50μm thickness on a sliding microtome. In total, 1420 sections were collected, but only 50% of the sections were processed with different cell bodies and fiber stains. This example shows an alternating series of sections at the level of the anterior temporal cortex (671, 673, 675, 677, and 679) stained with parvalbumin, SMI-32, AchE, Cresyl violet, and ChAT, respectively. The rostrocaudal locations of these sections are shown on the rendered brain image of this case on the left. This sequence of staining is followed from the frontal to the occipital cortex. We obtained a total of 142 stained sections in each series, and the interval between two adjacent sections in each series is 500μm. The high-resolution images of stained sections were captured using a Zeiss wide-field microscope and a Zeiss high-resolution slide scanner Axioscan (inset on the bottom right). These histology images were then aligned manually with the corresponding MAP/DTI (top) and other MRI parameters of the same specimen to allow visualization and delineation of subcortical structures in specific region-of-interest (see Data analysis section). Note the correspondence of sulci (red stars), gyri, and deep brain structures (e.g., putamen-pu) in both MRI and histology sections. #67 refers to the section number in each set/series of stained sections, and #179 indicates the matched MRI slice number in 3D volume. The unique characteristics of each stain are described in materials and methods. The inset on the top right shows the dorsal view of a perfusion-fixed macaque brain, a custom 3D brain mold generated from a structural MRI of this specimen, and a custom cylindrical container to position the 3D mold with the brain for MR imaging. For other details, see the “data acquisition” section above.

MAP-MRI data were acquired with an isotropic resolution of 200 µm, i.e., a 375x320x230 imaging matrix on a 7.5x6.4x4.6 cm field-of-view (FOV), using a 3D diffusion spin-echo (SE) echo-planar imaging (EPI) sequence with 50 ms echo time (TE), 650 ms repetition time (TR), 8 segments and 1.33 partial Fourier acceleration. A total of 112 diffusion-weighted images (DWIs) were acquired on multiple b-value shells: 100, 1000, 2500, 4500, 7000, and 10000 ms/mm^2^ with diffusion-encoding gradient orientations (3, 9, 15, 21, 28, and 36, respectively) uniformly sampling the unit sphere on each shell and across shells (Avram et al., 2018b; Koay et al., 2012). The diffusion gradient pulse durations and separations were δ=6 ms and Δ=28 ms. We also acquired a magnetization transfer (*MT*) prepared scan using a 3D gradient echo acquisition with 250 µm isotropic resolution (a 312x256x200 imaging matrix on a 7.8x6.4x5.0cm FOV), a 15° excitation flip angle, TE/TR=3.5/37ms. The parameters of the *MT* saturation pulse were 2 kHz offset, 12.5 ms Gaussian pulse with 6.74 µT peak amplitude, 540° flip angle. Two averages were obtained for each of the *MT* on and *MT* off scans. The total duration of the MAP-MRI scan was 93 hours and 20 minutes, and MT scan was 6 hours and 18 minutes.

#### Data processing

All DWIs were processed using the TORTOISE software packages (Pierpaoli et al., 2010) and the MTR as the structural template to correct for Gibbs ringing artifacts, motion (drift), and imaging distortions due to magnetic field inhomogeneities and diffusion gradient eddy currents. The mean apparent propagator was estimated in each voxel by fitting the diffusion data with a MAP series approximation, truncated at order 4 using a MATLAB implementation. We computed DTI parameters, fractional anisotropy (*FA),* mean diffusivity *(MD),* axial diffusivity *(AD),* and radial diffusivity *(RD*), as well as MAP-MRI tissue parameters, propagator anisotropy (*PA),* return-to-origin probability *(RTOP),* return-to-axis probability *(RTAP),* return-to-plane probability *(RTPP),* non-gaussianity *(NG*) and the non-diffusion attenuated image, which provides a T2-weighted contrast. We also estimated in each voxel the orientation distribution function (*ODF*) and fiber ODFs (*fODF*s) and visualized them with the MrTrix3 software package (Tournier et al., 2012). From the DTI model, we computed the linear, planar, and spherical anisotropy coefficients, *CL*, *CP*, and *CS*, respectively, that characterize the shape of the underlying diffusion tensor (Westin et al., 2002). The *MT* ratio (MTR) was computed from the images acquired with and without *MT* preparation. The high contrast between gray and white matter (GM and WM, respectively) in the MTR image allows for reliable image registration to the DWIs. Separately, the D99 was registered to the MTR volume to account for both linear and non-linear deformations using the ANTS software package (Avants et al., 2009). Finally, the inverse of the resulting deformation field was applied to all DTI/MAP parameters (computed from the corrected DWIs in the space of the MTR volume) to transform these parameters to the D99 space (Reveley et al., 2017; Saleem and Logothetis, 2012).

### Histological processing

Following MRI acquisition, the perfusion fixed brain specimen was prepared for histological processing with five different stains as follows. All histological processing (section cutting and staining) was done by FD NeuroTechnologies, Columbia, Maryland. The formaldehyde- perfusion-fixed macaque brain was split into rostral and caudal blocks before freezing and sectioning. The brain blocks were cryoprotected with FD tissue cryoprotection solution™ (FD NeuroTechnologies, Columbia, MD) for 15 days, and then rapidly frozen in isopentane pre-cooled to -70°C with dry ice. The frozen blocks were stored in a -80°C freezer before sectioning.

Serial frozen sections (50-µm thick) were cut on a sliding microtome coronally through the entire brain, including the cerebrum, brainstem, and cerebellum. In total, 1420 sections were collected and sorted into 10 parallel series (142 sections per set) but only five alternating series of sections (1^st^, 3^rd^, 5^th^, 7^th^, and 9^th^ sets) were processed for different cell bodies and fiber stains (Fig. 1). The remaining five sets were stored in a deep freezer with an antifreeze solution. We also collected blockface images of the frozen tissue block at every 250 µm interval.

#### Immunohistochemistry

The sections of 1^st^, 3^rd^, and 9^th^ sets were processed for parvalbumin (PV)-, neurofilament (SMI 32)-, and choline acetyltransferase [ChAT]-immunohistochemistry with the commercially available antibodies (see below). Briefly, after inactivating endogenous peroxidase activity with hydrogen peroxidase, sections were incubated free-floating in 0.01 M phosphate-buffered saline (PBS, pH 7.4) containing 0.3% Triton X-100 (MilliporeSigma, St. Louis, MO), 1% normal blocking serum (Jackson ImmunoResearch, West Grove, PA), and one of the following primary antibodies: mouse monoclonal anti-Parvalbumin antibody (Cat. # P3088, 1:3,000, MilliporeSigma), mouse monoclonal anti-nonphosphorylated neurofilament H (clone SMI 32, Cat. # 801701, 1:1,000, BioLegend, San Diego, CA), and goat anti-choline acetyltransferase antibody (Cat. # AB144P, 1:400, MilliporeSigma) for 67 h at 4°C. The immunoreaction products were then be visualized according to the avidin-biotin complex method (Hsu et al., 1981) using the Vectastin elite ABC kit (Vector Lab., Burlingame, CA) and 3’,3’-diaminobenzidine (MilliporeSigma, St. Louis, MO) as a chromogen. After thorough washes, all sections were mounted on gelatin-coated microscope slides, dehydrated in ethanol, cleared in xylene, and coverslipped with Permount® (Fisher Scientific, Fair Lawn, NJ, USA).

#### Histochemistry

The sections of the 7^th^ set were mounted on gelatin-coated microscope slides and stained with the FD cresyl violet solution™ (FD NeuroTechnologies) for Nissl substance. The sections of the 5^th^ set were processed with the modified acetylcholinesterase [AchE] histochemistry method (Naik, 1963). Briefly, after washing in PBS, sections were pretreated with 0.05 M acetic buffer (pH 5.3) for 3 times, 3 min each. Sections were incubated free-floating in solution A containing sodium acetate, copper sulfate, glycine, ethopropazine, and acetylthiocholine for 3 days. Sections were then incubated in solution B containing sodium sulfide, followed by incubation in solution C containing silver nitrate. Subsequently, sections were re-fixed in 0.1 M phosphate buffer containing 4% paraformaldehyde overnight. All the above steps were carried out at room temperature, and each step was followed by washes in distilled water. After thorough washes in distilled water, sections were mounted on gelatin-coated slides. Following dehydration in ethanol, the sections were cleared in xylene and coverslipped with Permount® (Fisher Scientific).

The histological stains used in this study labeled different types of neuronal cell- or both cell bodies and fiber bundles in cortical and subcortical regions. The calcium-binding protein, parvalbumin (PV) was thought to play an important role in intracellular calcium homeostasis, and the antibody against PV has been shown previously to recognize a subpopulation of non-pyramidal neurons (GABAergic) in the neocortex, and different types of neurons in subcortical structures (Jones, 1998; Jones and Hendry, 1989; Saleem et al., 2007). The SMI-32 antibody recognizes a non-phosphorylated epitope of neurofilament H (Goldstein et al., 1987; Sternberger and Sternberger, 1983) and stains a subpopulation of pyramidal neurons and their dendritic processes in the monkey neocortex (Hof and Morrison, 1995; Saleem and Logothetis, 2012). It is also a valuable stain for a vulnerable subset of pyramidal neurons in the visual, temporal, and frontal cortical areas visualized in postmortem brain of Alzheimer’s disease cases (Hof et al., 1990; Hof and Morrison, 1990; Thangavel et al., 2009). The SMI-32 can also detect axonal pathology in TBI brains (Johnson et al., 2016). The antibody against ChAT recognizes cholinergic neurons and has been a useful stain for motor neurons in the monkey and human brainstem (e.g., cranial nerve nuclei, (Horn et al., 2018). AchE is an enzyme that catalyzes the breakdown of acetylcholine and is shown to be a useful marker for the delineation of different cortical areas (Carmichael and Price, 1994), and major subcortical nuclei in the thalamus and brainstem (Horn et al., 2018; Jones, 1998).

#### Data analysis

The high-resolution images of all stained sections were captured using a Zeiss wide-field microscope and Zeiss high-resolution slide scanner at 5X objective, and these digital images were adjusted for brightness and contrast using Adobe Photoshop CS. These images were then aligned manually with the corresponding images of DTI/MAP parameters along with the estimated T2-weighted (i.e., non-diffusion weighted) and the MTR images to allow visualization and delineation of subcortical structures in specific region-of-interest (ROIs) (e.g., Fig. 4). Some structures like striatum and pallidum were demarcated by comparing them side-by-side on the matched MRI and histology sections, but for others (e.g., thalamus), we used a different approach to delineate their subregions as follows. We first superimposed a histology section onto the matched MRI slice and then manually rotated and proportionally scaled the histology section to match with the outlines of the thalamic subregions on MRI using the transparency function in Canvas X Draw software. Finally, the borders of the subregions were manually traced on the histology sections and translated these traced outlines onto the superimposed underlying MR images using the polygon-drawing tools with smooth and grouping functions in this software (e.g., Fig. 9). These steps were repeated to trace the subregions of the thalamus at different rostrocaudal levels and other deep brain structures.

The histological sections were matched well with the MRIs in this study, and the alignment of these images was only possible with careful blocking of a brain specimen before the histology work. To match the location of deep brain structures in MRI and histology sections, we blocked and sectioned the brain specimen corresponding to the MRI plane as follows (see also MRI Data acquisition and processing sections above). We first visually matched a number of reference points on the MR images with similar locations on the surface of the 3D rendered brain volume generated from these MR images (see Inline Supplementary Fig. 1, top row). The 3D brain volume was manually rotated in the anteroposterior direction to match with the reference points and the orientation of coronal MRI sections. These landmark points on the rendered brain surface were then manually translated onto the corresponding location of the brain specimen of this subject before blocking it (see Inline Supplementary Fig. 1, middle column). These steps enabled us to match the sulci, gyri, and the region of interest (ROI) in deep brain structures in both MRI and histological sections, as shown in Figure 1 and the result section below (see also Inline Supplementary Fig. 1, left and right columns). We did not resample the MRI volume to achieve accurate alignment to the histological images in the current study. A similar approach was used to block a macaque brain specimen in our previous study, but the histology sections did not align well with the MRI in this case. We digitally resliced the structural T1-weighted MRI volume of this specimen slightly to match with the histology sections (See Saleem and Logothetis atlas, 2012, their figures 1.3 and 1.4).

#### Segmentation of subcortical regions and generation of SC21

Subregions (ROIs) of the basal ganglia, thalamus, hypothalamus, amygdala, brainstem, and other deep brain areas, and selected fiber bundles were manually segmented through a series of 247, 200 μm thick coronal sections in *PA*, T2-weighted, or other MRI parameters using ITK-SNAP (Yushkevich et al., 2006). The spatial extent and borders of each segmented region in MRI were confirmed with the matched high-resolution histology images obtained from multiple stained sections (Fig. 1) and with reference to previous studies, as indicated in the result sections below. We then derived the location and spatial relationships among different subcortical regions from these segmented slices in three dimensions (3D) using ITK-SNAP and SUMA (Saad and Reynolds, 2012; Yushkevich et al., 2006). We adapted this new 3D volume with 219 segmented regions as “SC21” (SC stands for subcortical; Fig. 2A, B) and used it for registration to other anatomical *in vivo* and *ex vivo* T1w/MTR volumes from test subjects, described in the following sections.

**Fig. 2.**
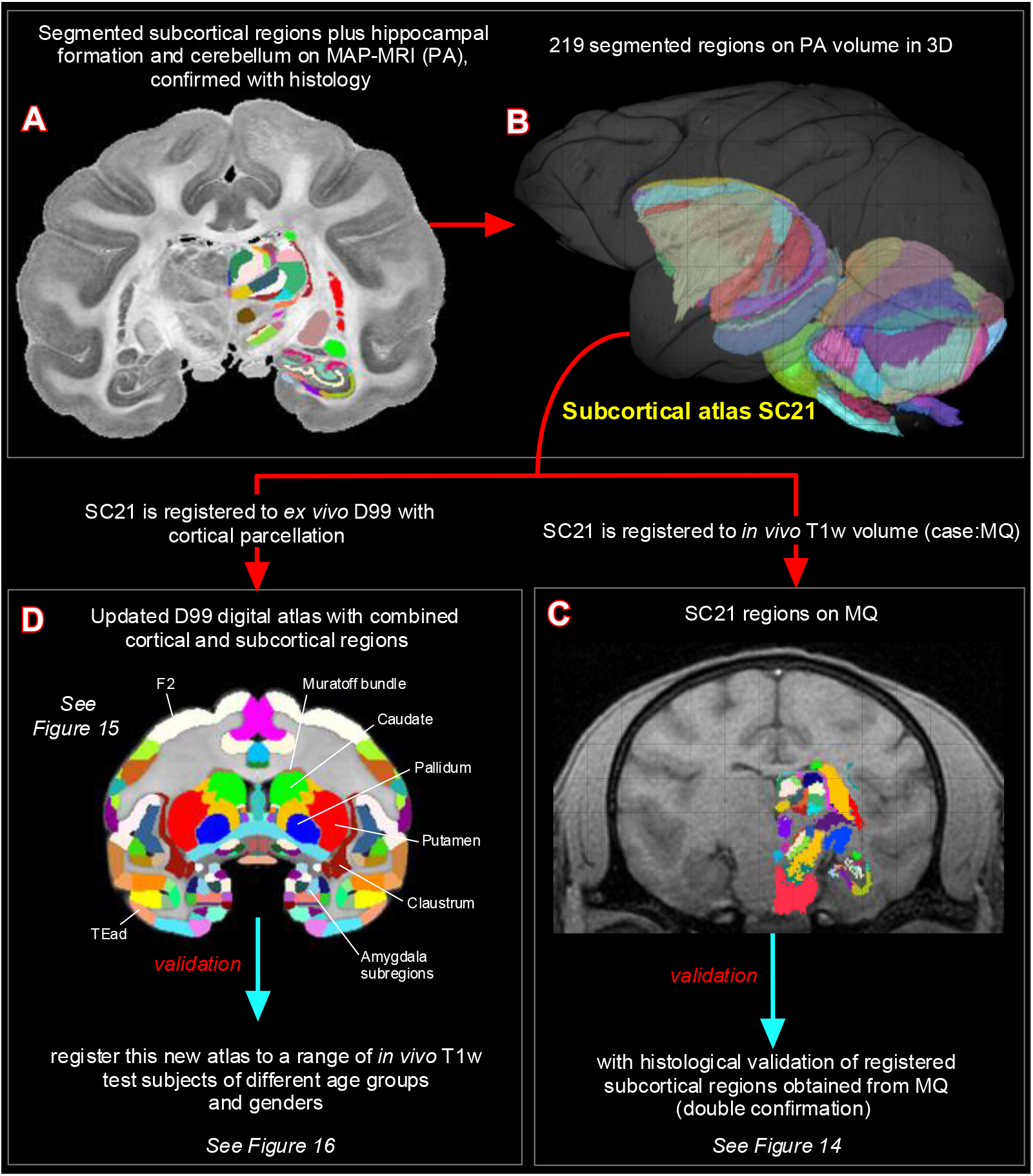
Subcortical segmentation, updated 3D digital template atlas, and registration of 3D atlas to test subjects. A total of 219 deep brain regions, including the hippocampal formation and the cerebellum were manually segmented through a series of 200 μm thick MAP-MRI sections **(A)** using ITK-SNAP and derived the spatial location of these regions in 3D **(B)**. This new MRI-histology based segmented volume (called “SC21”) is registered to *in vivo* T1-weighted MRI volume of a test subject MQ **(C).** The registered subcortical regions on MQ are confirmed again using the corresponding histological sections from the same test subject. The SC21 is also registered to the standard D99 cortical template atlas (Reveley et al., 2017), resulting in an “updated D99 atlas” with combined cortical and subcortical parcellations in the same volume **(D)**. This updated D99 is in turn registered to *in vivo* T1-weighted MRI volumes of 6 other subjects with different age groups. For more details, see figures 14-16.

#### Registration of SC21 to a test subject with histological confirmation of architectonic areas

We registered SC21 to an *in vivo* T1w volume (case, MQ) using affine and nonlinear registration steps in ANTS software package (Avants et al., 2009). The registered SC21 subcortical regions on MQ-MRI slices were confirmed using matched and stained histological sections obtained from MQ (Fig. 2C). Here we validate the accurate delineation of subcortical areas using histology in two cases: First in SC21 and then again in MQ after registration with SC21. For more details, see figure 14 and related text.

#### Updated D99 digital atlas: One volume with combined cortical and subcortical regions

The SC21 subcortical template is also registered to the *ex vivo* MTR D99 surrogate volume (Reveley et al., 2017) using the ANTS software, as described in the previous section. This registered subcortical 3D atlas dataset was integrated into the AFNI (Analysis of Functional NeuroImages; (Cox, 1996; Saad and Reynolds, 2012), and SUMA (Surface Mapper; (Cox, 1996; Saad and Reynolds, 2012) software packages with region labels. To preserve the contiguity of the regions, the transformed subcortical regions were *modally smoothed* with a simple regularization procedure where each voxel was replaced with the most common voxel label in the immediate neighborhood around each voxel (27 voxels). This procedure was used in our earlier D99 atlas (Reveley et al., 2017; Saleem and Logothetis, 2012). The method smooths edges caused by mismatches in 2D drawings applied to a 3D shape. The dataset was then subject to a series of manual verification and correction of areal extent and architectonic borders of different subcortical areas compared with the original segmented SC21 dataset, aided by histology (see above). Additionally, the dataset was tested for “lost clusters” with AFNI’s @ROI_decluster. In this procedure, each region was clustered to a minimum of half the total voxels for that label. The declustered dataset was compared to the original input volume, and differences were manually corrected. Finally, we combined the cortical plus claustrum, hippocampal, and amygdaloid segmentations from the original D99 atlas (Reveley et al., 2017) to this subcortical template, which resulted in both cortical and subcortical parcellations in the same volume (Fig. 2D). All the combined regions were manually checked and corrected again for the right side of the brain. The right-side brain atlas was mirrored around its x-axis (3dLRflip) and the results were combined to create a symmetric brain. This newly updated symmetric D99 digital atlas is now available in the AFNI and SUMA analysis packages to register and apply to the brains of other individual macaques, to guide a number of research applications for which accurate knowledge of areal boundaries is desirable. See figure 15 and related text in the results section.

#### Registration of D99 atlas to test subjects

We registered this updated D99 atlas to *in vivo* T1 MRI volume of 6 individual animals of different age groups using the @animal_warper program in AFNI (Jung et al., 2021). The data set was aligned to the D99 template using center-shifting, affine, and nonlinear warp transformations. The inverted transformations were combined and applied to the atlas to bring the atlas segmentation to the native space of each macaque. The default modal smoothing was applied here to replace each voxel with the most common neighbor in the immediate 27-voxel neighborhood. No histology information is available for these 6 cases. For more details, see the results section and related figure 16. The MR scanning methods to obtain high-contrast T1- weighted images of these 6 animals plus case MQ are described in the next section.

#### High-Contrast in vivo anatomical MRI scans

In 7 normal and healthy animals (age: 1.2–14.8 years), weighing between 2.55 and 5.5 kg, MR anatomical images were acquired in a 4.7 T/400 mm horizontal MRI scanner (Bruker, Billerica, USA) using an MDEFT method. Each monkey was anesthetized with isoflurane and placed into the scanner in a sphinx position with its head secured in a holding frame. A single loop circular coil with a diameter of 14–16.5 cm was placed on top of each animal’s head. The whole-brain MDEFT images were acquired in a 3D volume with a field of view 96x96x70 mm^3^, and 0.5mm isotropic voxel size. The read-out had an 11 ms repetition time, a 4.1ms echo time, and a 11.6° flip angle. The MDEFT preparation had a 1240ms pre-inversion time, and a 960ms post-inversion time for optimized T1 contrast at 4.7 T. Each 3D volume took 25.5 min to acquire without averaging. Most of the scans were acquired with 2 averages and took 51 min. These cases were illustrated in Figures 14 and 16. All these cases were used in our previous study (Reveley et al., 2017).

## Results

Using a combined MRI and histology, we identified and segmented 190 gray matter subregions in the deep brain structures, including the basal ganglia, thalamus, hypothalamus, brainstem (midbrain, pons, and medulla), amygdala, basal forebrain, and the bed nucleus of stria terminalis. It should be noted that the 190 segmented gray matter regions also include architectonically and functionally distinct non-subcortical regions such as different lobules of the cerebellar cortex and the hippocampal formation (for more details, see Figs. 10, 13). In addition, we also distinguished and segmented 29 fiber tracts of different sizes and orientations associated with the basal ganglia, thalamus, brainstem, and cerebellum (see Inline suppl. Table 1). This newly segmented volume is called SC21 or subcortical 21. Figure 2B illustrates the lateral view of SC21 with segmented subcortical regions in 3D, superimposed on the rendered brain volume from this case. Although we delineated 219 regions, it is beyond the scope of this study to describe the detailed anatomy and illustrate all the identified regions in this report. In the following sections (see also supplementary data), we first describe the spatial location, boundaries, and microarchitectural features of selected segmented subcortical regions, and associated white matter fiber pathways from SC21, identified with multiple MRI markers and histological stains, and each of which has provided useful distinctions between subregions. We then validate the anatomical accuracy of delineated subcortical areas by registering SC21 to test subjects, including D99 (“updated D99”). Finally, we test the validity of the updated D99 atlas with combined cortical and subcortical maps on multiple test subjects of different age groups.

### MRI markers **(Fig. 3)**

MAP-MRI and other MRI parameters showed different gray and white matter contrast outside the cerebral cortex. In particular, *PA* or *PA* with fiber orientation distribution functions (*fODFs*) and the corresponding direction encoded color (DEC) map (Pajevic and Pierpaoli, 1999), *RD*, *NG*, *RTAP*, as well as T2-weighted images revealed sharp boundaries and high contrast in the deep brain structures, resulting in a clear demarcation of anatomical structures such as nuclei (e.g., globus pallidus-GP, the anterior ventral nucleus of thalamus-AV), and fiber tracts in subcortical regions (Fig. 3). All qualitative findings observed in MRI were confirmed using adjacent and matched histology sections with several stains (see materials and methods). For convenience, we combined both terminologies: *fODFs* derived directionally encoded color (DEC) map and abbreviated as DEC-FOD in the text and figures.

**Fig. 3.**
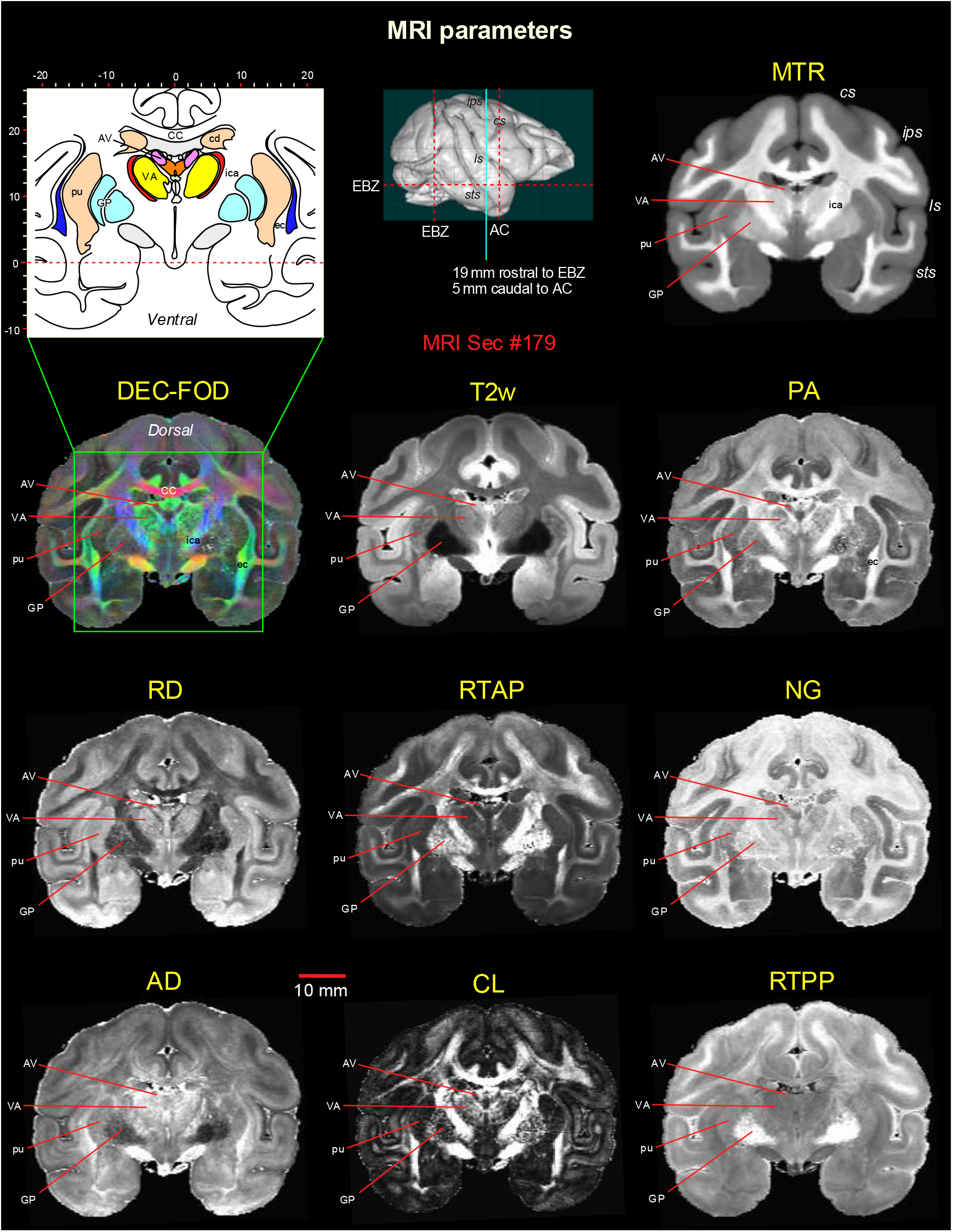
Subcortical regions in different MRI parameters. A coronal MR image from eight DTI/MAP-MRI parameters, T2- weighted (T2w), and the magnetization transfer ratio (MTR) images show four selected subcortical regions: two thalamic nuclei (AV and VA), and two basal ganglia regions (pu and GP). These areas are also illustrated in the corresponding drawing of the section on the top left (inset). This MRI slice is located 19 mm rostral to the ear bar zero (EBZ) or 5 mm caudal to the anterior commissure (AC) as illustrated by a blue vertical line on the lateral view of the 3D rendered brain image. Note that the contrast between these subcortical areas is distinct in different MRI parameters. ***Abbreviations: DTI/MAP-MRI parameters:*** *AD*-axial diffusivity; *CL*-linearity of diffusion from the tortoise; DEC-FOD-directionally encoded color-fiber orientation distribution; *NG*- non-gaussianity; *PA*-propagator anisotropy; *RD*-radial diffusivity; *RTAP*-return to axis probability; *RTPP*-return to plane probability. ***Subcortical regions:*** AV-anterior ventral nucleus; ec-external capsule; ica-internal capsule, anterior limb; VA-ventral anterior nucleus; GP-globus pallidus; pu-putamen. ***Sulci:*** cs: central sulcus; ips-intraparietal sulcus; ls-lateral sulcus; sts-superior temporal sulcus.

Delineation of selected subcortical gray and white matter regions

### Basal ganglia

#### Striatum **(Fig. 4)**

The basal ganglia extend from the prefrontal cortex to the midbrain and are composed of four major groups of nuclei: the striatum (caudate nucleus-cd, putamen-pu, and ventral striatum), pallidum, subthalamic nucleus [STN], and substantia nigra (SN). The spatial location, overall extent, and neighboring relationship between these structures can be visualized in 3D, as shown in Figure 4M. Both cd and pu exhibited similar hypo- or hyperintense contrast in MAP-MRI (*PA*/DEC-FOD*, RD, NG, RTPP*), T2w and MTR images (Fig. 4A-F), but this contrast is strikingly the opposite of the signal intensity found in the surrounding white matter regions such as the anterior limb of the internal capsule (ica), corpus callosum (CC), and external capsule (ec). The location and well-defined contour of these structures, including bridges connecting cd and pu in MRI are matched well with the corresponding histology sections stained with the AchE and PV (Fig. 4G, H). The ventral striatum (Haber, 2003; Heimer et al., 1982) includes the nucleus accumbens (NA) and the olfactory tubercle (OT). The NA, indicated within the red dashed outline in figure 4A-F shows similar contrast with the neighboring ventral portion of the cd and pu in all illustrated MR images. The border between NA and cd/pu is barely visible in any MRI or histological images. In RD, a hyperintense region was found on the dorsomedial part of the NA, which coincided with the dark and intensely stained regions in PV (arrows in Fig. 4C and H), and might correspond to the shell region of the nucleus accumbens (Haber, 2003; Heimer et al., 1982). For simplicity, we labeled the ventral striatal region as NA in all illustrations and 3D digital template atlases. In addition, all subregions of the striatum reveal mosaic-like patterns with bright and dark contrasts in several MRI parameters. They are most prominent in T2w images (Fig. 4A-B, I-J). This architectonic pattern did not match closely with light and dark compartments observed in adjacent SMI-32 and PV-stained sections, as shown by lightly stained regions from the SMI-32 section superimposed on the MR images (Fig. 4I-K; blue outlined regions).

**Fig. 4.**
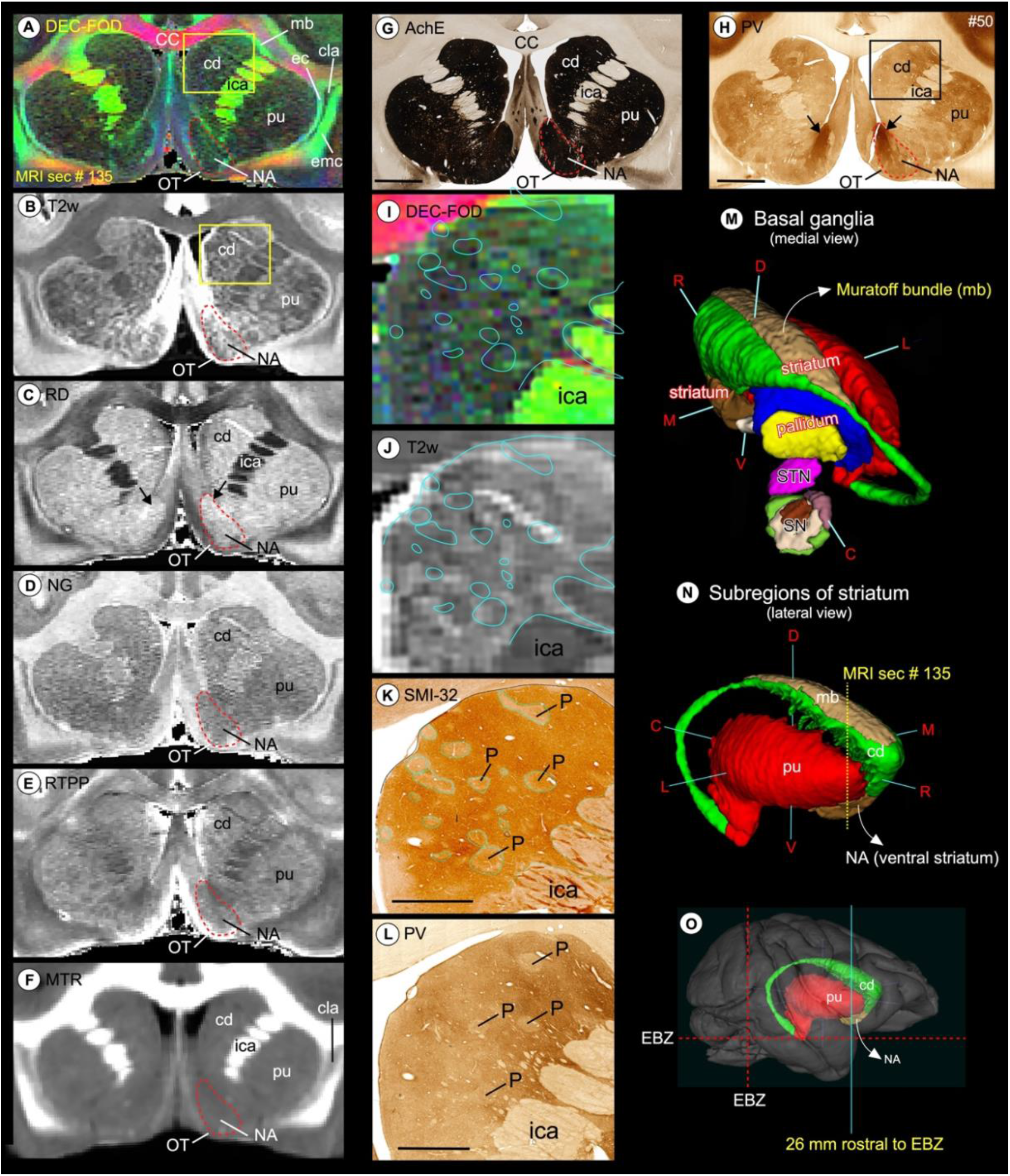
Striatum. Subregions of the striatum (caudate and putamen), and ventral striatum (nucleus accumbens and olfactory tubercle) in coronal MAP-MRI (DEC-FOD, *RD*, *NG*, *RTPP*), T2w, and *MTR* images **(A-F)**, and corresponding matched histology sections stained for AchE and PV **(G-H)**. Note the sharp contrast of these subregions from the surrounding white matter structures in MRI that corresponded well with the architectonic regions in histology sections, including the bridges connecting the caudate and putamen (compare A and G). The red-dashed outline indicates the nucleus accumbens and black arrows in H shows the dark and patchy staining within the medial portion of NA in PV stained section. **(I, J)** The high-power images of the head of the caudate nucleus (cd) from DEC-FOD and T2w images (yellow box in A, B) show mosaic-like patterns with bright and dark contrasts. The blue outlines on the MR images indicate the lightly stained regions, which are traced out from the SMI-32 stained section in K. These regions are neurochemically defined compartments called patches or striosomes. The striosomes did not match closely with bright regions in the corresponding T2w or DEC-FOD images (I, J). The spatial location and overall extent of the striatum with other subregions of the basal ganglia are also illustrated in 3D, reconstructed using ITK-SNAP (M, N), and on the rendered brain image from this case (O). The coronal slice is located 26 mm rostral to the ear bar zero (EBZ) at the frontotemporal junction or limen insula (a vertical blue line in O). #50 in G and H refers to the section number in each set/series of stained sections, and #135 indicates the matched MRI slice number in 3D volume. ***Abbreviations:*** CC-corpus callosum; cd-caudate nucleus; cla-claustrum; ec-external capsule; emc-extreme capsule; ica-internal capsule, anterior limb; mb-Muratoff bundle; NA-nucleus accumbens; OT-olfactory tubercle; P-patch or striosomes; pu-putamen; SN-substantia nigra; STN- subthalamic nucleus. ***Orientation on 3D:*** D-dorsal; V-ventral; R-rostral; C-caudal; M-medial; L-lateral. Scale bar: 5 mm (G, H) and 2 mm (K, L).

#### Pallidum **(Fig. 5)**

The pallidum is divided into the external and internal segments of the globus pallidus (GPe and GPi, respectively), and the ventral pallidum (VP) (Parent, 1990) that are readily identifiable as dark regions on the T2w and *RD*, bright structures on *RTPP*, and granular or gray regions on DEC-FOD and *NG* images (Fig. 5A-E; VP is not visible at this rostrocaudal level). The pallidal subregions are easily distinguished from the laterally adjacent putamen (pu), which shows the opposite signal intensity of GPe and GPi in different MRI parameters (Fig. 5A-F). These features corresponded well with the staining differences observed in the matched histology sections: lightly stained GPe/GPi and darkly stained pu in AchE and ChAT-stained sections, or vice-versa in SMI 32 stained section (Figure 5G-I). A clear separation (or lamina) between the putamen and pallidum or within the pallidum can be visualized in different MRI parameters. The lateral medullary lamina (lml) and medial medullary lamina (mml) that separate pu from GPe, and GPe from GPi, respectively, are visible on the DEC-FOD, *NG*, and *MTR* images (Fig. 5A, D, F; red arrows) but this separation is less distinct or barely visible in other MRI parameters. There is no lamina within the GPi but the lateral and medial subregions of the GPi are distinguished based on dark and light honeycomb-like neuronal processes, respectively, in AchE and SMI-32 stained sections but not in ChAT or MRI (Fig. 5G-I, magnified regions).

**Fig. 5.**
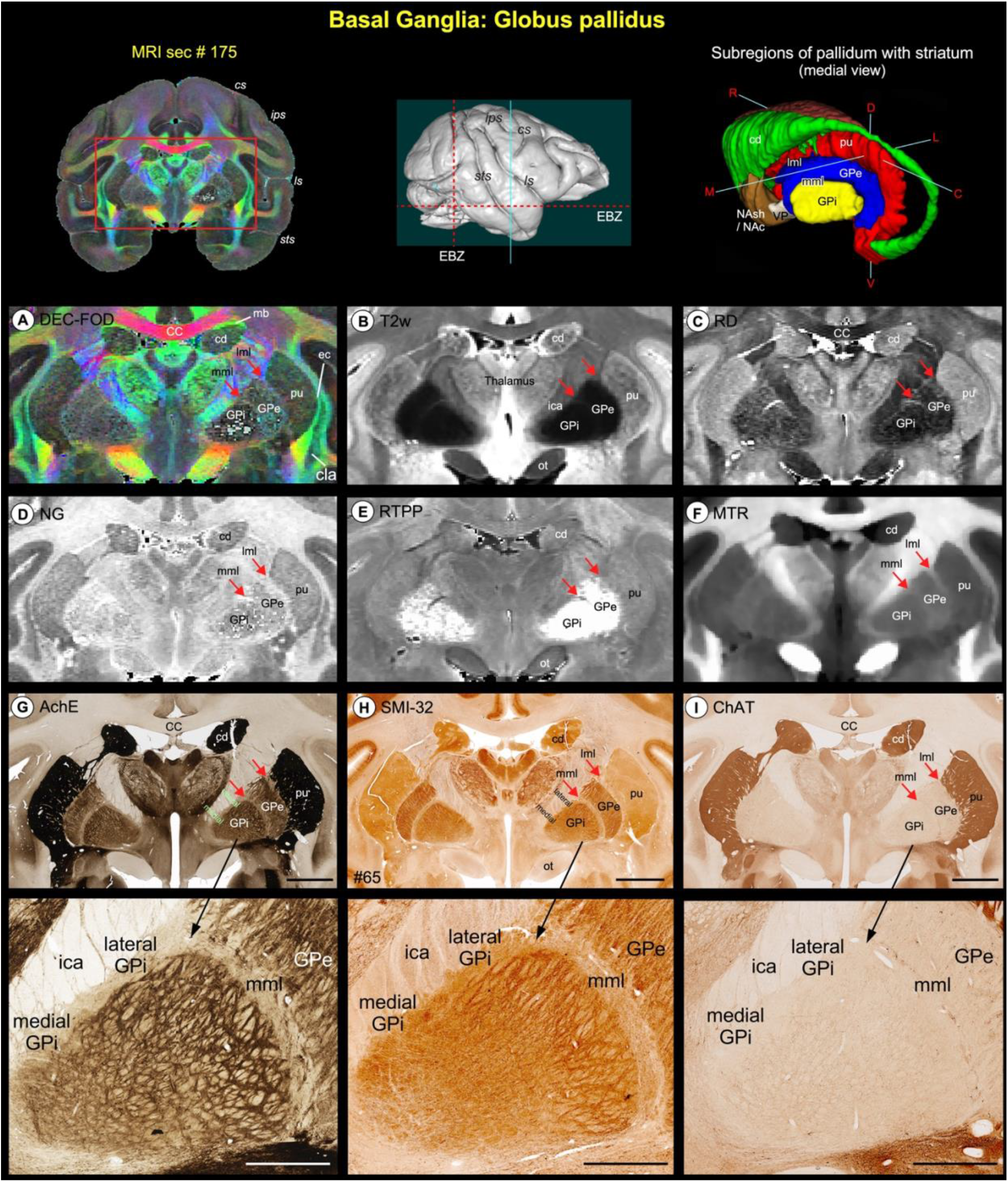
Pallidum. **(A-F)** illustrate the zoomed-in view of two subregions of the pallidum: globus pallidus external segment (GPe) and globus pallidus internal segment (GPi), medial to the putamen (pu) on the coronal MAP-MRI (DEC-FOD, *RD*, *NG*, *RTPP*), T2w, and *MTR* images. **(G-I)** show the same subcortical regions in the corresponding histology sections stained with AchE, SMI-32, and ChAT (sec #65). The rostrocaudal extent of this coronal slice in the anterior temporal cortex (green vertical line on the rendered brain image), and the spatial location and overall extent of the pallidum with other subregions of the basal ganglia in 3D are shown on the top. Note that both GPe and GPi are readily identifiable as hypointense regions on the T2w and *RD*, and hyperintense on *RTPP*, compared with the laterally adjacent pu and medially located thalamus and the internal capsule (ica). Red arrows indicate the location of lateral medullary lamina (lml) between pu and GPe, and the medial medullary lamina (mml) between GPe and GPi. These laminae are visible on the DEC-FOD, *NG*, and *MTR* images but less distinct or barely visible in other MRI parameters. In contrast to AchE and SMI-32, these laminae are not distinguishable in the ChAT-stained section (G-I). Unlike human brains, there is no separate lamina within the GPi in macaque monkeys but mediolateral subregions within this pallidal region can be distinguished based on the dark- and light honeycomb-like structures in AchE and SMI-32 (magnified regions in G and H). ***Abbreviations:*** CC-corpus callosum; cd-caudate nucleus; cla-claustrum; ec-external capsule; ica-internal capsule anterior limb; mb-Muratoff bundle; ot-optic tract; pu-putamen. ***Sulci:*** cs-central sulcus; ips-intraparietal sulcus; ls-lateral sulcus; sts-superior temporal sulcus. ***Orientation on 3D:*** D-dorsal; V-ventral; R-rostral; C-caudal; M-medial; L-lateral. Scale bar: 5 mm (G-I); 2 mm applies to magnified regions from G-I (black arrows).

#### Substantia nigra **(Fig. 6)**

The substantia nigra has been divided into two to four subregions by different studies based on the Nissl, myelin, AchE, tyrosine hydroxylase (TH), endogenous iron stains, or staining specific to dopamine cells in the midbrain (Arsenault et al., 1988; Francois et al., 1985; Jimenez-Castellanos and Graybiel, 1987). These subregions are the substantia nigra- pars reticulata (SNpr), pars compacta (SNpc), pars lateralis (SNpl), and pars mixta (SNpm). Unlike striatum and pallidum, it is difficult to identify all subregions of SN with one MAP-MRI parameter.

**Fig. 6.**
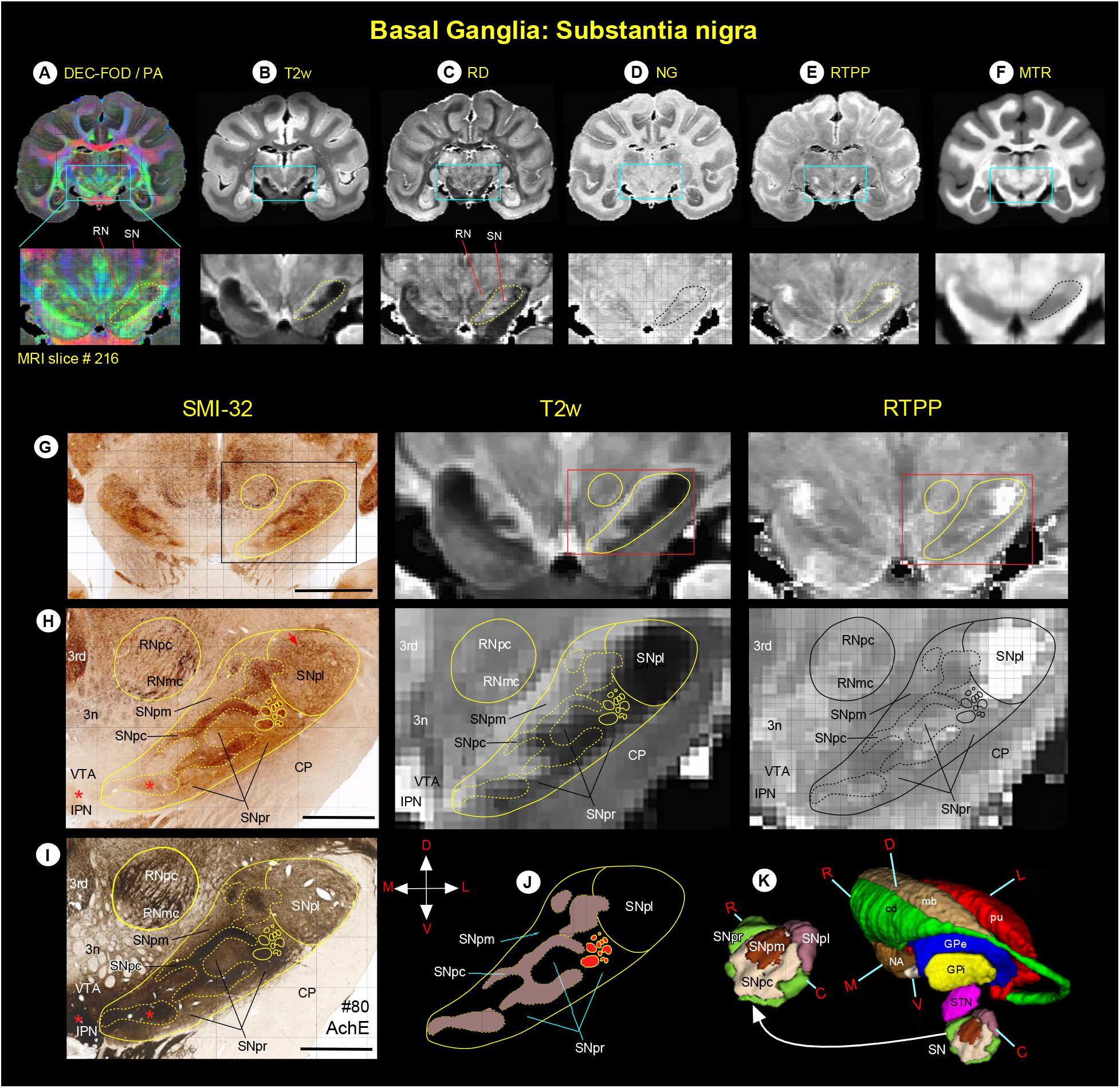
**(A-F)** Zoomed-in view of the green boxed region shows the spatial location of substantia nigra (SN), and the adjacent red nucleus (RN) in coronal MAP-MRI (DEC-FOD, *RD*, *NG*, *RTPP*), T2w, and *MTR* images. **(G-J, left and right columns)** The high-power photographs of the SMI-32 and AchE stained sections, and T2w and *RTPP* show four subregions of the SN: substantia nigra- pars compacta (SNpc), pars lateralis (SNpl), pars mixta (SNpm), and pars reticulata (SNpr). In addition, **H and I** also show the transition zone of the parvicellular and magnocellular subregions of the RN (RNpc and RNmc, respectively). The solid and dashed outlines on the MRI depicting the subregions are derived from histology section #80 on the left. **(K)** The spatial location and overall extent of the different subdivisions of the SN with other subregions of the basal ganglia in 3D, reconstructed using ITK-SNAP. For the abbreviations of basal ganglia regions, see figure legends 4 and 5. ***Orientation:*** D-dorsal; V-ventral; R- rostral; C-caudal; M-medial; L-lateral. ***Abbreviations:*** 3rd-oculomotor nucleus; 3n-oculomotor nerve; CP-cerebral peduncle; IPN-interpeduncular nucleus; VTA-ventral tegmental area. Scale bar: 5 mm (G) and 2 mm (H, I).

However, correlation with histology stains makes it possible to identify each of these unique subregions in different MR images. Thus, we first delineated the border of each subregion in SMI-32, AchE, and Nissl-stained sections and then translated this information onto MR images to identify the subregions. The SNpc is characterized by the presence of large densely packed and intensely stained cell bodies and fibers, as shown by the dashed outlined areas on the SMI- 32 and AchE stained sections (left panel in Figure 6H, I; Nissl-stained section is not shown).

This subregion has been known to contain dopaminergic cells (Arsenault et al., 1988). In contrast, the SNpr and SNpm are distinguished from SNpc by the presence of lightly-stained fiber bundles with fewer scattered cell bodies distributed among these fibers. The SNpl is located from mid-to the caudal part of SN and is also characterized by the presence of lightly stained fiber bundles, but they are more numerous and tightly packed than those of SNpr and SNpm. In addition, in SMI-32 stained sections, SNpl has moderately stained cell bodies located around its dorsolateral part that are more concentrated than in the SNpr (Fig. 6H, red arrow).

The staining patterns observed in different subregions of SN on the SMI-32 sections are comparable to those of the AchE, except the ventromedial part of the SN and the contiguous region medial to SN called the interpeduncular nucleus (IPN); both the regions showed darkly stained fibers and cell bodies in AchE stained sections (Fig. 6, see red stars in H and I). Some staining differences were also noted between the parvicellular and magnocellular subregions of the red nucleus (RNpc and RNmc, respectively) in AchE, SMI-32 (Fig. 6H, I), and other stained sections. The RNpc exhibited a more hypointense contrast than the RNmc at this mid-level of the red nucleus (a transition zone) in T2w and RTPP images. These subregions are described in detail with more illustrations in the next section (Fig. 7).

**Fig. 7.**
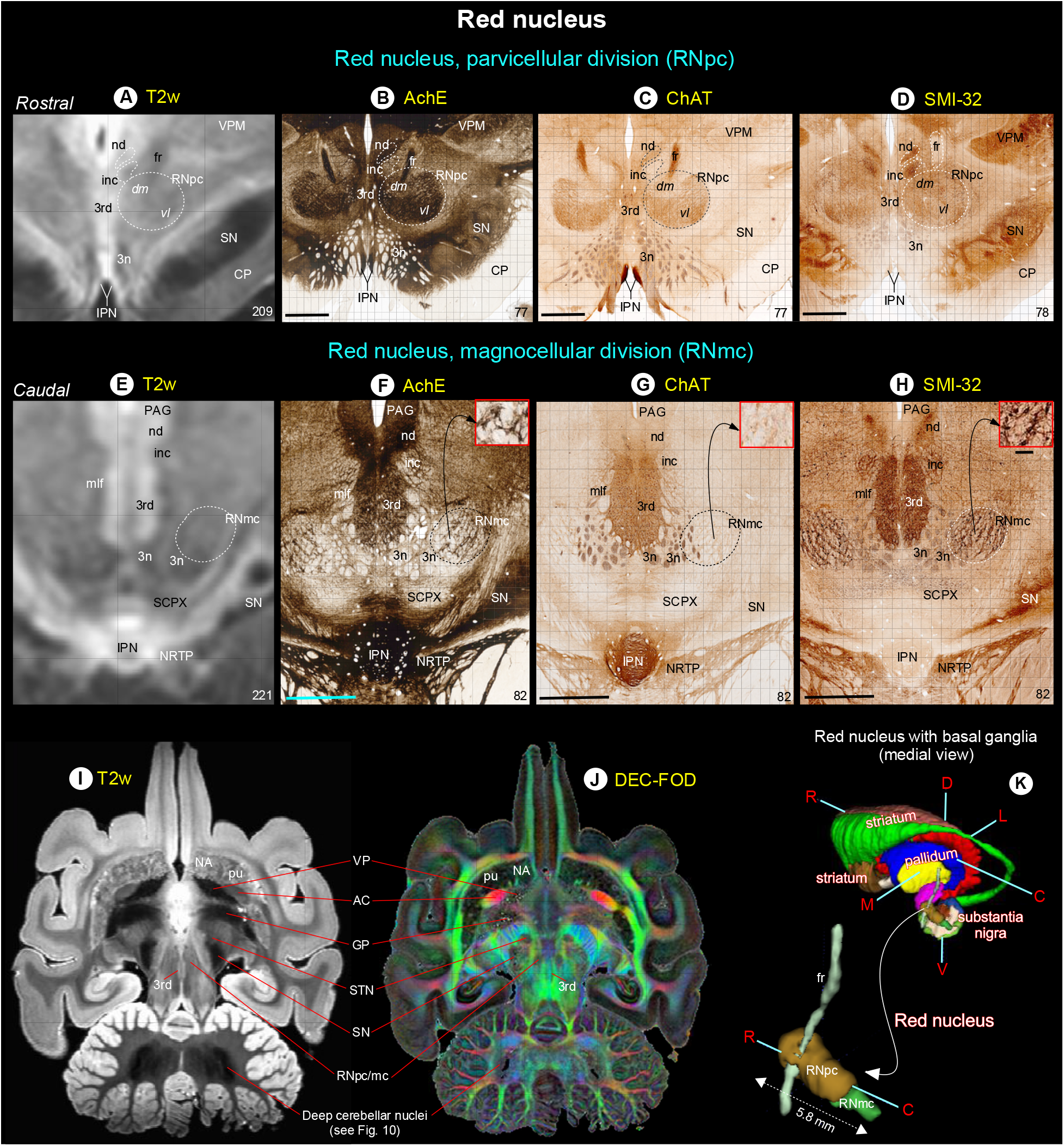
Red nucleus. **(A-H)** show the rostral parvicellular (RNpc) and the caudal magnocellular (RNmc) subregions of the red nucleus in coronal T2w, and matched histological sections. Note that the borders of RNpc and RNmc from the surrounding subcortical regions are less prominent in T2w than in AchE, ChAT and SMI-32 stained sections. The dashed outlines on the T2w images are reproduced from corresponding regions in the histology images. The numbers on the bottom right indicate the MRI slice number and histology section number. Other prominent gray and white matter regions surrounding the red nucleus are visible in these images (see abbreviations below). **(I-J)** illustrate the signal intensity differences between RN and subregions of the basal ganglia in horizontal T2w and MAP-MRI (DEC-FOD) images. In contrast to red nucleus (RNpc/mc), the globus pallidus (GP), ventral pallidum (VP), and substantia nigra (SN) exhibited significantly decreased (hypointense) signal in T2w image (see also deep cerebellar nuclei), probably due to the high level of iron content. These subregions show variable signal intensities in DEC-FOD image as shown in J. **(K)** 3D reconstruction shows the spatial location of the subregions of red nucleus with reference to basal ganglia and fr, a fiber tract which pierce through the RNpc. ***Abbreviations:*** 3^rd^-oculomotor nuclei; 3n- oculomotor nerve; AC-anterior commissure; CP-cerebral peduncle; fr-fasciculus retroflexus; inc-interstitial nucleus of Cajal; IPN-interpeduncular nucleus; mlf-medial longitudinal fasciculus; nd-nucleus of Darkschewitsch; NRTP-nucleus reticularis tegmenti pontis; PAG-periaqueductal gray; SCPX-superior cerebellar peduncle decussation; STN-subthalamic nucleus; VPM- ventral posterior medial nucleus. ***Orientation:*** D-dorsal; V-ventral; R-rostral; C-caudal; M-medial; L-lateral. Scale bars: 2 mm applies to all histology images; 0.25 mm applies to inset in H.

SN was distinguished from the surrounding areas on different MRI parameters (Fig. 6A-F, dashed outlines) but histologically defined subregions within SN were most strikingly discernible on the T2w and *RTPP* images (Fig. 6G, H, middle, and right panels). All four subregions can be identified on T2w and *RTPP* images, but the lateral subregion SNpl was the most prominent and readily identifiable as the hypointense region on T2w and hyperintense on *RTPP* images as shown at the mid-level of SN (Fig. 6H, middle, and right panels). The SNpc appears as a distinct hypointense region in T2w image, but it overlaps with the adjacent SNpr, and the border between these subregions is less distinct (Fig. 6H, see superimposed dashed lines from histology sections). In *RTPP*, the hyperintense signal is also found in the rostral part of SNpr but not in its caudal part (Fig. 6H, right panel).

Anterior and dorsolateral to the substantia nigra is the subthalamic nucleus (STN). The STN is readily identifiable as a hyper- or hypointense region on different MAP-MRI and T2w images. The signal intensity differences within the mediolateral and rostrocaudal extent of STN can be seen in MAP-MRI (for more details, see Inline Supplementary Fig. 2 and related text).

#### Red nucleus **(Fig. 7)**

The red nucleus (RN) extends for about 5.8 mm rostrocaudally and is divided into the rostral parvicellular (RNpc) region, which is contiguous with the caudal magnocellular (RNmc) region. The location and overall extent of RN with reference to the basal ganglia subregions and fiber bundle (fr) are illustrated in 3D (Fig. 7K). The RNpc is sharply demarcated from the surrounding gray and white matter structures in AchE, ChAT, and SMI-32 stained sections (Fig. 7B-C) and contains small neurons scattered within the heterogeneously stained neuropil. The staining differences of neuropil within RNpc prompted us to subdivide this region into a lightly stained dorsomedial zone (dm) and darkly stained ventrolateral zone (vl), as shown in AchE and ChAT stained sections. The demarcation between the two zones is less prominent in the SMI-32 section (Fig. 7D). The RNmc is also sharply demarcated, but it contains large intensely stained multipolar neurons with the lattice-like arrangement of their processes, identified in AchE and SMI-32 sections (Fig. 7F, H; inset). In contrast, this caudal subregion is less prominent with lightly-stained neurons and neuropil in ChAT stained section (Fig. 7G; inset). Both RNpc and RNmc are less conspicuous from the neighboring structures in T2w images (Fig. 7A, E, I), but these subregions are identified as hypointense areas in MAP-MRI (DEC-FOD; Fig. 7J). In T2w images, both subregions of RN showed heterogeneous hyperintense regions that differ significantly from the decreased (hypointense) signal observed in the substantia nigra (SN), globus pallidus (GP), and ventral pallidum (VP) (Fig. 7A, I). The other subcortical regions surrounding the RN: the oculomotor nuclei (3^rd^) and nerve (3n), the nucleus of Darkschewitsch (nd), interstitial nucleus of Cajal (inc), substantia nigra (SN), interpeduncular nucleus (IPN), and a fiber bundle-fasciculus retroflexus (fr) are delineated in both T2w and histology sections with different MR contrast and staining intensities of neuropil (Fig. 7).

### Fiber bundles associated with the basal ganglia (Fig. 8)

The fiber bundles associated with the striatum: Muratoff bundle (mb) on the dorsolateral part of the cd, anterior limb of the internal capsule (ica), and external capsule (ec) were identified on the track density (DEC-FOD) images (Fig. 4A). In addition, the pallidofugal axons that originate from different subregions of GPi form two fascicles, the ansa lenticularis (al) and lenticularis fasciculus (lf), and these fiber bundles course through Forel’s H fields (H, H1, and H2) on their way to the thalamus and brainstem (Baron et al., 2001; Lanciego et al., 2012; Neudorfer and Maarouf, 2018; Parent and Parent, 2004; Sidibe et al., 1997). The spatial locations of these fiber tracts with reference to Forel’s H field, and gray matter structures such as STN, zona inserta (zic), substantia nigra (SN), and thalamic nuclei were identified on coronal and sagittal DEC-FOD images and matched SMI-32 stained sections (Fig. 8).

**Fig. 8.**
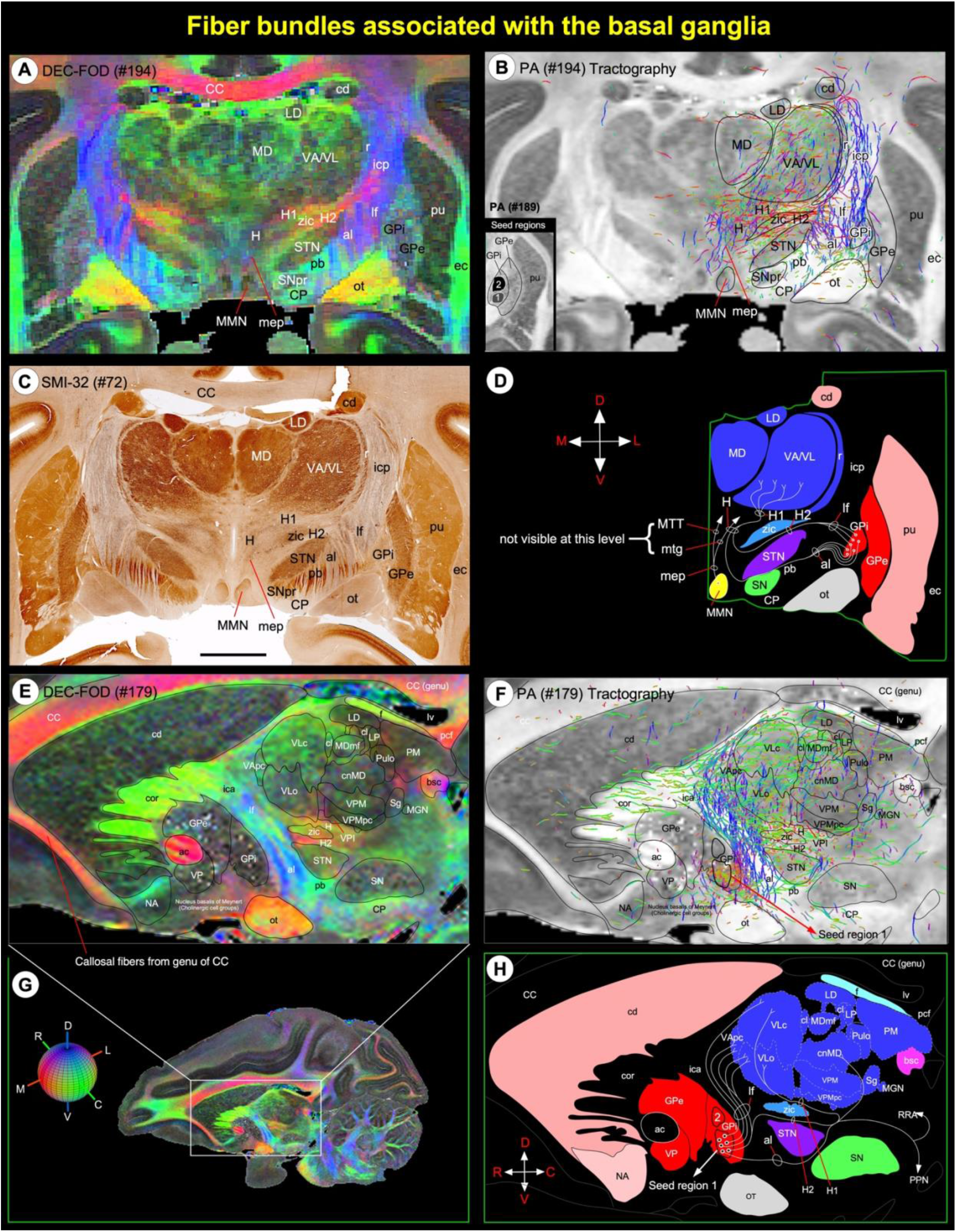
Fiber bundles linking basal ganglia and thalamus. **(A, C)** Subregions of the basal ganglia and dorsal thalamus, and associated fiber bundles on coronal DEC-FOD image, and corresponding histology section stained with SMI-32. **(E)** Zoomed-in region from sagittal DEC-FOD image in G illustrates the subregions of the basal ganglia and thalamus and associated fiber bundles. Red, green, and blue in A and E indicate the direction of the fibers (anisotropy) along mediolateral, rostrocaudal, and dorsoventral directions, respectively. For a key, see the color-coded sphere with directions at the bottom left. **(B, F)** Results of fiber tractography after placing small seed regions in the medial and lateral subregions of GPi (inset in B), as shown on the coronal and sagittal MAP-MRI (*PA*) images. Note that the direction of the fiber bundles is closely matched with the color schemes for fiber direction mapping from diffusion tensor imaging data on DEC-FOD images. **(D, H)** Schematic diagrams illustrate the projections from the GPi (internal segment of the globus pallidus) to different nuclei in the thalamus through ansa lenticularis (al), lenticular fasciculus (lf), and H, H1, and H2 fields of Forel. The direction of the fibers and projection targets is based on previous anatomical tracing studies in the macaque monkey. ***Abbreviations:*** ac-anterior commissure; al-ansa lenticularis; bsc-brachium of superior colliculus; CC-corpus callosum; cd-caudate nucleus; cor-corona radiata; cl-central lateral nucleus; cnMD-centromedian nucleus; CP-cerebral peduncle; ec-external capsule; f-fornix; GPe- globus pallidus external segment; GPi- globus pallidus internal segment; H, H1, H2-Fields of Forel; ica-internal capsule, anterior limb; icp-internal capsule, posterior limb; LD-lateral dorsal nucleus; lf-lenticular fasciculus; LP-lateral posterior nucleus; lv-lateral ventricle; MD- medial dorsal nucleus of thalamus; MGN-medial geniculate nucleus; MDmf-mediodorsal thalamus, multiform division; mep- mammillary efferent pathway; MMN-medial mammillary nucleus; mtg-mammillotegmental tract; MTT-mammillothalamic tract; NA-nucleus accumbens; ot-optic tract; pb-pontine bundle; pcf-posterior column of fornix; PM-medial pulvinar; pu-putamen; PPN-pedunculopontine tegmental nucleus; Pulo-pulvinar oralis nucleus; r-reticular nucleus; RRA-retrorubral area; Sg- suprageniculate nucleus; SN-substantia nigra; SNpr-substantia nigra pars reticulata; STN-subthalamic nucleus; VA-ventral anterior nucleus; VApc-ventral anterior nucleus, parvicellular division; VL-ventral lateral; VLc-ventral lateral caudal nucleus; VLo- ventral lateral oral nucleus; VP-ventral pallidum; VPI-ventral posterior inferior nucleus; VPM-ventral posterior medial nucleus; VPMpc-ventral posterior medial nucleus, parvicellular division; zic-zona incerta. ***Orientation:*** D-dorsal; V-ventral; R-rostral; C-caudal; M-medial; L-lateral. Scale bar: 5 mm (C).

The projections that originate from neurons located in the lateral or medial subregions of GPi (Parent et al., 2001) form the ansa lenticularis (al) that courses ventromedially around the posterior limb of the internal capsule (icp) and then bifurcates posteriorly to enter Forel’s H field (also known as the prerubral field; Fig. 8C-D). Similarly, axons from both subregions of GPi neurons course through the internal capsule to form the lenticular fasciculus (lf), located between STN and zona inserta (also called Forel’s field H2; Fig. 8D). Ultimately, the lenticular fasciculus merges with the ansa lenticularis at the level of field H of Forel to enter the thalamic nuclei through the thalamic fasciculus (H1 field of Forel) (Fig. 8D, H). The main thalamic targets of pallidum are the ventral anterior (VA), subregions of ventral lateral (VL, VLc, VLo), and the centromedian (cnMD) and intralaminar thalamic nuclei (Fig. 8A, C, E). These and other thalamic nuclei are described in detail in the next section below.

We also confirmed the trajectory of the different fiber bundles (see above) using fiber tractography by placing small seed regions in the medial and lateral subregions of GPi, as shown in both coronal and sagittal sections of PA images (Fig. 8B, F). The direction of the fiber bundles can be matched with the color schemes for fiber direction mapping from diffusion tensor imaging (DTI) data on DEC-FOD images (Fig. 8A, E, G).

#### Thalamus (Fig. 9)

We adapted the terminology of thalamic nuclei similar to that of (Olszewski, 1952). The thalamus is divided into the dorsal thalamus, epi-thalamus, and geniculate region (Jones, 1998) (Fig. 9). The dorsal thalamus is further divided into anterior, medial, lateral, intralaminar, and posterior groups and has significant roles in motor, emotion, memory, arousal, and other sensorimotor functions (Halassa and Kastner, 2017; Mitchell et al., 2014; Pergola et al., 2018). The epithalamus is located in the posterior dorsal part of the diencephalon, and its principal gray matter structure is habenular nuclei, which play a pivotal role in reward processing, aversion, and motivation (Hikosaka et al., 2008; Roman et al., 2020). The geniculate region includes medial and lateral geniculate bodies, and it serves as an important relay nucleus in the auditory and visual pathways, respectively. These multiple subregions of the thalamus with different architectonic characteristics (Jones, 1998) prompted us to use a combination of MRI parameters and matched AchE, SMI-32, and other immunostained sections to distinguish different thalamic nuclei (Fig. 9; see Inline Supplementary Fig. 3).

**Fig. 9.**
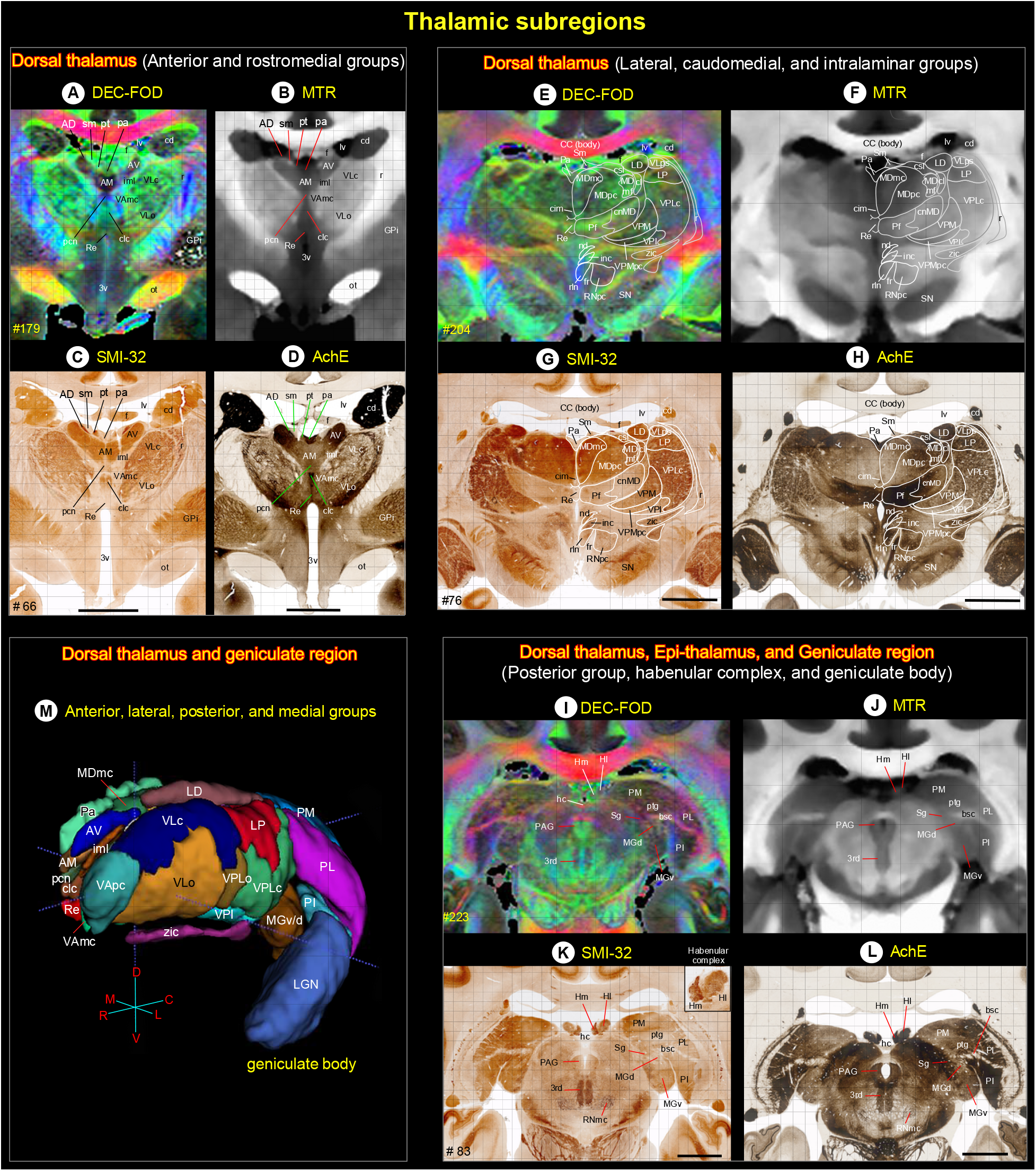
Thalamus. **(A-L)** The MR signal intensity differences between dorsal thalamus (anterior, medial, intralaminar, and posterior groups), epithalamus (habenular complex), and geniculate region (geniculate body) in *PA*/DEC-FOD and MTR images, and the corresponding subregions in the histological sections stained with SMI-32, and AchE. See also inline Suppl Fig. 3 for these thalamic subregions in other MRI parameters and histological sections. **(K, inset)** Heterogenous staining of neuropil within the medial (Hm) and lateral (Hl) subregions of the habenular complex in SMI-32 stained section. **(M)** The spatial location and overall extent of the anterior, lateral, posterior, and some medial groups of thalamic nuclei in 3D, reconstructed using ITK-SNAP. ***Abbreviations:*** 3^rd^-third cranial (oculomotor) nuclei; 3v-3^rd^ ventricle; bsc-brachium of superior colliculus; AD-anterior dorsal nucleus; AM-anterior medial nucleus; AV-anterior ventral nucleus; CC-corpus callosum; cd-caudate nucleus; cim-central intermediate nucleus; cl-central lateral nucleus; clc-central latocellular nucleus; cnMD-centromedian nucleus; csl-central superior lateral nucleus; f-fornix; fr-fasciculus retroflexus; GPi-globus pallidus, internal segment; hc-habenular commissure; Hl-lateral habenular nucleus; Hm-medial habenular nucleus; iml-internal medullary lamina; inc-interstitial nucleus of Cajal; LD-lateral dorsal nucleus; LGN-lateral geniculate nucleus; LP-lateral posterior nucleus; lv-lateral ventricle; MDmc-medial dorsal nucleus, magnocellular division; MDmf-medial dorsal nucleus, multiform division; MDpc-medial dorsal nucleus, parvicellular division; MGd-medial geniculate nucleus, dorsal division; MGv-medial geniculate nucleus, ventral division; nd-nucleus of Darkschewitsch; ot-optic tract; Pa-paraventricular nucleus; PAG-periaqueductal gray; pcn-paracentral nucleus; Pf-parafascicular nucleus; PI-inferior pulvinar; PL-lateral pulvinar; PM-medial pulvinar; pt-parataenial nucleus; ptg-posterior thalamic group; r- reticular nucleus; Re-reunions nucleus; rln-rostral linear nucleus; RNmc-red nucleus, magnocellular division; RNpc-red nucleus, parvicellular division; Sg-suprageniculate nucleus; Sm-stria medullaris; SN-substantia nigra; VAmc-ventral anterior nucleus, magnocellular division; VApc-ventral anterior nucleus, parvicellular division; VLc-ventral lateral caudal nucleus; VLo-ventral lateral oral nucleus; VLps-ventral lateral postrema nucleus; VPI-ventral posterior inferior nucleus; VPLc-ventral posterior lateral caudal nucleus; VPLo-ventral posterior lateral oral nucleus; VPM-ventral posterior medial nucleus; VPMpc-ventral posterior medial nucleus, parvicellular division; zic-zona incerta. Scale bars: 5 mm applies to C-D, G-H, and K-L.

### Dorsal Thalamus

#### Anterior and rostromedial groups

The anterior group of nuclei has a distinct pattern of connections with the hippocampal formation and other cortical areas, and it is thought to be involved in certain categories of learning and memory and spatial navigation (Aggleton et al., 2010; Jankowski et al., 2013; Shah et al., 2012). It is located in the dorsal part of the rostral thalamus and divided into the anterior dorsal (AD), anterior medial (AM), and anterior ventral (AV) nuclei. The AM, AV, and AD together form a distinct “V”-shaped band, visible as hypo- or hyperintense clusters on DEC-FOD, MTR, RTAP, and RD (Fig. 9A, B; see Inline Supplementary Fig. 3A, B). This band is separated from the lateral group of thalamic nuclei (e.g., VAmc/VLc) ventrolaterally by a distinct hypo- or hyperintense fiber bundle called the intermedullary lamina (iml). It is a continuation of the mammillothalamic tract (MTT) that originates from the mammillary bodies. Caudal to the AV is the prominent lateral dorsal (LD) nucleus, which shows hypointense contrast similar to that of the AV in DEC-FOD (Fig. 9A, E, M).

The anterior group of nuclei is bordered dorsally by the medial group (paraventricular-Pa, parataenial-pt) and ventrally by the intralaminar group of nuclei (see below). Pa is located along the midline, adjacent to AM rostrally but adjacent to mediodorsal thalamic nuclei (MD) caudally (Fig. 9E, F). The pt is a small strip of gray matter with a limited rostrocaudal extent and is difficult to distinguish from the neighboring gray or white matter. It shows variable signal intensities in different MRI parameters. The anterior group is also bordered dorsally by two distinct hyperintense fiber bundles, the stria medullaris (Sm) and fornix (f), which are easily delineated from the anterior group on the MTR image (Fig. 9B). The spatial location of these nuclei in MRI is confirmed in adjacent stained histological sections. Within anterior and rostromedial groups, the AV, AD, and Pa are more prominent and intensely stained in AchE than in SMI-32, ChAT, and PV stained sections (Fig. 9C, D; see Inline Supplementary Fig. 3C, D).

#### Lateral, caudomedial, and intralaminar groups

The lateral group is a large division of the thalamus with 11 subnuclei and extended for about 9 mm in the rostrocaudal direction. It includes the ventral anterior (VApc, VAmc), ventral lateral (VLc, VLo, VLps), ventral posterior (VPLc, VPLo, VPM, VPMpc, VPI), and lateral posterior (LP) nuclei. The overall extent of these subregions is visualized in 3D segmentation and coronal sections at different rostrocaudal levels (Fig. 9A, E, M). Anatomical tracing studies in non-human primates show that these nuclei represent the principal thalamic relays for inputs from the substantia nigra pars reticulata (SNpr), internal segment of the globus pallidus (GPi), deep cerebellar nuclei, and/or medial/trigeminal lemniscus (Asanuma et al., 1983; Kaas, 2012; Rouiller et al., 1994; Sakai et al., 1996).

The VAmc (ventral anterior, magnocellular division), VLo (ventral lateral oral), and VLc (ventral lateral caudal nucleus) are located in the rostral part of the lateral group. Both VAmc and VLo exhibited hypo- or hyperintense contrast, but the opposite of VLc in different MRI parameters, and this distinction was prominent in MTR, RTAP, and RD images (Fig. 9B; see Inline Supplementary Fig. 3A, B). The VLc is continuous caudally with the LP, which shows a more hypointense signal than the VLc in both DEC-FOD and MTR. Similarly, the rostrally located VLo exhibited a more hyperintense contrast than the caudally located VPLc in both MR images (Fig. 9, top row). The staining intensity of neuropil is also different in these regions in SMI-32 and AchE stained sections (Fig. 9, middle row).

The caudal portion of the lateral group is mainly dominated by the ventral posterior and lateral posterior nuclei. Among this group, the VPM (ventral posterior medial), VPMpc (ventral posterior medial, parvicellular division), and LP (lateral posterior) are the most prominent and exhibited similar hyper- or hypointense contrast in different MRI parameters (Fig. 9E, F; see Inline Supplementary Fig. 3E, F). This restricted microarchitectural feature is easily distinguished from the adjacent nuclei: VPI (ventral posterior inferior), VPLc (ventral posterior lateral caudal), and VLps (ventral lateral postrema nucleus), with opposite signal intensity.

The prominent caudomedial group includes the mediodorsal thalamus with three subregions (MDmc, MDpc, and MDmf), and it is involved in many cognitive tasks such as decision making, working memory, and cognitive control related to the prefrontal cortex (Ouhaz et al., 2018). Each of these subregions exhibited different signal intensities in DEC-FOD, MTR, and other MRI parameters, which coincided with the neuropil staining in SMI-32, AchE, and other stained sections (Fig. 9G, H; see Inline Supplementary Fig. 3G, H). For example, MDmc (mediodorsal magnocellular) exhibited a hypointense signal compared to MDpc (mediodorsal parvicellular) in DEC-FOD, and this feature coincided with the dark- and lightly stained neuropil in these subregions, respectively in SMI-32 stained section (Fig. 9E, G).

The intralaminar groups with 10 nuclei (pcn, Clc, cdc, cim, cif, cl, Re, csl, cnMD, and Pf) are distributed rostrocaudally along the midline of the thalamus (Fig. 9). Due to the small size (except Pf and cnMD), the spatial location of these nuclei on MRI was difficult to distinguish from neighboring structures. These midline nuclei were marked on the MR images based on the spatial location and staining patterns observed in histologically stained sections. The Pf (parafascicular) and cnMD (centromedian) nuclei are visible as a mediolaterally oriented band of hypointense to hyperintense regions, respectively (or vice versa) in different MRI parameters (Fig. 9E, F; see Inline Supplementary Fig. 3E, F). The distinction between Pf and cnMD is less prominent in MTR, but the cnMD is more hypo- or hyperintense than Pf in DEC-FOD, RTAP, and RD as shown in figure 9E, F, and Inline Supplementary figure 3E, F. The staining differences between these two nuclei are more prominent in AchE and ChAT than in SMI-32 and PV sections (Fig. 9G, H; see Inline Supplementary Fig. 3G, H). Among the lateral group of nuclei, Pf showed the darkly stained neuropil in AchE (Fig. 9H).

#### Posterior group

The posterior group is dominated by medial, lateral, and inferior pulvinar nuclei (PM, PL, and PI, respectively), and has strong connections with the visual cortical areas in the dorsal and ventral stream. These three subregions of the pulvinar are demarcated by a prominent fiber bundle, the brachium of the superior colliculus (bsc), which divides PM and PL from the PI and the medial geniculate body (MGB, part of the geniculate region, see below) (Fig. 9I, J). The PL and PI exhibited greater hypointense contrast relative to the PM in DEC-FOD, and this feature corresponded well with the dark neuropil staining in these subregions in AchE (Fig. 9I, L).

#### Epi-thalamus

The medial and lateral subregions of the habenular nuclei (Hm and Hl, respectively) and the associated fiber bundle, the habenular commissure (hc) form the principal gray and white matter structures of the epithalamus. The Hl is more hypointense than Hm in DEC-FOD, but both subregions are hyperintense in the MTR image (Fig. 9I, J). The neuropil within the Hm is more intensely stained than in Hl in SMI-32, but the staining differences between these two subregions are less prominent in AchE stained section (Fig. 9K, L). In addition, both Hm and Hl show heterogeneous compartments of variable-staining intensities in SMI-32 (see inset in Fig. 9K).

#### Geniculate region

It is composed of the lateral geniculate nucleus and the medial geniculate body (LGN and MGB, respectively). The contour of the LGN is easily distinguished from the surrounding regions in MR images, but the signal intensity differences between different laminae within LGN are less distinct (Fig. 9G, H). The MGB is divided into different subregions (e.g., MGd and MGv) and can be distinguished based on different signal intensities in different MRI parameters and staining patterns in histological sections (Fig. 9I-L).

### Hypothalamus (Inline Supple Fig. 4)

The hypothalamus can be divided rostrocaudally into the rostral, middle (tuberal), and caudal groups (Rempel-Clower and Barbas, 1998; Wells et al., 2020) and has strong connections with the orbital and medial prefrontal cortex (Ongur et al., 1998). Two of the MAP-MRI parameters (RD and AD) and matched AchE and SMI-32 stained sections are useful in delineating different subnuclei in the hypothalamus. For other details, see Inline Supplementary Fig. 4 and related text.

### Brainstem (Figs. 10-11)

The sagittal sections of MAP-MRI and T2w are very useful in delineating gray and white matter subregions in the different rostrocaudal extent of the brainstem (midbrain, pons, and medulla) and cerebellum. We delineated several sensory and motor nuclei, and fiber tracts of different sizes and orientations linking cerebral cortex and diencephalon with the spinal cord in the midbrain, pons, and medulla on three sagittal DEC-FOD images spaced 0.8mm, 2.2mm, and 4.2mm from the midline (Fig. 10A-C). The orientation of the fiber bundles was confirmed with fODFs (fiber orientation distribution function) as illustrated on the directionally encoded color images in A-C. We also confirmed the spatial location of identified structures on selected coronal DEC-FOD and *RTPP* images from the midbrain, pons, and medulla, aided by AchE histology staining (Fig. 11). The contours of brainstem nuclei and fiber bundles on sagittal or coronal sections were derived from the serially segmented 3D dataset (SC21-see *“Segmentation of subcortical regions”* in materials and method section). The architectonic regions of the brainstem were identified with reference to the photographic atlas of the human brain (DeArmond et al., 1989) and Duvernoy’s atlas of the human brainstem and cerebellum (Naidich et al., 2009). We also used another atlas (Paxinos et al., 2009) to identify few selected nuclei in the brainstem. The spatial location of some brainstem nuclei and fiber bundles in our macaque MRI is comparable to humans.

**Fig. 10.**
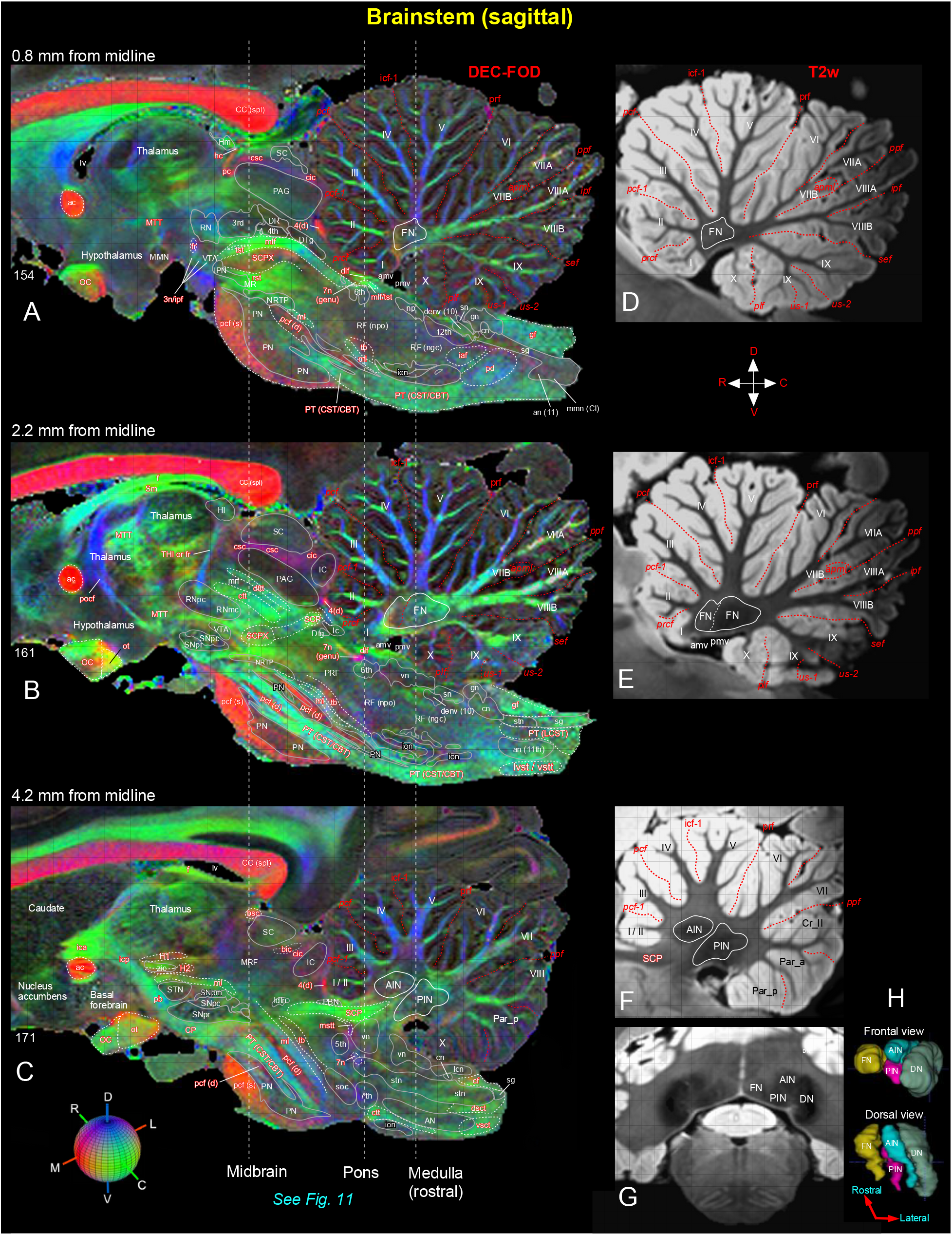
Brainstem and cerebellum. **[A-C]** Mediolateral extent of the brainstem (midbrain, pons, and medulla) with the cerebellum in three sagittal DEC-FOD images, spaced 0.8 mm, 2.2 mm, and 4.2 mm from the midline. ***Brainstem***: white letters with solid white outlined regions indicate the gray matter regions or nuclei, and the reddish white letters with or without white dashed lines illustrate the major fiber tracts running in different directions in the brainstem (see the color-coded sphere with directions at the bottom left. Red, green, and blue in these images indicate the anisotropy along mediolateral (ML), rostrocaudal (RC), and dorsoventral (DV) directions, respectively. Three dashed lines passing through the sagittal sections indicate the coronal sections at the level of the midbrain, pons, and medulla as shown in Figure 11. Other gray and white matter subregions rostral to the brainstem are also included. ***Cerebellum***: white or black letters indicate different cerebellar lobules (I-X), and red dashed lines show the sulci separating different lobules. **(D-F)** The corresponding location of the cerebellum with different lobules and sulci in three sagittal T2w images. **(G)** Coronal T2w slice at the level of the rostral medulla corresponds to the right dashed line on the left panel shows deep cerebellar nuclei (also check Figure 11G-I). **(H)** The 3D reconstruction shows the spatial location of four deep cerebellar nuclei in different views. ***Abbreviations for Brainstem and surrounding regions [A-C]:*** 3^rd^-oculomotor nuclei; 3n-oculomotor nerve; 4^th^-trochlear nuclei; 4(d)-trochlear nerve decussation; 5^th^-trigeminal nuclei; 6^th^-abducent nuclei; 7^th^- facial nuclei; 7n-facial nerve; 7n (genu)-genu of the facial nerve; 12^th^-hypoglossal nucleus; ac-anterior commissure; AN- ambiguous nucleus; an (11)-accessory nucleus; bic-brachium of inferior colliculus; bsc-brachium of the superior colliculus; CBT- corticobulbar tract; CC (spl)-splenium of corpus callosum; cf-cuneate fasciculus; cic-commissure of inferior colliculus; cn- cuneate nucleus; CP-cerebral peduncle; csc-commissure of superior colliculus; CST-corticospinal tract; ctt-central tegmental tract; denv (10)-dorsal efferent nucleus of vagus; dlf-dorsal longitudinal fasciculus; DR-dorsal raphe; dsct-dorsal spinocerebellar tract; DTg-dorsal tegmentum; dttt-dorsal trigemino thalamic tract; f-fornix; fr-fasciculus retroflexus; gf-gracile fasciculus; gn- gracile nucleus; H1-H1 field of Forel; H2-H2 field of Forel; hc-habenular commissure; Hl-lateral habenular nucleus; Hm-medial habenular nucleus; iaf-internal arcuate fibers (sensory decussation); IC-inferior colliculus; ica-anterior limb of the internal capsule; icp-posterior limb of the internal capsule; ion-inferior olivary nucleus; ipf-inter peduncular fossa; IPN-interpeduncular nucleus; lc-locus coeruleus; lcn-lateral cuneate nucleus; ldtn-lateral dorsal tegmental nucleus; LCST-lateral corticospinal tract; lv-lateral ventricle; lvst-lateral vestibulospinal tract; ml-medial lemniscus; mlf-medial longitudinal fasciculus; MMN-medial mammillary nucleus; mmn (CI)-medial motor nucleus, cervical level; MR-median raphe; mrf-mesencephalic reticular formation; MRF-midbrain reticular formation; mstt:mesencephalic trigeminal tract; MTT-mammillothalamic tract; np-nucleus proprius; NRTP-nucleus reticularis tegmenti pontis; OC-optic chiasm; ot-optic tract; PAG-periaqueductal gray; pb-pontine bundle; PBN- parabrachial nucleus; pc-posterior commissure; pcf (d)-pontocerebellar fiber, deeper part; pcf (s)-pontocerebellar fibers, superficial part; pd-pyramidal decussation; PN-pontine nuclei; pocf- postcommissural fornix; PRF-pontine reticular formation; PT-pyramidal tract; RF (ngc)-reticular formation, nucleus gigantocellularis; RF (npo)-reticular formation, nucleus pontis centralis oralis; RNmc-red nucleus, magnocellular division; RNpc-red nucleus, parvicelluar division; rst-rubrospinal tract; SC-superior colliculus; SCP-superior cerebellar peduncle; SCPX-superior cerebellar peduncle decussation; sg-substantia gelatinosa; Sm-stria medullaris; sn-solitary nucleus; SNpc-substantia nigra pars compacta; SNpm-substantia nigra pars mixta; SNpr-substantia nigra pars reticulata; soc-superior olivary complex; stn-spinal trigeminal nucleus; STN-subthalamic nucleus; tb-trapezoid body; THI- habenular interpeduncular tract; tst-tectospinal tract; vn-vestibular nuclei; vsct-ventral spinocerebellar tract; vstt-ventral spinotectal tract; VTA-ventral tegmental area; zic-zona incerta. ***Abbreviations for cerebellar regions (A-H):*** I to X-cerebellar lobules; AIN-anterior interposed nucleus; amv-anterior medullary velum; Cr_II-crus II of ansiform lobule; DN-dentate nucleus; FN-fastigial nucleus; Par_a-paramedian lobule anterior part; Par_p-paramedian lobule posterior part; PIN-posterior interposed nucleus; pmv-posterior medullary velum. ***Cerebellar sulci/fissures (A-F):*** apml-ansoparamedian lobule; icf1-intraculminate fissure 1; ipf-intrapyramidal fissure; pcf-preculminate fissure; pcf1-preculminate fissure 1; plf-posterolateral fissure; ppf- prepyramidal fissure; prcf-precentral fissure; prf-primary fissure; sef-secunda fissure; us1-uvular sulcus 1; us2-uvular sulcus 2.

**Fig. 11.**
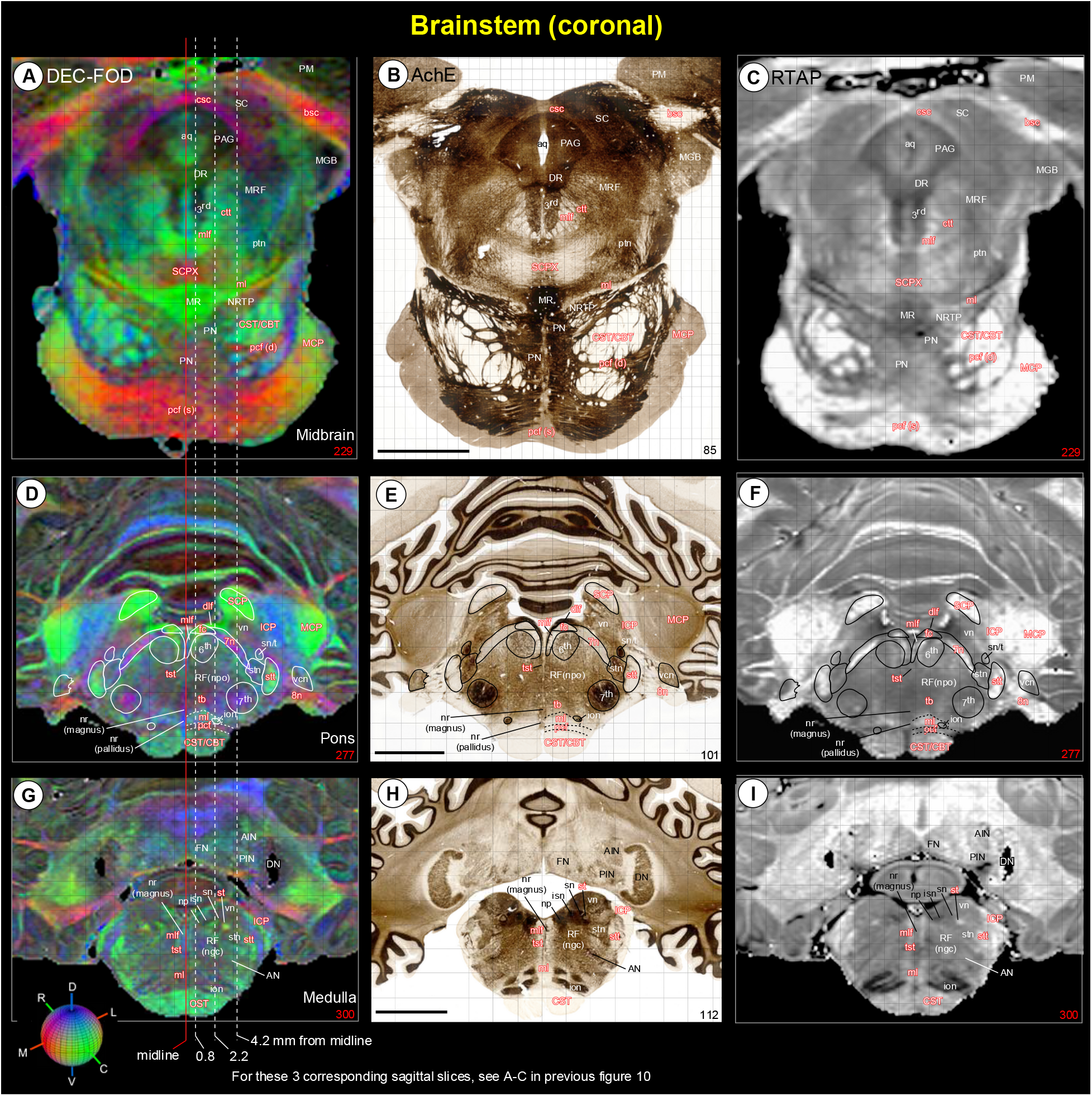
Brainstem. The spatial location of different nuclei and fiber tracts at the level of the midbrain **[A-C]**, pons **(D-F)**, and rostral medulla **(G-I)** in coronal MAP-MRI (DEC-FOD and *RTAP*) images and histology sections stained with AchE. The white or black letters indicate the gray matter regions or nuclei, and the reddish white letters illustrate the major fiber tracts running in different directions in the brainstem (see the color-coded sphere with directions at the bottom left; DV-dorsoventral; RC- rostrocaudal; ML-mediolateral). Three dashed lines passing through the coronal sections indicate the sagittal sections, spaced 0.8 mm, 2.2 mm, and 4.2 mm from the midline as illustrated in Figure 10. Note that many of the identified gray and white matter regions in high-resolution MAP-MRI closely matched with the adjacent histology section (compare left and right panels with the middle panel). ***Abbreviations:*** 3^rd^-oculomotor nuclei; 6^th^-abducent nuclei; 7^th^-facial nuclei; 7n-facial nerve; 8n-vestibulocochlear nerve; AIN-anterior interposed nucleus; AN-ambiguous nucleus; aq-aqueduct; bsc-brachium of the superior colliculus; CBT- corticobulbar tract; csc-commissure of superior colliculus; CST-corticospinal tract; ctt-central tegmental tract; dlf-dorsal longitudinal fasciculus; DN-dentate nucleus; DR-dorsal raphe; fc-facial colliculus; FN-fastigial nucleus; ICP-inferior cerebellar peduncle; ion-inferior olivary nucleus; isn-inferior salivatory nucleus; MGB-medial geniculate body; ml-medial lemniscus; mlf- medial longitudinal fasciculus; MR-median raphe; MRF-midbrain reticular formation; np-nucleus proprius; nr (magnus)-nucleus raphe magnus; nr (pallidus)-nucleus raphe pallidus; NRTP-nucleus reticularis tegmenti pontis; PAG-periaqueductal gray; pcf- pontocerebellar fibers; pcf (d)-pontocerebellar fiber, deeper part; pcf (s)-pontocerebellar fibers, superficial part; PIN-posterior interposed nucleus; PM-medial pulvinar; PN-pontine nuclei; ptn-pedunculotegmental nucleus; RF (ngc)-reticular formation, nucleus gigantocellularis; RF (npo)-reticular formation, nucleus pontis centralis oralis; SC-superior colliculus; SCP-superior cerebellar peduncle; SCPX-superior cerebellar peduncle decussation; sn-solitary nucleus; sn/t-solitary nucleus/tract; st-solitary tract; stn-spinal trigeminal nucleus; stt-spinal trigeminal tract; tb-trapezoid body; tst-tectospinal tract; vcn-ventral cochlear nucleus; vn-vestibular nuclei. Scale bar: 5 mm (B, E, H).

#### Midbrain

At the midbrain level, three well-defined structures with low diffusion anisotropy are identified on the DEC-FOD image, 0.8 mm lateral to the midline. These structures are the periaqueductal gray [PAG], and oculomotor [3^rd^], and trochlear (4^th^) cranial nerve nuclei (Figs. 10A, 11A, the 4^th^ nucleus is not visible at this coronal level). The PAG is located ventromedial to the superior and inferior colliculus (SC and IC), and crossing fibers associated with the colliculus, the commissures of the superior and inferior colliculi (csc and cic, respectively). The oculomotor and trochlear nerve nuclei are situated ventral to the dorsal raphe (DR), and dorsal to the prominent rostrocaudally oriented medial longitudinal fasciculus (mlf) tract, and mediolaterally oriented superior cerebellar peduncle decussation (SCPX) fibers (Fig. 10A). The midbrain reticular formation (MRF), a large diffuse region with indistinct border and variable signal intensities occupies the lateral part of the midbrain, as shown on the DEC-FOD images (Fig. 10C, 11A). The MRF has lattice-like neuropil staining in AchE stained section (Fig. 11B). The other major gray matter structure, spanning the entire rostrocaudal extent of the midbrain is the pontine nuclei (PN). The PN is most strikingly delineated from rostrocaudally oriented corticospinal/corticobulbar tracts (CST/CBT or pyramidal tract-PT), and mediolaterally oriented superficial and deeper parts of the pontocerebellar fibers (pcf-s and pcf-d, respectively), as revealed by fODFs with DEC map (Figs. 10A-C; 11A). These subregions are less distinct and spatially not distinguishable from each other on a conventional MRI, or a population-averaged anatomical MRI volume (see Inline Supplementary Fig. 6). Two other gray matter regions with moderate-to-low anisotropy: median raphe (MR) and nucleus reticularis tegmenti pontis (NRTP) are distinguished dorsal to PN and ventral to SCPX in DEC-FOD and RTAP images (Fig. 11A, C). Both MR and NRTP are well-defined regions with darkly stained neuropil in AchE (Fig. 11B).

The other structures located rostral to the midbrain in the diencephalon region with variable signal intensities are the red nucleus (RN), different subregions of the substantia nigra (SN), subthalamic nucleus (STN), zona incerta (zic), and associated fiber bundles H1 and H2 (Fig. 10A-C), which are described in detail in the basal ganglia section (see Figs. 6, 8). Ventral to RN and dorsomedial to SN is the ventral tegmental area (VTA), which exhibited low anisotropy in the DEC-FOD image (Fig. 10B). The other prominent fiber tracts with different orientations: the stria medullaris (Sm), habenular interpeduncular tract or the fasciculus retroflexus (THI/fr), postcommissural fornix (pocf), and the mammillothalamic tract (MTT) are visible on the sagittal slice spaced 2.2 mm from the midline (Fig. 10B). Sm innervates the habenular nucleus and THI/fr forms the output of the habenular nucleus and projects to the midbrain and hindbrain monoaminergic nuclei (Roman et al., 2020). The pocf originates from the hippocampus and projects to the mammillary bodies, which in turn terminates on the anterior thalamic nuclei through MTT (Aggleton et al., 2010).

#### Pons

Six major cranial nerve nuclei with low to moderate diffusion anisotropy, the trigeminal (5^th^), abducens (6^th^), facial (7^th^), vestibular (vn), spinal trigeminal [stn] nuclei, and superior olivary complex (soc) are distinguished in the different rostrocaudal and mediolateral extent of the pons on DEC-FOD images (Fig. 10A-C). These nuclei, except the 6^th^, are visible ventral to the superior cerebellar peduncle (SCP) and caudal to the medial lemniscus (ml), and auditory-related trapezoid body (tb) on the DEC-FOD image spaced 4.2 mm from the midline (Fig. 10C). The 6^th^ nucleus is located close to the midline and immediately ventral to the crossing fibers of the 7^th^ nerve, which forms the genu of the facial nerve or the facial colliculus-fc (Figs. 10B, 11D-F).

The anterior and ventral to the 6^th^ nucleus is the pontine reticular formation (PRF) and the nucleus pontis centralis oralis (RF-npo), and these are diffuse regions with less distinct borders (Fig. 10B). Among these identified cranial nerve nuclei in the pons, the 7^th^ (facial) is the most intensely stained region in the AchE section (Fig. 11E).

#### Medulla

The rostroventral part of the medulla is dominated by the well-defined hypointense inferior olivary nucleus [ion], which is strikingly delineated from ventrally adjacent pyramidal tract (CST/CBT) and dorsally located reticular formation (RF-ngc) in DEC-FOD and *RTPP* images (Figs. 10A, B; 11G, I). AchE stained section also revealed a clear distinction between these subregions comparable to the *RTPP* image (Fig. 11F, I). In addition, several nuclei are delineated at the different mediolateral and rostrocaudal extent of the medulla: nucleus prepositus (np), the hypoglossal nucleus (12^th^), the dorsal motor nucleus of the vagus (denv-10), solitary nucleus (sn), gracile nucleus (gn), cuneate nucleus (cn), vestibular nucleus (vn), and spinal trigeminal nucleus (stn). The 12^th^ nucleus and vn exhibited lower anisotropy than other nuclei (Fig. 10A, B). The crossing fibers of the sensory and motor pathways: the internal arcuate fibers (iaf), and pyramidal decussation (pd) dominate the caudal part of the medulla and exhibited higher anisotropy than the surrounding nuclei (Fig. 10A). The accessory nucleus (11^th^), medial motor neurons (mmn-C1), and the substantia gelatinosa (sg), which are located at the medulla and cervical spinal cord junction also exhibited lower anisotropy than the dorsally located gracile fasciculus (gf) sensory tract (Fig. 10A). The ambiguous nucleus (AN) is visible as a rostrocaudally extended hypointense region on the lateral part of the medulla and can be demarcated from the surrounding hyperintense dorsal and ventral spinocerebellar tracts (dsct and vsct), and the central tegmental tract (ctt) (Fig. 10C).

### Cerebellum

Although unrelated to the present study, the unique foliation and compartmental organization of the cerebellar cortex, prompted us to identify and segment this non-subcortical region in our MAP-MRI parameters and include it with the subcortical atlas SC21. We identified the spatial location of 10 lobules (I-X), paramedian lobules (Par), simple lobule (Sim), ansiform lobules Crus I and II, flocculus (Fl), and paraflocculus (PFl) with reference to 11 landmarks (nine fissures and two sulci) through the rostrocaudal and mediolateral extent of the cerebellum on the sagittal MR images (Fig. 10A-C). The nomenclature and abbreviations used to identify these lobules and landmarks are according to (Larsell, 1953), but with some modifications to include the simplified version of the abbreviations for the fissures and sulci (see Fig. 10 legend). The lobules of the cerebellar cortex and the orientation of associated fiber bundles exhibited similar signal intensities in the different mediolateral extent of the cerebellum on DEC-FOD (Fig. 10A-C) and other MRI parameters.

The deep cerebellar nuclei (DCN), the sole output of the cerebellum, are key structures of the cortico-cerebellar circuitry. In macaques, four DCN embedded within the cerebellar white matter of each hemisphere were distinguished: the dentate nucleus (DN), anterior and posterior interposed nuclei (AIN and PIN), and the fastigial nucleus (FN). The AIN and PIN can be comparable to the emboliform and globose nuclei in humans (Tellmann et al., 2015). To illustrate the location of all DCN, we combined sagittal DEC-FOD and T2w images with the coronal T2w image and 3D views of segmented DCN in figure 10. The signal intensity differences and the border between four DCN are more prominent in T2w than in MAP-MRI parameters, and these subregions are visible as hypointense in T2w due to their high iron content (Fig. 10D-G). DN is large and most lateral of the 4 DCN, and it exhibited more hypointense signal intensity than the two subregions of the interposed nuclei (AIN and PIN) and FN. These signal-intensity differences are comparable to the dark staining of neuropil in DN and moderate to light staining of neuropil in other DCN nuclei in AchE (compare Figs. 10G and 11H).

### Amygdala (Fig. 12)

The nuclear subdivisions of the amygdaloid complex follow (Price et al., 1987), as slightly modified by (Amaral and Bassett, 1989; Pitkanen and Amaral, 1998). Three of the MAP-MRI parameters (DEC-FOD, *RTAP*, and NG), and matched histology sections stained with ChAT, AchE, and SMI-32 are effective in specifying structural differences between different subregions of the amygdala. The deep nuclei of the amygdala are divided into lateral (L), basal (B), accessory basal (AB), and their subregions, as shown in the mid-level of the amygdala in coronal sections (Fig. 12). The lateral nucleus is further divided into dorsal and ventral regions (Ld and Lv), and both subregions exhibited more hypointense signals than the medially adjacent intermediate (Bi) and parvicellular (Bpc) subdivisions of the basal nucleus in *RTAP* and *NG* images (Fig. 12B, C). The signal intensity in the magnocellular (Bmc) division of the basal nucleus is comparable to that of the lateral nucleus in these MRI parameters. In contrast, the ChAT and AchE stained sections revealed darker staining of neuropil in different subregions of the basal nuclei (Fig. 12D, E), and this pattern of staining is the opposite of the signal intensity seen in these nuclei in NG and *RTAP*. The signal intensity of basal nuclei is comparable to the lateral in DEC-FOD, although the signal appears weaker in Ld (Fig. 12A). In SMI-32 stained sections, we found dark and patchy labeling of neuropil within Ld (Fig. 12F), and this localized staining suggests more compartments within this subregion of the lateral nucleus.

**Fig. 12.**
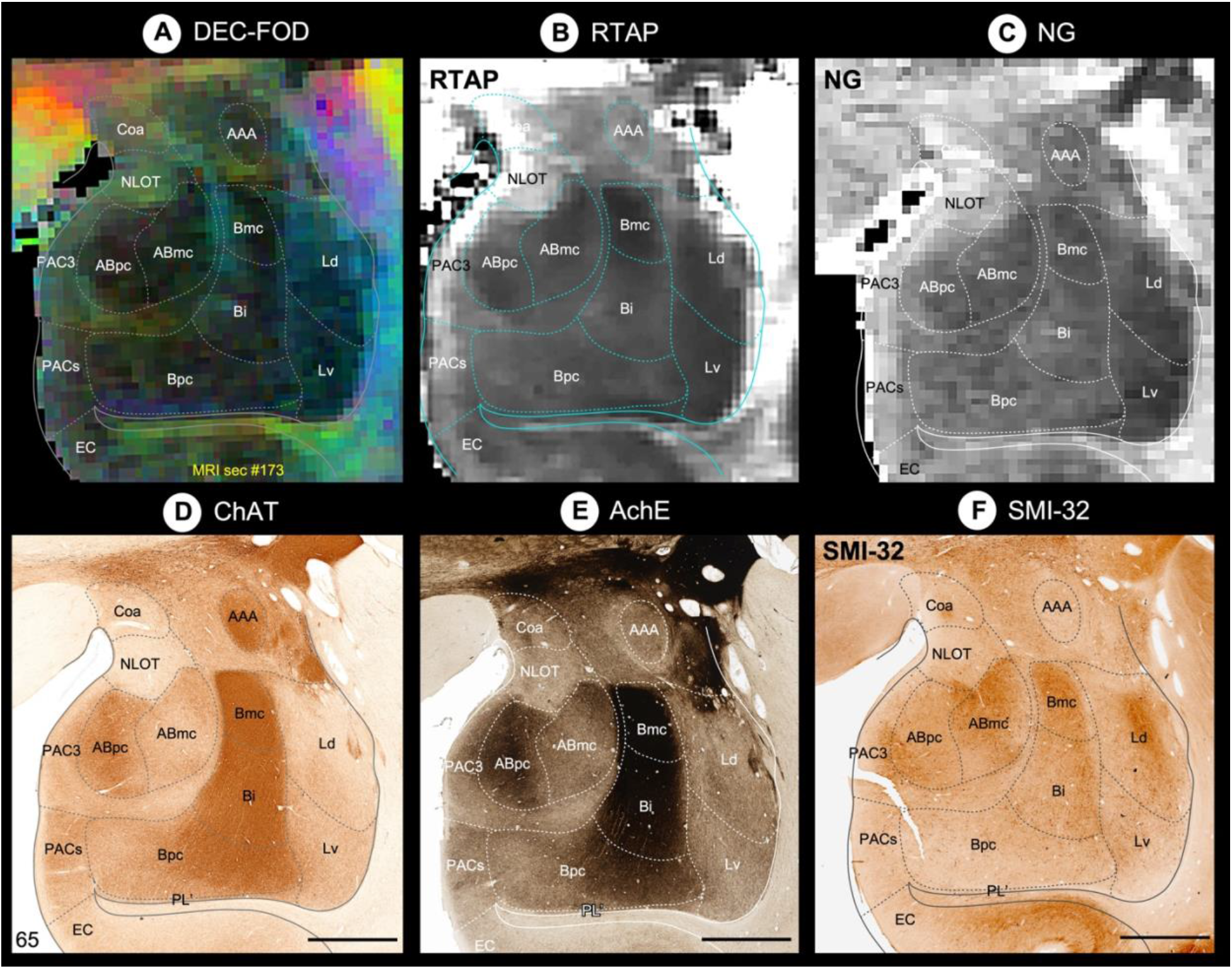
Amygdala. **(A-F)** Subregions of the amygdaloid complex in different MAP-MRI (DEC-FOD, *RTAP, NG*), and corresponding histology sections stained with ChAT, AchE, and SMI-32 (Section # 65). This coronal slice corresponds to the midportion of the right amygdala. The dashed outlines of different subregions on the MAP-MRI images are based on the ChAT- stained section (D). The lateral is on the right and the dorsal is on the top. ***Abbreviations***: AAA-anterior amygdaloid area; ABmc- accessory basal nucleus, magnocellular subdivision; ABpc- accessory basal nucleus, parvicellular subdivision; Bi-basal nucleus, intermediate subdivision; Bmc-basal nucleus, magnocellular subdivision; Bpc-basal nucleus, parvicellular subdivision; Coa- anterior cortical nucleus; EC-entorhinal cortex; Ld-lateral nucleus, dorsal subdivision; Lv-lateral nucleus, ventral subdivision; NLOT-nucleus of lateral olfactory tract; PAC3-periamygdaloid cortex 3; PACs-periamygdaloid cortex, sulcal portion; PL’- paralaminar nucleus. Scale bar: 2 mm (D-F).

The accessory basal nucleus is divided into magnocellular (ABmc) with weak neuropil staining and parvicellular (ABpc) with strong neuropil staining in ChAT and AchE stained sections (Fig. 12D-E). In MAP-MRI, both subregions revealed hypointense signals, but are distinct from the surrounding hyperintense anterior cortical area (Coa), the nucleus of the lateral olfactory tract (NLOT), and periamygdaloid cortex (PAC) (Fig. 12A-C). These latter regions are lightly stained in ChAT, AchE, and SMI-32 sections (Fig. 12D-F). The anterior amygdaloid area (AAA), which lies dorsal to the Bmc, exhibited more hyperintense in *RTAP* than DEC-FOD and *NG*. The other subdivisions of the amygdala, which are located caudal to the illustrated section (posterior cortical nucleus-Cop, central nucleus-CE, medial nucleus-ME, and amygdalohippocampal area- AHA), also showed hyperintense signals in different MR images (not shown in the illustration). We illustrated all subregions of the amygdala in our previous version of the D99 atlas, which is based on Saleem and Logothetis (2012) atlas.

### Hippocampal formation (Fig. 13)

Although unrelated to this study, the distinct architectonic characteristics of the gray and white matter regions in the hippocampal formation (hippocampus proper and subicular complex), prompted us to investigate and segment these allocortical areas in our MAP-MRI parameters with reference to various histological stains. Three of the MAP-MRI parameters (*AD, RTPP*, and DEC-FOD), and matched histology sections stained with SMI-32, PV, and AchE are effective in specifying structural differences between different subfields and their laminar subdivisions in the hippocampal formation (HF). Nine distinct subfields were identified (Palomero-Gallagher et al., 2020; Rosene DL and Van-Hoesen GW, 1987) and delineated within HF: Fascia dentata (FD), Cornu Ammonis (CA1, CA2, CA3, and CA4), prosubiculum (proS), subiculum proper (Sub), presubiculum (preS), and parasubiculum (paraS) (Fig. 13A-F), and a transition region at the rostral part of the HF, the hippocampal amygdaloid transition area (HATA). The FD, together with the CA4 region constitutes the dentate gyrus/DG (Palomero-Gallagher et al., 2020). The spatial location, rostrocaudal extent, and neighboring relationship between these subregions are shown on different views in 3D, reconstructed through a series of 70x200 μm thick coronal MRI slices (Fig. 13G-I). Although different subregions of HF were identified in MAP-MRI, the exact borders of these subregions, including laminae/layers on the MAP-MRI were delineated based on the histology stains.

**Fig. 13.**
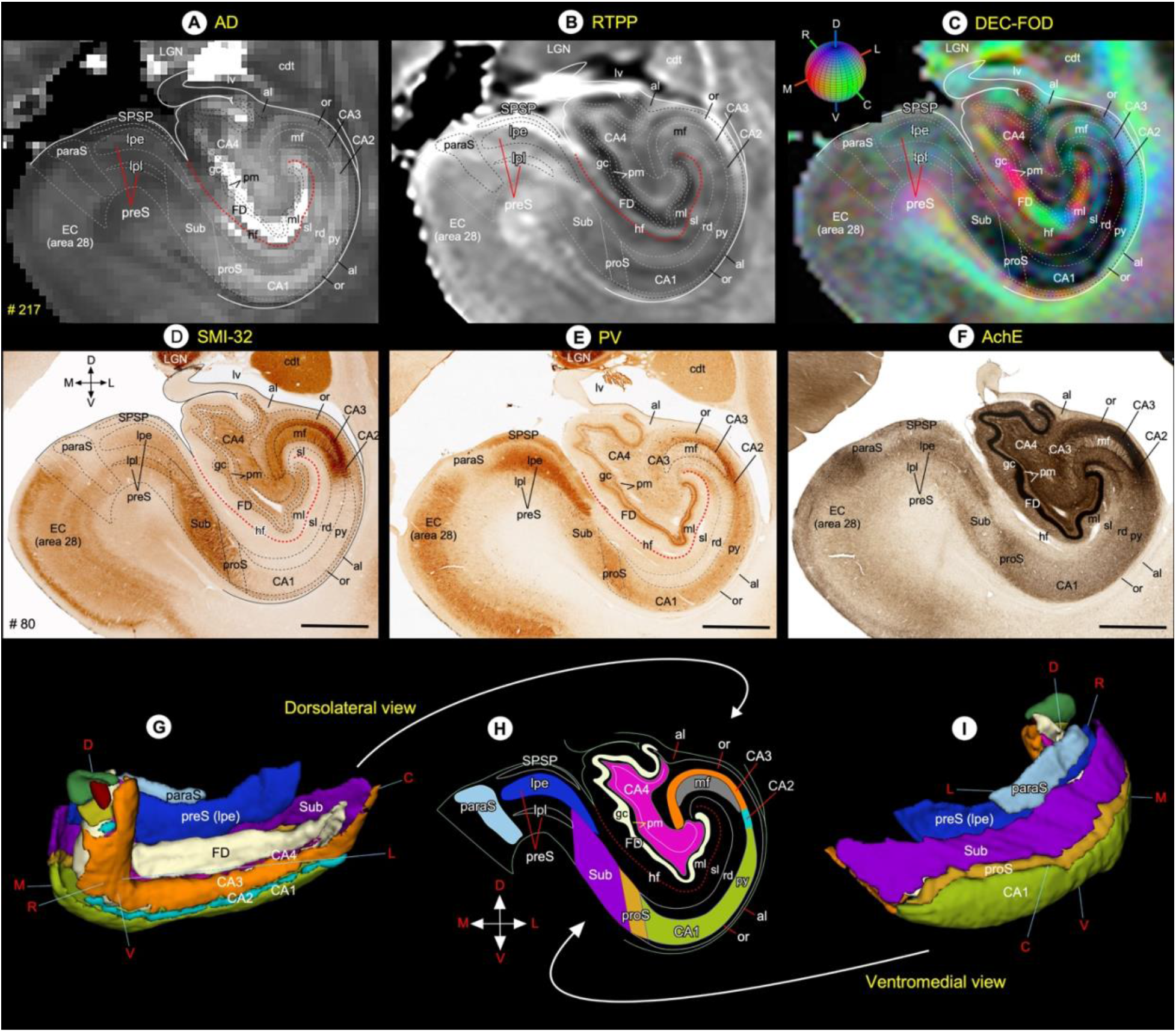
Hippocampal formation. **(A-F)** Subregions of the hippocampal formation in different MAP-MRI (*AD, RTPP*, DEC- FOD), and corresponding histology sections stained with SMI-32, PV, and AchE. The outlines of different subfields and layers on the coronal MAP-MRI images are based on the SMI-32 section (D). The red dashed line indicates the hippocampal fissure. **(G-I)** The rostrocaudal extent and spatial relationship between different subregions of the hippocampal formation in different angles in 3D, reconstructed through a series of 70x200 mm coronal MRI slices. The color-coded regions in “H” are reproduced from the SMI-32 section in “D”. Note that the signal intensity differences between layers of CA1 (py, rd, sl) in MAP-MRP parameter (AD) closely matched with the PV staining in E. ***Abbreviations***: al-alveus; CA1-CA4-subfields of the hippocampus; cdt-tail of the caudate nucleus; EC (area 28)-entorhinal cortex; FD-fascia dentata; gc-granule cell layer of the dentate gyrus; hf- hippocampal fissure; LGN-lateral geniculate nucleus; lv-lateral ventricle; mf-mossy fiber layer (stratum lucidum); ml-molecular layer; or-stratum oriens; paraS-parasubiculum; pm-polymorph cell layer of the dentate gyrus; preS-presubiculum; preS (lpe)-lamina principalis externa of the presubiculum; preS (lpi)-lamina principalis interna of the presubiculum; proS-prosubiculum; py- pyramidal cell layer; rd-stratum radiatum; sl-stratum lacunosum; SPSP-superficial presubicular pathway; Sub-subiculum. ***Orientation:*** D-dorsal; V-ventral; R-rostral; C-caudal; M-medial; L-lateral. Scale bar: 2 mm (D-F).

The fascia dentata (FD) is the most medial gray region, and it is separated from the subiculum, CA1, CA2, and part of the CA3 by the indistinct hippocampal fissure (hf) (Fig. 13). It is characterized by a densely packed layer of small granule cells (gc) that form an intensely stained dark layer in PV and AchE-stained sections (Fig. 13E, F). The gc layer is surrounded by a lightly stained superficial molecular layer (ml), dominated by the dendrites of the gc, and the deep polymorphic layer (pm), which is distinct from the clusters of darkly stained cells in CA4 (Fig. 13D-F). The FD (gc and part of the ml and pm) is strikingly distinguished as the most hyperintense region on AD and hypointense on *RTPP* images but is different from the signal intensity of the adjacent CA4 region (Fig. 13A, B). The subregions of FD are dominated by fiber bundles of different orientations throughout its mediolateral extent on DEC-FOD image, as illustrated in Figure 13C.

The CA3 is an arch-shaped narrow region situated between CA4 medially and CA2 laterally (Fig. 13). In the SMI-32, PV, and AchE sections, the CA3 is distinguished into a lateral region with intensely stained neuropil and a medial region with lightly stained neuronal processes, but this subfield is different from the staining intensity of the CA4 subfield (Fig. 13D-F).

Immediately ventral to the CA3 is the well-defined SMI 32-rich but PV and AchE-poor zone called the mossy fibers (mf or stratum lucidum), which demarcates the lateral border of the CA3 and the beginning of the CA2 subfield (Fig. 13D-F). The mossy fibers are the efferents of the granule cells of the FD and project to the CA3 pyramidal neurons (Kondo et al., 2008). The CA2 is a small region located between CA3 and CA1. The overall staining intensity of CA2 is similar to that of the CA3 in our illustrated histology materials, except that this region does not receive mossy fiber input from FD (Rosene DL and Van-Hoesen GW, 1987). Based on this criterion, CA2 lacks the mossy fiber layer or stratum lucidum, and this feature is readily observed in histology sections. In SMI-32, the CA2 is also characterized by the presence of a large number of intensely stained neuronal processes in the radiatum (rd), a layer adjacent to the pyramidal layer (py), but this architectonic feature is absent in most of the rd layer in CA3, below the mf layer (Fig. 13D). In MAP-MRI (*AD*), the lateral part of the CA3 and mf layer exhibited lower signal intensity than the medial part of the CA3 and CA2 (Fig. 13A). In contrast, the signal intensity differences between CA3/CA2 and adjacent layers are less prominent in *RTPP* and DEC-FOD images (Fig. 13B, C).

The CA1 is the largest subfield of the Cornu Ammonis, and it is readily demarcated from the adjacent darkly stained CA2 and prosubiculum (proS) in all illustrated histology sections. It is characterized by the presence of three prominent layers (py, rd, and sl-stratum lacunosum) with different neuropil staining intensities (Fig. 13D-F). These three layers of the CA1 are dominated by the cell body, proximal, and distal parts of the apical dendrites, respectively (Rosene DL and Van-Hoesen GW, 1987). In addition, the py and rd layers of CA1 receive projections through axon (Schaffer) collaterals from CA3 pyramidal cells (Kondo et al., 2009). The three layers of CA1 also exhibited different signal intensities in *AD* and *RTPP,* but its mediolateral border with proS and CA2 is less distinct (Fig. 13A, B). In contrast, the DEC-FOD image shows relatively low uneven anisotropy in all three layers of the CA1, and adjacent proS, as reflected in reduced color saturation (Fig. 13C).

The subiculum proper (Sub) is located between the proS and presubiculum (preS), and it is characterized by the presence of light- or darkly stained neuropil in SMI-32, PV, and AchE sections (Fig. 13D-F). This staining pattern is the opposite of the proS, parasubiculum (paraS), and two subregions of the preS (lamina principalis externa-lpe; and lamina principalis interna- lpi). These subregions revealed different signal intensities on the DEC-FOD image, and they closely matched with the neuropil staining of these subregions in the SMI-32 stained section (Fig. 13C, D). In contrast, the signal intensity differences between these subregions are less prominent in *AD* and *RTPP* images (Fig. 13A, B).

A number of major fiber bundles and interhemispheric crossing fibers are associated with hippocampal formation and are described in detail in the supplementary section (see Inline Supplementary Fig. 5 and related text).

### Validating SC21 in a test subject with histological confirmation (Fig. 14)

In a different subject, we demonstrated accurate matching of subcortical regions determined by MRI registration with those identified using architectonic analysis of histological sections from the same brain (Fig. 14). Here the segmented MRI subcortical template SC21 was registered to the *in vivo* T1-weighted MRI volume of one individual brain referred to as case MQ. We identified and matched subcortical areas in the coronal slice of registered brain volume (Fig. 14A) with the corresponding histology section, processed immunohistochemically with an antibody against the non-phosphorylated epitope of the neurofilament protein, recognized by the SMI-32 antibody. In the registered brain volume of MQ, the subregions of the basal ganglia (caudate-cd, putamen-pu, and substantia nigra-SN), thalamus (e.g., MDmc/pc/mf, VPM, VPI, cnMD, Pf, LGN), brainstem (VTA, RN, IPN, PN), and the hippocampal formation (CA1-4 and subiculum) closely matched the areas that we identified in the corresponding histology section from the same case (Fig. 14B). The chemoarchitectonic features of these selected areas were previously identified in a similar staining method described here (Saleem and Logothetis 2012). For the abbreviation of all illustrated regions, see figure 14 legend.

**Fig. 14.**
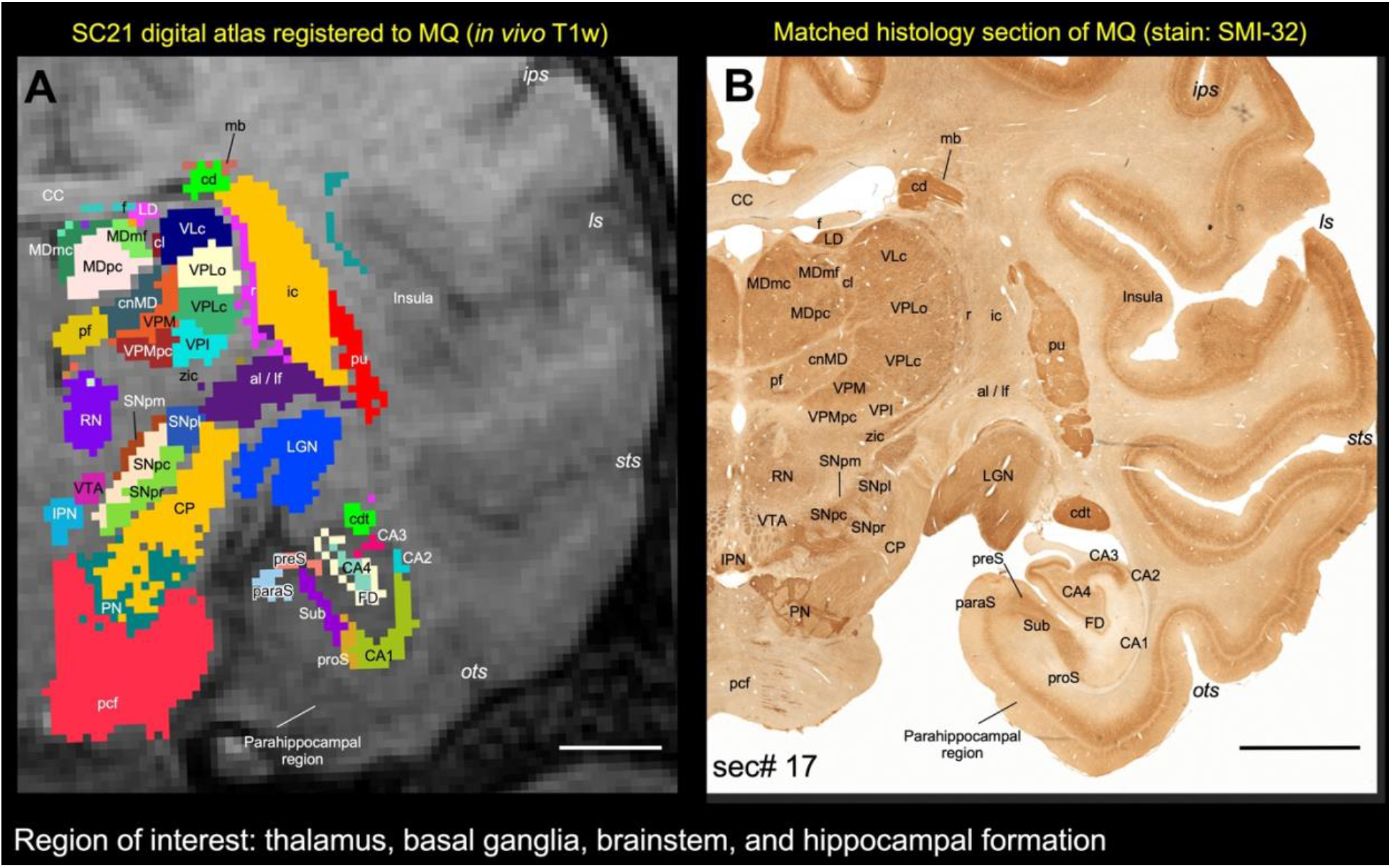
Registration of subcortical atlas to a different test subject with histological confirmation of architectonic areas. In this example, the segmented subcortical 3D volume (SC21) is registered to the T1w MRI volume of a different individual brain (Case MQ). **(A)** Shows the registered subcortical areas from SC21 overlaid on the MQ coronal MRI slice. None of the registered regions were altered or adjusted in this MRI slice or 3D volume. **(B)** Matched histology section from MQ stained immunohistochemically for the neurofilament protein, recognized by SMI-32 antibody (sec# 17). For consistency we flipped both MRI and histology in this illustration (i.e., left hemisphere is on the right). We digitally rotated the T1w MRI volume to match with the histology sections before the registration process. Note the correspondence of sulci and gyri in both the registered volume and histology section. We also confirmed the spatial location and the architectonic features of the selected subcortical targets (subregions of the thalamus, basal ganglia, brainstem, and hippocampal formation) in the registered coronal slice with the corresponding histology section as illustrated in A and B. For more details on the subregions of these deep brain structures see the result section. **Abbreviations**-***Thalamus:*** cl-central lateral nucleus; cnMD-centromedian nucleus; LD-lateral dorsal nucleus; LGN-lateral geniculate nucleus; MDmc-medial dorsal nucleus, magnocellular division; MDmf-medial dorsal nucleus, multiform division; MDpc-medial dorsal nucleus, parvicellular division; Pf-parafascicular nucleus; r-reticular nucleus; VLc-ventral lateral caudal nucleus; VPI-ventral posterior inferior nucleus; VPLc-ventral posterior lateral caudal nucleus; VPLo-ventral posterior lateral oral nucleus; VPM-ventral posterior medial nucleus; VPMpc-ventral posterior medial nucleus, parvicellular division; zic- zona inserta. ***Basal ganglia and related fiber tracts:*** al-ansa lenticularis; cd-caudate nucleus; cdt-caudate tail; ic-internal capsule; lf-lenticular fasciculus; mb-Muratoff bundle; pu-putamen; SNpc: substantia nigra, pars compacta; SNpl: substantia nigra, pars lateralis; SNpm: substantia nigra, pars mixta; SNpr: substantia nigra, pars reticulata. ***Brainstem structures:*** CP-cerebral peduncle; IPN-interpeduncular nucleus; pcf-pontocerebellar fibers; PN-pontine nuclei; RN-red nucleus; VTA-ventral tegmental area. ***Hippocampal formation:*** CA1-CA4-subfields of the hippocampus; FD-fascia dentata; paraS-parasubiculum; preS- presubiculum; proS-prosubiculum; Sub-subiculum. ***Other structures:*** CC-corpus callosum; f-fornix. ***Sulci:*** ips-intraparietal sulcus; ls-lateral sulcus; ots-occipitotemporal sulcus; sts-superior temporal sulcus. Scale bar: 5 mm (A and B).

### NEW digital template atlas “D99” with combined cortical and subcortical regions (Fig. 15)

We registered SC21 subcortical templates to the *ex vivo* MTR D99 surrogate volume with cortical segmentation using a sequence of affine and nonlinear registration steps (see method section). An initial affine step gave an approximate scaling and rotation to the template. The affinely warped subject brain was gradually warped to the template by progressively smaller nonlinear warps. This procedure resulted in the SC21 data in register with the D99 template space, showing both cortical and subcortical parcellations in the same volume. Figure 15A-C illustrates the new D99 digital template atlas with cortical and subcortical regions in the horizontal, coronal, and sagittal planes of sections. We mapped a total of 368 (149 cortical+219 subcortical) regions in this atlas (see suppl Table 1). It is important to note that the 219 parcellated subcortical regions also include the cerebellar cortex and the hippocampal formation, as indicated above. Figure 14D-E also shows the spatial location of delineated subcortical regions in 3D on the dorsal and lateral views. This new three-dimensional “D99” template atlas (version 2.0) is intended for use as a reference standard for macaque neuroanatomical, functional, and connectional imaging studies involving both cortical and subcortical targets.

**Fig. 15.**
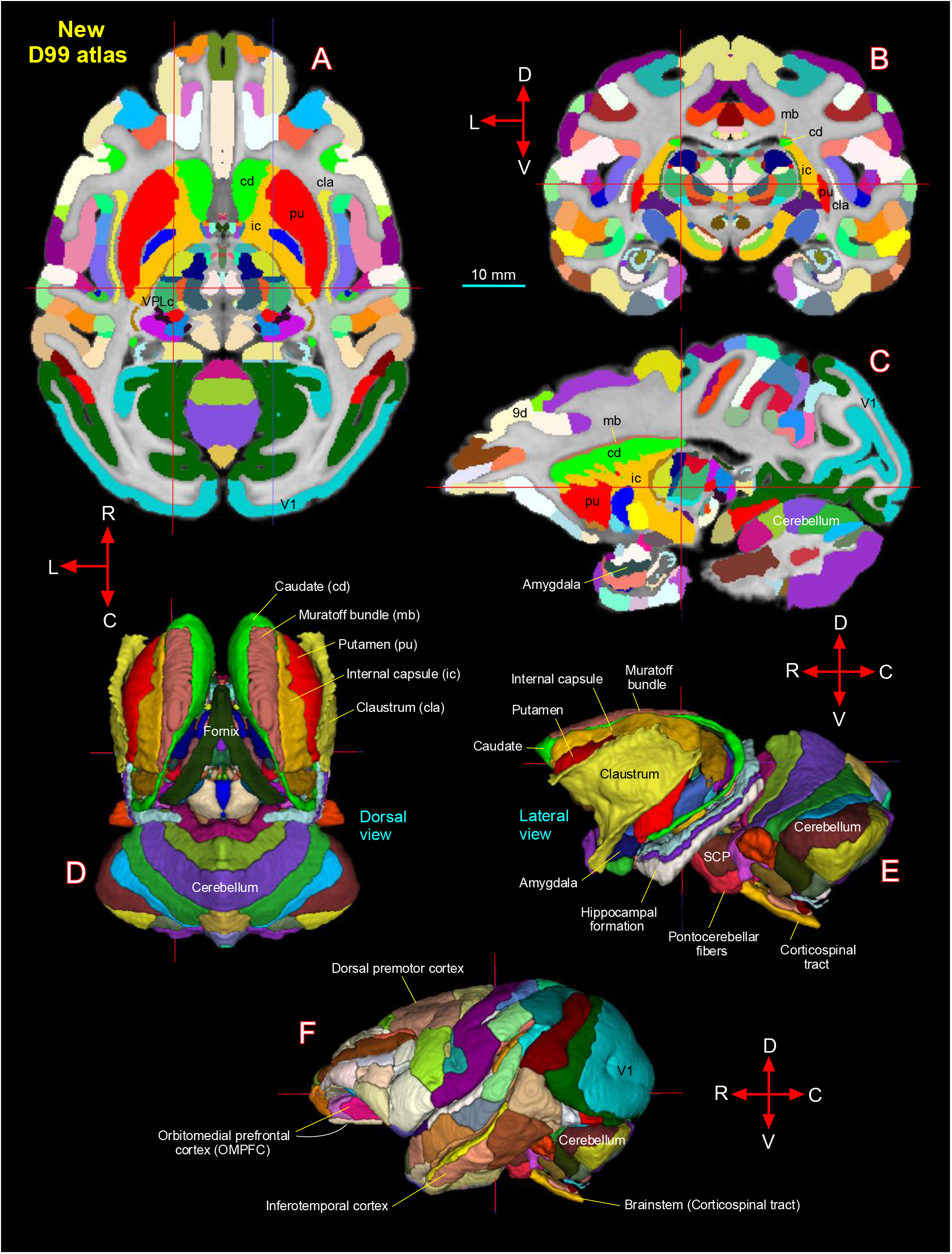
**[A-C]** New D99 digital atlas (version 2.0) with combined cortical and subcortical segmentation overlaid on the horizontal, coronal, and sagittal D99 ex-vivo MRI template. The cross-hairs in A-C show the same location of thalamic subregion VPLc (ventral posterior lateral caudal nucleus). **(D-E)** The spatial location of segmented subcortical regions shown on the dorsal and lateral views in 3D. The selected subcortical regions in D-E are also indicated with cortical areas in A-C. **(F)** Segmentation of cortical areas. ***Abbreviations:*** 9d-dorsal prefrontal area; SCP-superior cerebellar peduncle; V1-primary visual cortex. ***Orientation:*** D-dorsal; V-ventral; R-rostral; C-caudal; L-lateral. Scale bar: 10 mm applies to A-C only.

With AFNI’s @animal_warper, the D99 atlas can be automatically registered to the 3D anatomical scans from a wide range of macaque subjects (see next section, Fig. 16), and corresponding functional (fMRI) scans (Reveley et al., 2017; their Fig. 8), and thus used to specify the areal designation relative to experimental locations of interest.

**Fig. 16.**
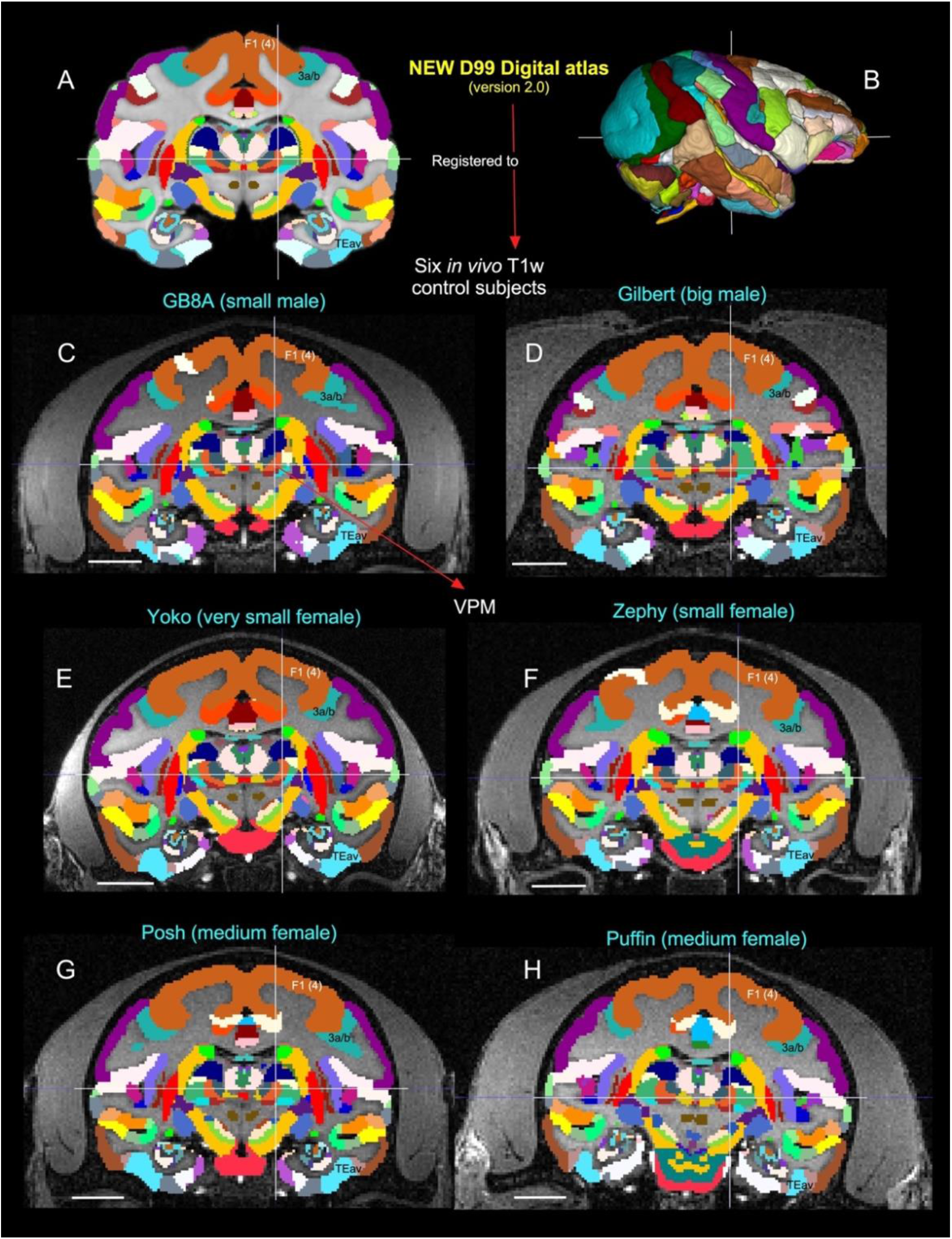
Validation of 3D atlas. Registration of new D99 digital atlas (version 2.0) to various *in vivo* T1w test subjects of different age groups using a novel-processing pipeline developed within AFNI and SUMA (see the method section). **[A]** Mid coronal section from D99 atlas with delineated cortical and subcortical regions. **[C-H]** Coronal slices from 6 control animals, with the D99 atlas registered to the T1w MRI images of each animal in its own native space. None of the registered regions were altered or adjusted in these animals, and for consistency the width of the slice in each animal is matched with D99. Note the corresponding location of selected subcortical region VPM in the lateral thalamus, indicated by cross-hair, and cortical areas F1 (4), 3a/b, and TEav in D99 digital atlas and 6 other animals. The corresponding location of VPM is also shown with the D99 rendered brain volume in **B**. ***Abbreviations:*** 3a/b-somatosensory areas 3a and 3b; F1 (4)-agranular frontal area F1 or area 4; TEav-ventral subregion of anterior TE; VPM-ventral posterior medial nucleus. Scale bar: 10 mm (C-H).

### Registering new D99 template atlas to a range of *in vivo* test subjects (Fig. 16)

To this end, we developed a novel macaque processing pipeline within AFNI and SUMA to optimally register the new D99 atlas to T1w-MR images of individual macaque brains. The results of this pipeline are illustrated in Figure 16 for six monkeys of different ages, genders, and sizes. In this example, the corresponding location of a subcortical region at the cross-hair, the VPM (ventral posterior medial nucleus of lateral thalamus), and cortical areas, e.g., F1 (4), 3a/b, and, TEav matched well in both D99 atlas and six other animals (Fig. 16C-H). These results demonstrate that affine and nonlinear warpings are sufficient to distinguish and provide atlas- based estimates of areal boundaries of macaque subjects *in vivo*.

Atlases and templates are available as both volumes and surfaces in standard NIFTI and GIFTI formats. While this 3D digital atlas can be used in different image registration and analysis software packages, here we use the AFNI and SUMA programs with their advanced atlas features for purposes of demonstration (Cox, 1996; Saad and Reynolds, 2012). The atlas is integrated into the most recent versions of AFNI and SUMA, making for straightforward identification of areal identity in any macaque subject registered to the template and for the individual macaque subject in its own native space by the inverse transformations. The 3D template volume and atlas, and script for atlas registration of *in vivo* scans are now available for download through the AFNI and SUMA website, at https://afni.nimh.nih.gov/pub/dist/atlases/macaque/D99_Saleem/D99_v2.0_dist.tgz. The AFNI software can install this simply with @Install_D99_macaque command. For other details, see the “*implementation and distribution*” section in (Reveley et al., 2017).

## Discussion

Despite its essential use as a model for mental illness in humans, to date the rhesus macaque lacks comprehensive well-organized MRI-histology based templates of subcortical regions. Here, we have described the spatial location and detailed microarchitectural characteristics, and boundaries of subcortical regions of the macaque monkeys using *ex vivo* MAP-MRI at high resolution (i.e., 200 μm), aided by postmortem histology of the same brain. Our results demonstrate that, at the high spatial resolution, MAP-MRI in particular, can distinguish a large number of gray matter structures such as sensory and motor nerve nuclei in deep brain structures. Additionally, the combined use of nine microstructural MRI parameters and five histological stains enabled detailed noninvasive mapping of tissue structures, including larger and smaller fiber pathways and cytoarchitectonic features in the basal ganglia, thalamus, brainstem, and limbic structures. In contrast, the anatomical details that are evident in MAP-MRI parameters are invisible in typical single- or multi-subject T1w templates (see Inline Supplementary Fig. 6). We believe that our current findings have produced a more objective and reproducible description of subcortical targets in macaque monkeys. In the following sections, we first discuss our findings along with previous studies of these deep brain targets in primates and then compare our new 3D digital template atlases of the rhesus macaque (SC21, and updated D99) with other available atlases in the field. Finally, we highlight the potential use of DTI/MAP parameters to construct the atlases of deep brain structures in humans.

### Mapping of subcortical gray and white matter regions at high-field MRI in primates

The high-resolution *ex vivo* MAP-MRI with nine microstructural parameters and T2-weighted images revealed sharp borders and high contrast between gray and white matter (GM/WM) outside the cerebral cortex, resulting in the precise demarcation of fine anatomical structures such as nuclei and sub-laminae in subcortical regions (e.g., basal ganglia, thalamus), consistent with the histological sections with multiple stains prepared from the same brain. Our *fODF*s- derived directionally encoded color (DEC) maps also revealed several nuclei and the orientation of small and large fiber tracts that link basal ganglia with the thalamus (e.g., lf-lenticular fasciculus, Fields of Forel H, H1, H2) and other long-range fiber bundles such as corticospinal tract, pontocerebellar fibers, and medial lemniscus (Fig. 10). The fields of Forel in human brains are the most recognized regions in deep brain stimulation of the subthalamic nucleus in Parkinson’s disease (Neudorfer et al., 2017). Our high-dimensional DTI/MAP parameters and T2w images also revealed microarchitectural details within the subcortex (e.g., striatum; Fig. 4) that can be comparable to the neurochemically defined compartments with distinct input and output connections called Patch/striosomes and matrix (Brimblecombe and Cragg, 2017; Crittenden and Graybiel, 2011; Eblen and Graybiel, 1995; Gerfen, 1989; Graybiel and Ragsdale, 1978; Hirsch et al., 1989; Martin et al., 1993; Smith et al., 2016).

Our high-dimensional MAP-MRI parameters showed several laminae separating subcortical gray matter regions in the basal ganglia and thalamus. These are the lateral and medial medullary laminae (lml and mml, respectively) located between the putamen and pallidum, and the intermedullary lamina (iml), located between anterior and lateral thalamic groups. The lml and mml in macaque are comparable to the lamina pallidi lateralis and lamina pallidi medialis, respectively, in humans visualized in susceptibility-weighted imaging (SWI) at 7T *in vivo* (Abosch et al., 2010; Schaltenbrand and Wahren, 1977). Additionally, the third lamina (lamina pallidi incompleta) that subdivides GPi into lateral and medial divisions was found in human SWI (Abosch et al., 2010). Although, this lamina is absent or less prominent in the macaque in different MRI parameters, the lateral and medial subregions of the GPi are distinguished based on dark and light honeycomb-like neuronal processes, respectively, in our histology stains (magnified regions in Fig. 5G-I).

The location of basal ganglia subregions, such as striatum (caudate and putamen), substantia nigra, subthalamic nucleus, and subregions of globus pallidus and some of these fiber bundles in macaque monkeys is comparable to humans (Deistung et al., 2013a; Straub et al., 2019; Tang et al., 2018). In humans, many of the deep brain structures were mapped *in vivo* or *ex vivo* with ultrahigh-resolution MRI using multiple image contrasts (Abosch et al., 2010; Deistung et al., 2013a; Deistung et al., 2013b; Keuken et al., 2014; Lenglet et al., 2012). These deep brain structures represent common targets for deep brain stimulation (DBS) (for review see (Plantinga et al., 2014) for treating cognitive dysfunction in patients with traumatic brain injury (TBI), motor-related symptoms in Parkinson’s disease, and neuropsychiatric disorders (Kundu et al., 2018; Sullivan et al., 2021; Wichmann and Delong, 2011). Taken together, MAP-MRI with different contrasts could facilitate accurate delineation of the potential DBS targets in the macaque model of Parkinson’s disease and other psychiatric illnesses (Min et al., 2016; Vitek and Johnson, 2019; Xiao et al., 2016).

Our histochemical and immuno-histochemical stained sections also revealed rostral and caudal subregions of the red nucleus (RNpc and RNmc, respectively) with different chemoarchitectonic features, but these subregions were less conspicuous from the surrounding gray and white matter regions in T2w images (Fig. 7). In macaques, both the substantia nigra (SN) and the globus pallidus (GP) revealed a more hypointense signal than RNpc/mc in T2w images (Fig. 7I), probably due to the high level of iron content in the SN and the GP. In contrast, the RN demonstrated a clear boundary with significantly decreased (hypointense) signal, similar to the GP and the SN in T2- and susceptibility-weighted MRI in healthy human subjects (Abosch et al., 2010) and T2w images in postmortem human brain specimens (Massey et al., 2012). Thus, the T2w MRI marker is not suitable for delineating the RN in macaque monkeys. We also identified the medial and lateral subregions of the habenular complex based on the MR contrast in MAP-MRI (DEC-FOD), and staining differences of neuropil in the corresponding histological sections stained with SMI-32 (Fig. 9K). The distinction between the medial and lateral parts of the habenular complex observed in the macaque can be comparable to the human habenular complex (Diaz et al., 2011). It is worth noting that the subregions of the red nucleus and the habenular complex (including the substantia nigra) were not parcellated in a recently developed subcortical atlas displayed on NMT2, called SARM (Hartig et al., 2021). In the SARM atlas, despite its undefined shape in the underlying MRIs, the red nucleus (RN, for e.g.,) was manually drawn onto the NMT2 volume based on the relative position of the neighboring structures (the VTA and SN), as illustrated in figure 12a in that atlas. Mapping the RN based on MRI alone, without corresponding histological images from the same brain to serve as a control, may produce inaccurate boundaries that include additional gray and white matter regions leading to biases in size/volume estimation.

### SC21/D99 Digital atlas versus other atlases

The 3D digital template atlas is intended for use in anatomical, connectional, or functional imaging studies using macaque monkeys. In particular, registering the 3D atlas to a given macaque’s brain can provide the spatial distribution of labeled axon terminals and neurons after the anatomical tracer injections, areal location of fMRI responses, or DBS targets for non-human primate’s model of Parkinson’s disease, dystonia, or other brain disorders. The present macaque digital template atlas of the subcortical regions derived from correlations between MRI and histology is one of the few digital monkey brain atlases that have been created in recent years (Calabrese et al., 2015a; Hartig et al., 2021; Rohlfing et al., 2012). In the following paragraphs, we briefly compare and contrast these atlases and highlight some advantages of the present digital template atlas.

Rohlfing and colleagues constructed a new *in vivo* T1w MRI template, termed INIA19 for imaging-based studies of non-human primate brains, with comprehensive cortical and subcortical labels derived from NeuroMaps atlas (Rohlfing et al., 2012). These authors pointed out that the segmentation of the hypothalamus, amygdala, and basal forebrain regions in their maps is incomplete, and regions in the brainstem (midbrain and hindbrain) are also not fully described. In another recent study, Calabrese and colleagues (Calabrese et al., 2015a) presented MRI-DTI based atlas of rhesus macaque brain based on 10 postmortem brain specimens and provided detailed three-dimensional segmentation of major cortical areas and white matter pathways, and a number of subcortical regions (part of the basal ganglia, thalamus, and amygdala). However, this study did not provide the parcellation of brainstem structures. The anatomical segmentation in their map was initialized through automated label registration (ALR) using labels from the histology-based atlas of a different macaque monkey brain (Paxinos et al., 2009). While useful for some applications, this automated ALR does not attempt to preserve the native geometry of the brain, thus the registration accuracy of the resulting volumetric or surface atlas will always be in question.

A recent study generated a subcortical atlas of the rhesus macaque or SARM (Hartig et al., 2021) based on a single subject *ex vivo* structural scan for MRI data analysis, but it relied on 2D histology materials obtained from a different subject (Paxinos et al., 2009) to delineate subcortical regions on *ex vivo* MR images. This anatomical scan and its subcortical parcellations were then mapped onto the T1-weighted NMT-v2.0 *in vivo* population template (Jung et al., 2021). Because subcortical gray matter structures (e.g., thalamic- and brainstem nuclei) are usually concentrated in a small region of the brain with uncertain borders, delineating architectonic regions (nuclei) on *ex vivo* or *in vivo* structural scans such as T2w and T1w MRI alone without corresponding matched histological information from the same subject is prone to be inaccurate. In addition, the fiber pathways of different sizes and orientations course through deep brain structures, and delineation of nuclei from fiber bundles requires more than typical or population-averaged structural scans. Additional information like diffusion MRI with directionally encoded color (DEC) maps and other DTI/MAP parameters is critical to assist neuroanatomical delineation of these structures in detail. Other MRI parameters such as those obtained from quantitative susceptibility mapping (QSM) can be used to delineate the overall border of some subcortical structures like substantia nigra in both younger and older macaque monkeys *in vivo* better than the conventional T1w images (Yoshida et al., 2021).

As shown in the results section, high-resolution DTI/MAP provides new microstructural parameters and directional information (*fODF*s/DEC) that can complement multiple histological stains. Collectively, these are crucial in delineating nuclei and fiber tracts of different sizes and orientations in deep brain structures (e.g., pontocerebellar fibers from the pyramidal tract, medial lemniscus, and pontine nuclei; figs. 10, 11). Many of these deep brain targets and their subregions, are less prominent or spatially not distinguishable from neighboring structures with a single MRI parameter, or a population-averaged anatomical MRI volume NMT2 (Jung et al., 2021) (see Inline Supplementary Fig. 6). The most important unique feature of our subcortical atlas (SC21) is the strict adherence to an MRI scan with the adjacent and matched histology sections from the same brain. As a result, the alignment accuracy between areal boundaries and gross anatomical features is optimized for the identification of region-of-interest in this case (e.g., Figs. 9, 11). The subcortical SC21 volume is also registered to a standard high-resolution D99 atlas with cortical parcellation (Reveley et al., 2017; Saleem and Logothetis, 2012), using widely available tools of whole-brain MRI registration. The transformation derived from this warping, when then applied to the 3D digital atlas volume, allows for labeling of both cortical and subcortical targets in the brains of individual animals as accurately as possible.

### Validation of 3D atlas

We performed two analyses that indicate that validation of SC21 atlas or an updated D99 atlas, such as estimating the architectonic boundaries between different brain areas for a population of *in vivo* T1w macaque brains is generally good and useful. In the first analysis, we showed a good match between subcortical areas in a new monkey subject (MQ) using SC21 digital template registered to MQ with histological confirmation of architectonic areas obtained from the same animal (Fig. 14). The validation of brain regions in a given subject is useful for neurosurgical navigation of electrode or implantable devices to a potential target for deep brain stimulation (DBS) in the macaque model of psychiatric or neurological disorders (e.g., Parkinson’s disease (Min et al., 2016; Vitek and Johnson, 2019). In the second analysis, we revealed that the MRI registration procedure using updated D99 (Fig. 15) can be smoothly applied to test subjects of different genders, age groups (1.2–14.8 years old), and sizes (2.55 and 5.5 kg). Thus, it is possible to estimate histological boundaries of cortical and subcortical areas in any monkey subject (Fig. 16).

### The potential of MAP-MRI based atlases in humans

In this study, we mapped 3D subcortical structures using MAP-MRI data acquired with very high resolution. We further illustrated how our atlas can be used to locate small subcortical structures in a test monkey based on a single conventional T1w image acquired *in vivo* and verified the accuracy of the localization with histological brain sections from the test monkey. These results suggest the utility of high-resolution atlases in studies of macaque monkey disease models and highlight the potential of high-resolution MAP-MRI in delineating small subcortical structures based on differences in microstructural properties. The spatial resolution of MAP-MRI data acquired on clinical scanners are significantly lower, approximately 1-2 mm. Nevertheless, promising advances in gradient coil design (McNab et al., 2013), RF coil engineering (Keil et al., 2013; Truong et al., 2014), spatial encoding (Feinberg et al., 2010; Setsompop et al., 2018), and dMRI pulse sequence design (Avram et al., 2014b) are expected to significantly improve the spatial resolution and SNR to allow submillimeter clinical MAP-MRI scans in the near future (Huang et al., 2021). Concurrently, new clinically feasible diffusion encoding strategies (Avram et al., 2010; Avram et al., 2019; Avram et al., 2021) are being developed in order to quantify specific microscopic tissue water pools without the need to increase the spatial resolutions.

Taken together, these advances will enable the construction of high-resolution cortical and subcortical atlases of the human brain that will improve neurosurgical navigation, localization of fMRI responses, high-precision placement of recording and stimulating electrodes in patients with Parkinson, mTBI, epilepsy, and other diseases. MAPMRI has been applied to numerous studies using healthy volunteers (Avram et al., 2016) and patient populations with Parkinson’s Disease (Le et al., 2020), multiple sclerosis (Boscolo Galazzo et al.), ischemic stroke (Boscolo Galazzo et al., 2018), epilepsy (Ma et al., 2020), brain injury and cancer (Jiang et al., 2021).

### Summary and Conclusion

Using combined high-resolution DTI/MAP parameters and corresponding histological sections with multiple stains, we have mapped the spatial locations, boundaries, and micro-architectural features of subcortical gray and white matter regions in the macaque monkey *ex vivo*. The strengths of this work compared to others are that 1) it uses a large range of high-resolution microstructural DTI/MAP parameters with *fODF*s/DEC, 2) utilizes both high-resolution MRI (200 μm) and corresponding histology information from the same brain (i.e., MRI-histology correlations) to map anatomical structures, 3) provides a complete, accurate, and detailed mapping of subcortical structures, including the brainstem with the cerebellum, and 4) validates its application to datasets acquired *in vivo* with the second histological confirmation; a useful corroboration for identifying potential DBS targets in the macaque model of Parkinson’s disease and other neurological and psychiatric illnesses. Finally, an updated comprehensive MRI-histology-based D99 atlas with combined cortical and subcortical parcellations provides a readily available and usable standard for region definition in the brains of individual animals of different ages, genders, and sizes. The current atlas, template MRI data sets, surfaces, and user scripts for aligning individual subjects to this template are publicly available in the following link: https://afni.nimh.nih.gov/pub/dist/atlases/macaque/D99_Saleem/D99_v2.0_dist.tgz. The continued development of novel MRI stains and contrasts and sensitive, along with specific, quantitative histological stains should enable the elucidation of structures that were previously invisible radiologically. These developments can provide a continued impetus for various neuroscience explorations and furnish/put forward a roadmap for future neuroradiological developments to be able to scan salient brain areas *in vivo*.

## Credit author statement

***Kadharbatcha S. Saleem:*** first and corresponding author, designed the study, coordinated the project, prepared the specimen for MRI and histology, collected high-resolution images of histology sections; mapped, segmented, and verified all the anatomical regions of interest with reference to MRI and histology, generated new subcortical 3D atlas template/Table, made illustrations; wrote, edited, and streamlined the manuscript.

***Alexandru V Avram:*** first co-author, designed and conducted all MRI experiments including MAP-MRI scans, processed and analyzed all MRI data, helped with atlas data registration, wrote and edited the manuscript.

***Daniel Glen:*** Integrated the atlas dataset into AFNI and SUMA software packages, wrote code for the atlas region regularization, integration of cortical atlas and edited the manuscript.

***Cecil Chern-Chyi Yen:*** Assisted with MRI data collection and provided feedback on the manuscript.

***Frank Q Ye:*** Helped in optimization of MRI data acquisition and commented on the manuscript. ***Michal Komlosh:*** Prepared the specimen for MRI and assisted with MRI data collection, including MAP-MRI.

***Peter J Basser:*** helped design study, edited the manuscript, and provided research resources.

## ACKNOWLEDGMENTS

This work was supported by the Intramural Research Program of the *Eunice Kennedy Shriver* National Institute of Child Health and Human Development, the Intramural Research Program of the National Institute of Neurological Disorders and Stroke, “Connectome 2.0: Developing the next generation human MRI scanner for bridging studies of the micro-, meso- and macro-connectome”, NIH BRAIN Initiative 1U01EB026996-01 and the CNRM Neuroradiology/Neuropathology Correlation/Integration Core, 309698-4.01-65310, (CNRM-89- 9921). We thank Joel Price for comments on the manuscript; Drs. Betsy Murray and Richard Saunders, and Alex Cummins in the Laboratory of Neuropsychology, NIMH for providing perfusion-fixed rhesus monkey brains for our experiments; Drs. Bernard Dardzinski and Alexandru Korotcov for providing the RF coil used in this experiment; and Thomas J. Pohida and Marcial Garmendia-Cedillos for help with the 3D-printed brain mold. Finally, we thank Vincent Schram and his team in Microscope imaging core (MIC) at NICHD and Sarah Williams- Avram at NIMH for help with the high-resolution imaging of histology sections. All histological processing of the brain tissue was done by Dr. Du and his team at FD NeuroTechnologies in Columbia, Maryland.

## Supplementary material

**Inline Supplementary Figure 1** Blocking of the brain specimen with reference to MRI plane (see Data analysis section in Materials and methods). We first visually matched a number of reference points (e.g., sulci) on the MR images with similar locations on the surface of the 3D rendered brain volume generated from these MR images (top row). These sulcal points on the rendered brain surface were then manually translated onto the corresponding location of the brain specimen of this subject before blocking it (middle column). These steps enabled us to match the sulci, gyri, and the region of interest (ROI) in deep brain structures in both MRI and blockface/histological sections (left and right columns). The brain blocks were prepared for sectioning and staining, followed by high-resolution imaging of stained sections. ***Abbreviations:*** amts-anterior middle temporal sulcus; asd- anterior subcentral dimple; cs-central sulcus; ips-intraparietal sulcus; ls-lateral sulcus; spcd-superior precentral dimple; sts-superior temporal sulcus.

**Supplementary Figure 1.**
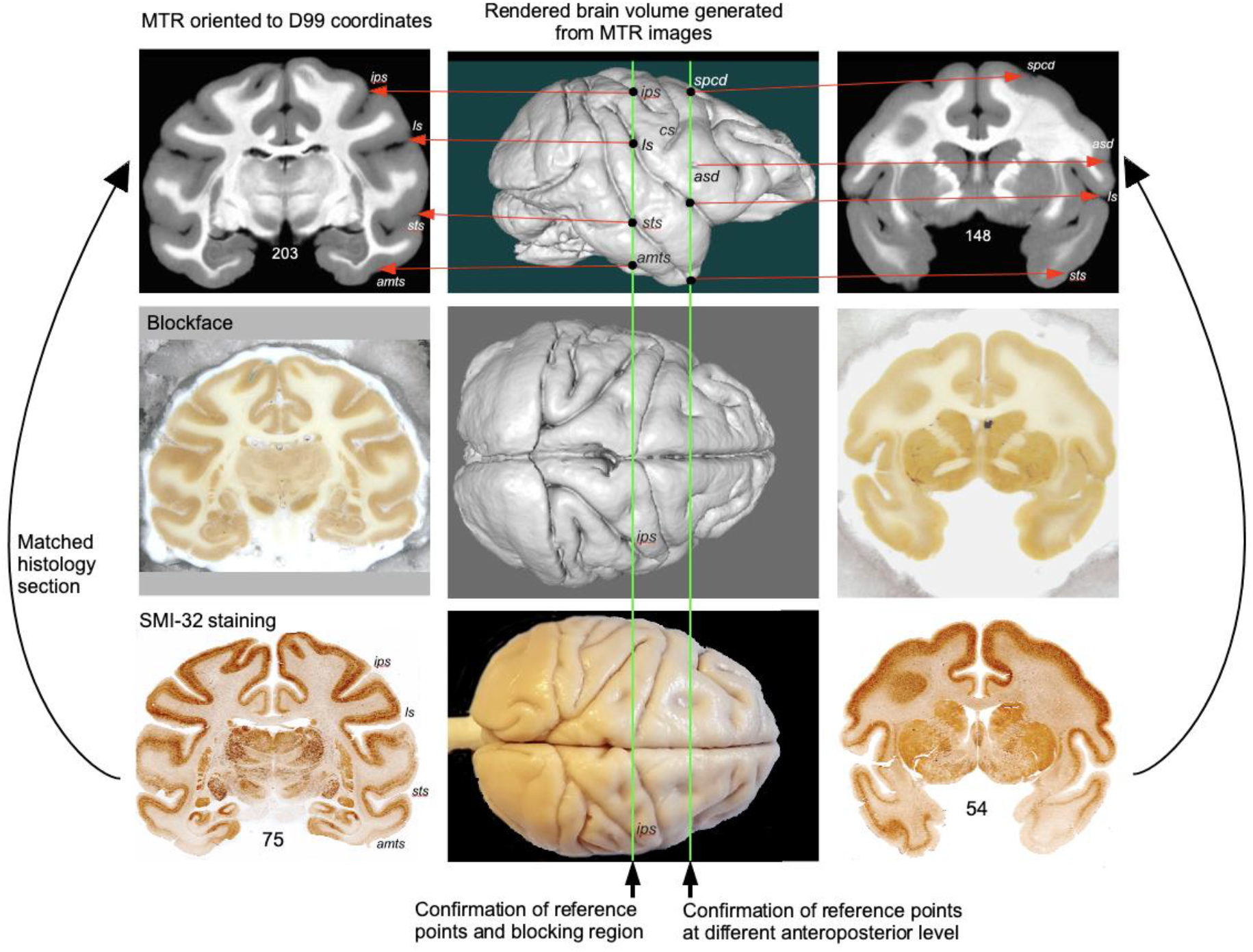
Blocking of the brain specimen with reference to MRI plane.

**Inline Supplementary Figure 2 –** Subthalamic nucleus (Basal ganglia)

The subthalamic nucleus (STN) is an obliquely-oriented almond-shaped structure, located dorsal to the cerebral peduncle (CP), dorsolateral to the substantia nigra (SNpr), and ventral to the zona inserta (zic) (Suppl Fig. 2). In non-human primate macaques, different subregions of STN along the rostrocaudal axis receive direct projections from the motor, prefrontal, and anterior cingulate areas coupled with indirect projections from the globus pallidus (Coude et al., 2018; Haynes and Haber, 2013; Nambu et al., 1996; Parent and Hazrati, 1995; Sato et al., 2000). The convergence of these projections in STN provides the anatomical substrate for integrating motor, emotional, motivational, and cognitive information toward the selection of complex behaviors, a key for our understanding of motor and non-motor effects of deep brain stimulation of this basal ganglia structure in Parkinson’s disease (Haynes and Haber, 2013; Mallet et al., 2007).

The STN is sharply defined from the neighboring gray and white matter structures (see above) and is readily identifiable as a hyper- or hypointense region on different MAP-MRI and T2w images (Suppl Fig. 2A-E). The STN is also distinguished as a hyperintense region on the MTR image, but its ventromedial boundary with the SNpr is less prominent (Suppl Fig. 2F). In contrast, the outer border of STN from the surrounding regions is less distinct in high-resolution multi-subject T1-weighted images, called NMT (Suppl Fig. 2G; (Seidlitz et al., 2018). In some MAP-MRI parameters, the internal signal characteristics of the STN are well defined. For example, in *RD*, patches of hyper- and hypointense regions were distinguished in the different mediolateral extent of STN that corresponded well with the dark- and lightly stained regions, respectively, in the AchE section (Suppl Fig. 2H, I; red boxes). These subregions of different signal intensities in STN can roughly correspond to the overlapping zones of convergent projections from the prefrontal, premotor, and primary motor areas as previously shown by anatomical tracing studies (Suppl Fig. 2H, inset; see (Haynes and Haber, 2013).

**Supplementary Figure 2.**
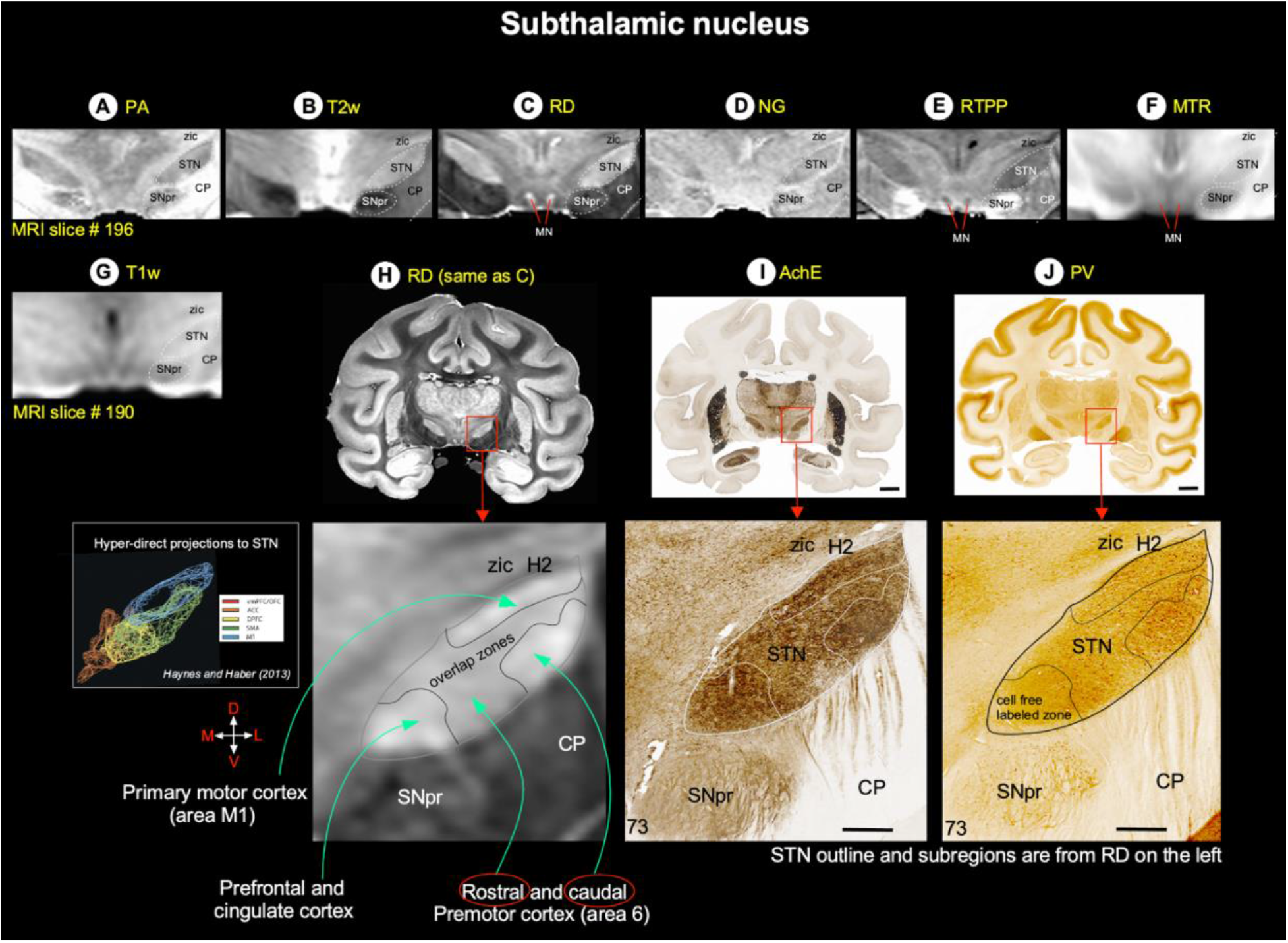
Subthalamic nucleus. **[A-G]** The spatial location of the subthalamic nucleus (STN, dashed outline) with reference to other gray and white matter structures in coronal MAP-MRI (PA, RD, NG, RTPP), T2w, MTR, and NMT images. NMT is a multi-subject in vivo T1w MRI volume (Seidlitz et al., 2018). **(H)** Zoomed-in view of the mid STN in MAP-MRI (RD) shows the mediolaterally restricted zones of different signal intensities, which can roughly correspond to the overlapping zones of convergent projections from diverse cortical areas as previously shown by anatomical tracing studies (inset; see Haynes and Haber, 2013). **(I-J)** The magnified view of the STN with outlined regions shows the compartments of dark- and lightly stained neuropil in AchE and PV sections. These outlines are derived from the matched MRI (RD) image on the left. **Orientation:** D-dorsal; V-ventral; M-medial; L-lateral. **Abbreviations:** CP-cerebral peduncle; H2-H2 field of Forel; SNpr- substantia nigra pars reticulata; zic-zona incerta. Scale bar: 5 mm (I, J); 1 mm applies to magnified regions from I-J (red boxes).

**Inline Supplementary Figure 3** - Thalamus

**Supplementary Figure 3.**
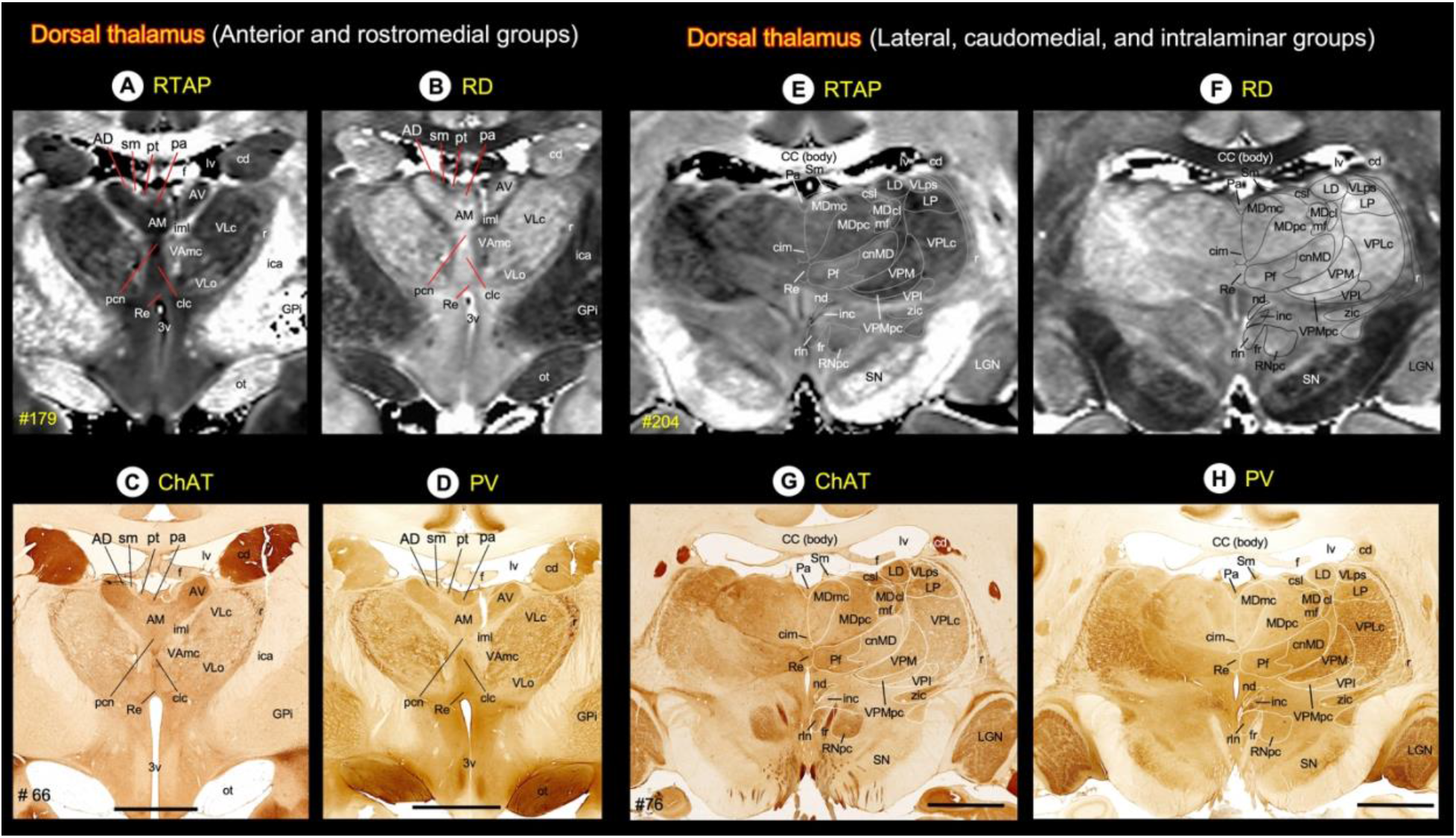
Thalamus. (Top row) MR signal intensity in different subregions of the dorsal thalamus (anterior, medial, and intralaminar groups) in RTAP and RD images. **(Bottom row)** The corresponding regions of the dorsal thalamus in ChAT and PV stained sections. **Abbreviations:** AD-anterior dorsal nucleus; AM-anterior medial nucleus; AV- anterior ventral nucleus; CC-corpus callosum; cd-caudate nucleus; cim-central intermediate nucleus; cl-central lateral nucleus; clc-central latocellular nucleus; cnMD-centromedian nucleus; csl-central superior lateral nucleus; f-fornix; fr-fasciculus retroflexus; GPi-globus pallidus, internal segment; iml-internal medullary lamina; inc-interstitial nucleus of Cajal; LD-lateral dorsal nucleus; LGN-lateral geniculate nucleus; LP-lateral posterior nucleus; lv-lateral ventricle; MDmc-medial dorsal nucleus, magnocellular division; MDmf-medial dorsal nucleus, multiform division; MDpc-medial dorsal nucleus, parvicellular division; nd-nucleus of Darkschewitsch; ot-optic tract; Pa-paraventricular nucleus; pcn-paracentral nucleus; Pf-parafascicular nucleus; pt- parataenial nucleus; ptg-posterior thalamic group; r-reticular nucleus; Re-reunions nucleus; rln-rostral linear nucleus; RNpc-red nucleus, parvicellular division; Sm-stria medullaris; SN-substantia nigra; VAmc-ventral anterior nucleus, magnocellular division; VApc-ventral anterior nucleus, parvicellular division; VLc-ventral lateral caudal nucleus; VLo-ventral lateral oral nucleus; VLps- ventral lateral postrema nucleus; VPI-ventral posterior inferior nucleus; VPLc-ventral posterior lateral caudal nucleus; VPLo- ventral posterior lateral oral nucleus; VPM-ventral posterior medial nucleus; VPMpc-ventral posterior medial nucleus, parvicellular division; zic-zona incerta. Scale bars: 5 mm applies to C-D, and G-H.

**Inline Supplementary Figure 4** - *Hypothalamus*

Two DTI/MAP parameters (*RD* and *AD*) and matched AchE and SMI-32 stained sections are particularly useful in delineating different subnuclei in the hypothalamus. The hypothalamus can be divided rostrocaudally into the rostral, middle (tuberal), and caudal groups (Suppl Fig. 4) (Rempel-Clower and Barbas, 1998; Wells et al., 2020) and has strong connections with the orbital and medial prefrontal cortex (Ongur et al., 1998). The anterior group is mostly occupied by the hyperintense preoptic area (POA), which is strikingly delineated from the hypointense internal capsule (ica), the internal segment of the globus pallidus (GPi), and the fornix (f) on *RD* image (Suppl Fig. 4A). The two small discrete regions, located above the optic chiasm (OC), the supraoptic and supra-chiasmatic nuclei (SON and SCN, respectively), also exhibited hyperintense signal intensities in *RD*. The spatial locations of SON and SCN, including medially located hypointense paraventricular nucleus (PVN) in MRI, corresponded well with the darkly stained regions in AchE and SMI-32 sections (Suppl Fig. 4A, right). The PVN also extended caudally into the middle group (see below).

The ventromedial (VM), lateral (LT), and dorsomedial (Brodmann) hypothalamic areas dominate the large part of the middle group. In addition, a small discrete infundibular (inf), arcuate (Arh), and tuberomammillary (TM) nuclei occupy the tuberal region, dorsomedial to the optic tract (ot) (Suppl Fig. 4B). The VM and DM exhibited more hyperintensity than the LT and PVN in *AD*, and the signal intensity differences between these nuclei corresponded well with the light and darkly stained neuropil in these areas in the SMI-32 section. In contrast, the neuropil of the DM is intensely stained in AchE, which clearly distinguishes this nucleus from the VM and LT hypothalamic regions (Suppl Fig. 4B, right). The neuronal cell bodies of Arh and TM are positive for AchE and SMI-32, which demarcates from the lightly stained inf nucleus (Suppl Fig. 4B, right). The Arh and inf regions exhibited more hypo-signal intensities than the TMN in *AD*.

The medial and lateral mammillary nuclei (MMN and LMN) form the caudal group of the hypothalamus (Suppl Fig. 4C). The MMN is distinguished into hyperintense medial and hypointense lateral subregions, making it the most easily identifiable structure in *RD* images. The signal intensity of LMN is comparable to the lateral part of the MMN, with no distinct border between these nuclei. In contrast, LMN and both subregions of MMN showed different staining intensities of the neuropil in AchE and SMI-32. The LMN stands out as the darkly stained region in both stained sections (Suppl Fig. 4C, middle and right). Located immediately dorsal to the MMN and LMN is the hyperintense para-mammillary nucleus. It is easily demarcated from the surrounding mammillothalamic tract (MMT), H2 field of Forel and STN with different signal contrast in MRI and staining intensities of neuropil in histology sections (Suppl Fig. 4C).

**Supplementary Figure 4.**
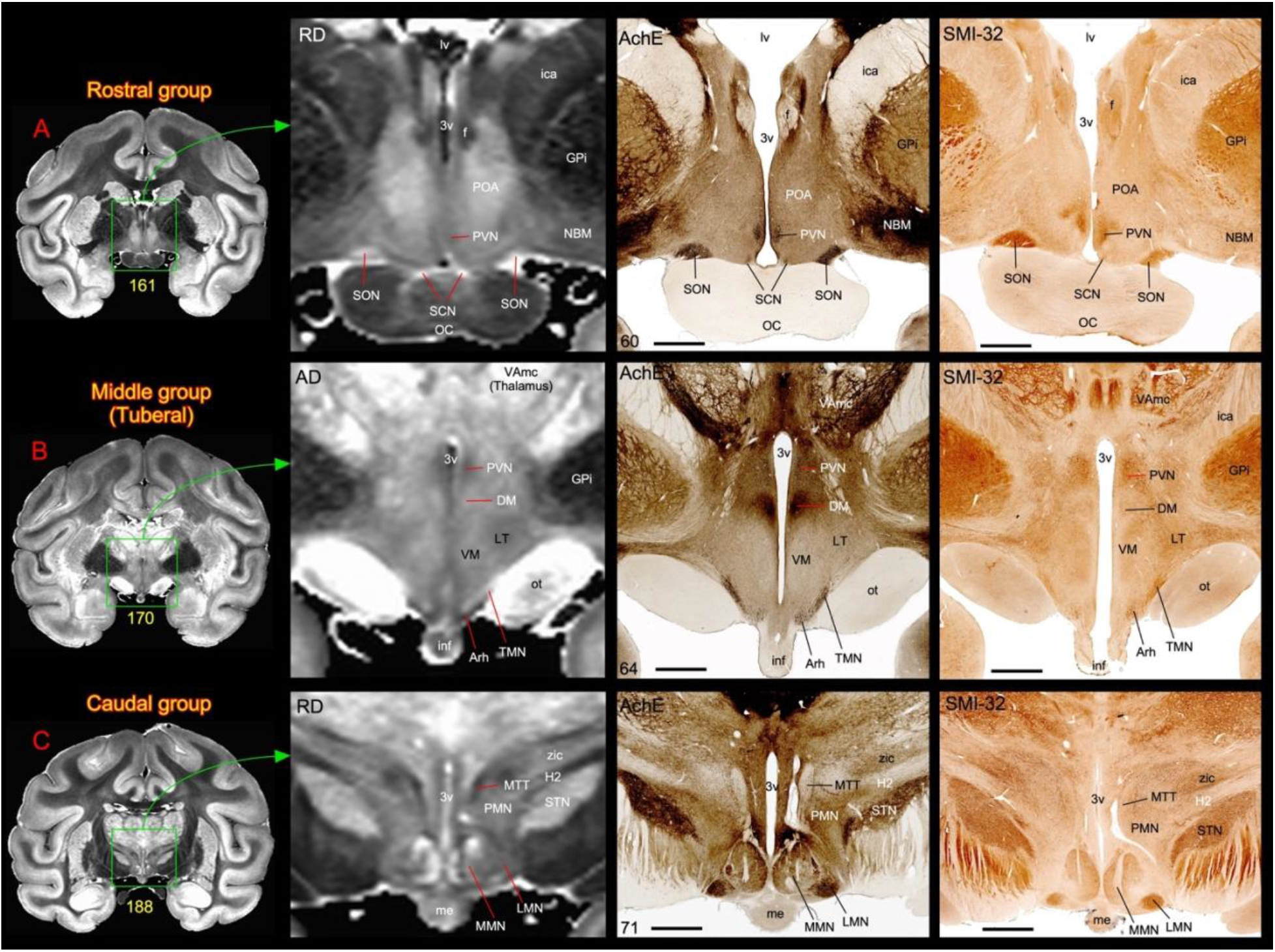
Subdivisions of the hypothalamus. **[A-C]** The zoomed-in view from coronal images (green box with arrow) shows different nuclei in the rostral, middle (or tuberal), and caudal subregions (group) of the hypothalamus in MAP-MRI (RD and AD) and corresponding histology sections stained with AchE and SMI-32. Note that the anterior group is mainly dominated by the hyperintense preoptic area (POA), which is strikingly delineated from the hypointense surrounding regions in RD image (A). Also note the subregions within the medial mammillary nucleus (MMN) with different signal intensities in the caudal group in RD image (C). For other details see the text. **Abbreviations:** 3v-3^rd^ ventricle; Arh-arcuate hypothalamic nucleus; DM-dorsomedial hypothalamic area; f-fornix; GPi-globus pallidus, internal segment; H2-H2 field of Forel; ica-anterior limb of the internal capsule; inf-infundibulum; LMN-lateral mammillary nucleus; LT-lateral hypothalamic area; lv-lateral ventricle; me-median eminence; MMN-medial mammillary nucleus; MTT-mammillothalamic tract; NBM-nucleus basalis of Meynert (Cholinergic neurons); OC-optic chiasm; ot-optic tract; PMN-paramammillary nucleus; POA-preoptic area; PVN-paraventricular hypothalamic nucleus; SCN-suprachiasmatic nucleus; SON-supraoptic nucleus; STN-subthalamic nucleus; TMN-tuberomammillary nucleus; VAmc-ventral anterior nucleus, magnocellular division; VM-ventromedial hypothalamic area; zic-zona incerta. Scale bars: 2 mm (A-C, middle and right panels).

**Inline Supplementary Figure 5 -** *Fiber bundles associated with the hippocampal formation*In the macaque monkey, the extrinsic ipsilateral and contralateral interhemispheric pathways originating from the hippocampal formation, presubiculum, and the rostrocaudal extent of cortical areas surrounding the hippocampal formation (entorhinal cortex, posterior parahippocampal gyrus, and ventral V4, respectively) course through the fornix system or the splenium of the corpus callosum (Demeter et al., 1985). The “fornix system” includes all fibers that enter the fornix from alveus (al), fimbria (fm) (Suppl Fig. 5D), and three associated interhemispheric crossing fibers that originate on one side of the fornix and decussate with the corresponding fibers of the opposite fornix (see below). As shown by the anterograde tracing method, the interhemispheric crossing fibers are segregated into three overlapping regions, oriented in the rostral to caudal axis: ventral hippocampal commissure (VHC), hippocampal decussation (HD), and dorsal hippocampal commissure (DHC), respectively, and are collectively referred to as the hippocampal commissures (Demeter et al., 1985). We identified all three commissures and other fiber bundles on direction-encoded fiber orientation distribution (DEC-FOD) images with or without reference to histology sections stained with ChAT, AchE, and PV (Suppl Fig. 5A-D). We also distinguished these crossing fibers from the adjacent body of the corpus callosum on DEC-FOD images based on the 3D fiber orientation distribution functions (*fODFs*) in different voxels, visualized with MRtrix (Tournier et al., 2012). Notably, the crossing fibers in the fornix constitute a majority of fibers at the midline caudally, whereas they represent a small fraction of fibers at the midline rostrally (Demeter et al., 1985). The VHC contains crossing fibers originating from the most anterior subdivisions of the hippocampal formation, including the genu and uncus (Demeter et al., 1985). It is located at the transition between the body and anterior columns of the fornix adjacent to the subfornical organ (SFO) and interventricular foramina (ivf) of Monro as illustrated on DEC-FOD images (Suppl Fig. 5A). Based on measurements from MR images in two cases, using the landmarks defined above, the VHC extends for approximately 1.6 mm in the rostrocaudal direction (Suppl Fig. 5E). The mediolaterally oriented crossing fibers in VHC are sharply demarcated from the rostrocaudally running fibers in the fornix on DEC-FOD images and based on *fODFs* within different voxels of VHC and fornix (Suppl Fig. 5A, arrow-Pink versus green regions). The distinction between these two fiber bundles is also confirmed in adjacent ChAT stained sections, where obliquely oriented fibers decussating across the midline are seen in VHC. The SFO appears to be devoid of fibers and is separated from VHC by a pale zone with a few scattered fibers (Suppl Fig. 5A, arrows in middle and right panels).

The DHC is a thin strip of fiber bundle located ventral to the posterior end of the body of the corpus callosum, and it extends for approximately 6.4 mm caudal to HD and rostral to the splenium of the corpus callosum (Suppl Fig. 5C, E). Despite its name, the DHC contains no fibers arising from the hippocampal formation but it contains crossing fibers originating from the presubiculum, entorhinal cortex, and the posterior parahippocampal cortex (Demeter et al., 1985, 1990; Schmahmann J and Pandya D, 2009). Unlike VHC, the DHC (as well as HD, see below) is more difficult to distinguish on DEC map alone as this ventrally located fiber bundle overlaps and is contiguous with adjacent sets of crossing fibers along the mediolateral axis within the body of the corpus callosum (Suppl Fig. 5C). To distinguish DHC from other fibers on DEC- FOD, we used matched histology sections stained with ChAT and AchE, or *fODFs*. In both stained sections, the DHC is characterized by the presence of darkly stained thin strip of fiber bundles originating from both ends of the fornix, and it stands out as a distinct layer from the adjacent lightly stained fibers in the body of the corpus callosum (Suppl Fig. 5C, right panel, plus inset).

The HD extends roughly 6.8 mm in the rostrocaudal direction and lies between the VHC and DHC and within the ventral end of the fornix (Suppl Fig. 5B, E). It carries fibers from the body of the hippocampal formation. In contrast to the fibers of the VHC and DHC, which terminate in contralateral cortical areas, these decussating fibers terminate in the contralateral septum (Demeter et al., 1985). Like DHC, the crossing fibers of HD are difficult to delineate from the rostrocaudally running fibers in the fornix on DEC-FOD images. However, the examination of matched ChAT stained section revealed a dense concentration of horizontally running fibers close to the ventral border of the fornix, and these fibers constitute the HD (Suppl Fig. 5B, middle and right). In contrast, the *fODFs* plot revealed very few crossing fibers below the fornix (see the inset in Suppl Fig. 5B, left).

The other fiber tracts, the alveus (al) and fimbria (fm) that carry subcortical and commissural fibers to and from hippocampal and parahippocampal areas are the most prominent and readily identifiable fiber bundles at the caudal level of the hippocampal formation on DEC-FOD images (Suppl Fig. 5D, F). The alvear fibers merge with the fimbrial fibers to form the posterior column of the fornix (PCF). The PCF ascends and moves medially to reach the inferior surface of the splenium of the corpus callosum (Suppl Fig. 5D, arrows) and continuous anteriorly as fornix that generates three contiguous interhemispheric hippocampal commissures. In addition, there is a dark band of fibers overlying the presubiculum (preS) on DEC-FOD, and corresponding histology sections stained with PV called the superficial presubicular pathway (SPSP) (Suppl Fig. 5D, F). Although SPSP is unrelated to fornix or hippocampal commissures, it is composed of heavily myelinated fibers that originate in the subiculum and prosubiculum and course through the molecular layer of the preS to reach the retrosplenial cortex (Rosene DL and Van- Hoesen GW, 1987).

**Supplementary Figure 5.**
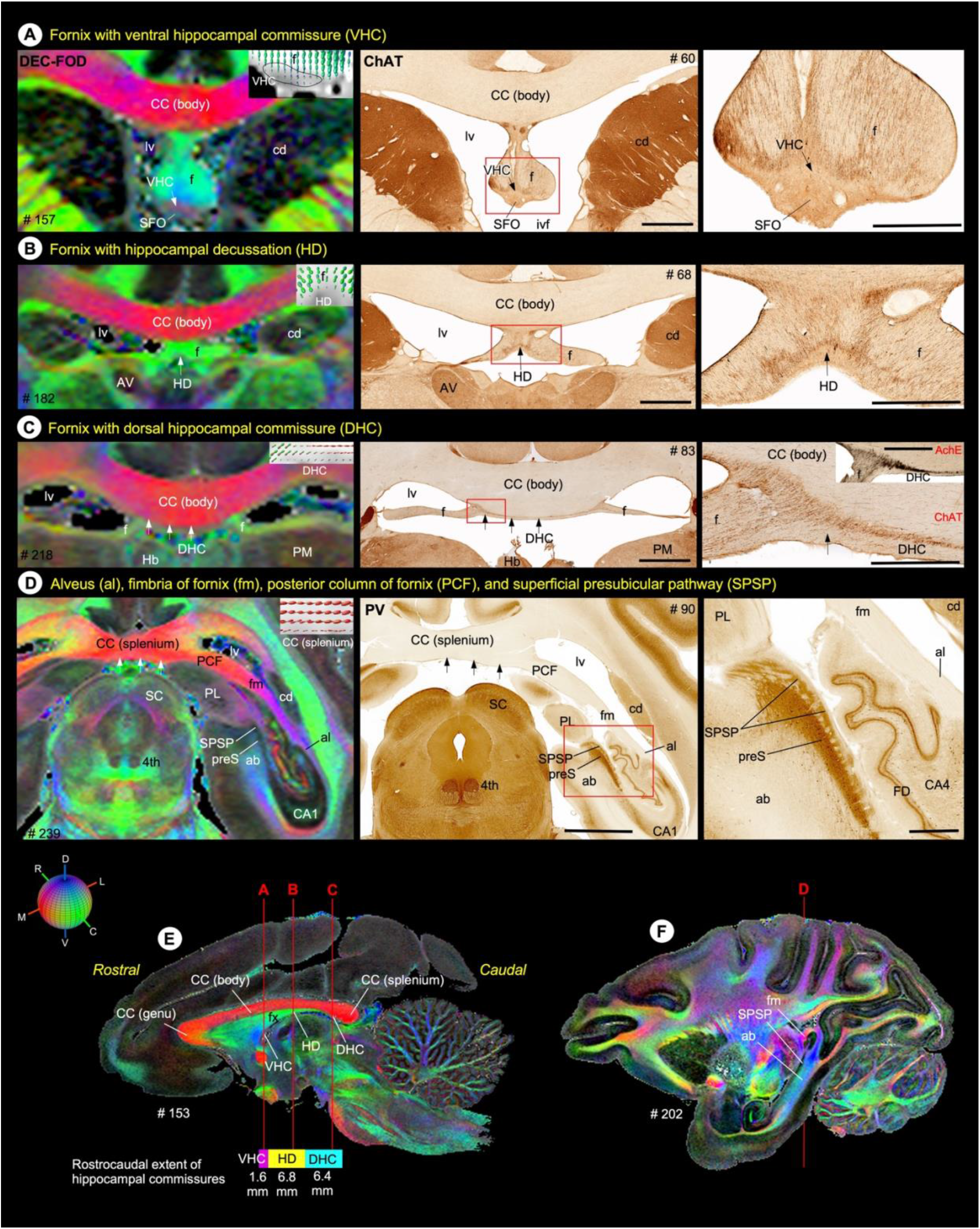
Fiber bundles of the hippocampal formation. **(A-D left column)** The spatial location of ventral hippocampal commissure (VHC), hippocampal decussation (HD), and dorsal hippocampal commissure (DHC) with the fornix, and other associated fiber bundles (alveus-al, fimbria of fornix-fm, posterior column of the fornix-pcf, and superficial presubicular pathway-SPSP) on DEC-FOD images. White arrows indicate the location of different hippocampal commissures and splenium of the corpus callosum (CC). The color-coded sphere at the bottom left illustrates the directions of the fiber bundles in DEC map; DV-dorsoventral; RC-rostrocaudal; ML-mediolateral). Inset in A-D indicates the fiber orientation distribution functions (fODFs) in different voxels at different commissures and splenium of the corpus callosum, visualized with a software package called MRtrix. The ODF inset in A-D corresponds to the regions indicated by white arrows. **(A-C, middle and right column)** Low and high-power photomicrographs of different commissures (VHC, HD, DHC; red boxed regions) in ChAT- stained sections. The high-power photomicrograph of DHC stained with AchE is also included with ChAT section as in the inset in C, right column. **(D, middle and right column)** Low and high-power photomicrographs of presubiculum (preS) and superficial presubicular pathway (SPSP) in PV-stained sections. **(E)** The rostrocaudal extent of the VHC, HD, and DHC with reference to the surrounding fiber bundles on sagittal DEC-FOD image. Three vertical lines in E indicate the corresponding location of coronal sections illustrated in A-C. **(F)** Illustrate the location of angular bundle (ab), SPSP, and fm. **Abbreviations**: 4^th^-trochlear nuclei; ab-angular bundle; al-alveus; AV-anterior ventral nucleus of thalamus; CA1 and CA4-subfields of the hippocampus; CC (body)-body of the corpus callosum; CC (splenium)-splenium of the corpus callosum; cd-caudate nucleus; DHC-dorsal hippocampal commissure; f-fornix; FD-fascia dentata; fm-fimbria of fornix; Hb-habenular nucleus; HD- hippocampal decussation; lv-lateral ventricle; PCF-posterior column of fornix; PL-lateral pulvinar; PM-medial pulvinar; preS- presubiculum; SC-superior colliculus; SPSP-superficial presubicular pathway; VHC-ventral hippocampal commissure. Scale bars: 2 mm (A-C middle column), 5 mm (D middle column), 1 mm (A-C right column), and 1 mm (D right column).

**Inline Supplementary Figure 6** *Subcortical regions in ex vivo and in vivo MRI.* The closely matched sagittal MR slices from *ex vivo* MAP-MRI (**A**, current study), *ex vivo* MTR (**B**), population-averaged *in vivo* T1w (**C**), and standard *in vivo* T1w MRI (**D**) volumes show the selected brainstem nuclei and fiber tracts. Note that the nuclei (dark gray regions) are sharply delineated from the surrounding fiber bundles of different orientations, identified on the MAP-MRI with directional information (*fODF*s/DEC). In contrast, the delineations of nuclei from the surrounding white matter pathways are less prominent (B) or barely visible (C-D) in other MRIs. ***Abbreviations***: cd-caudate nucleus; CST-corticospinal tract; ion-inferior olivary nucleus; MD-mediodorsal nucleus; ml-medial lemniscus; NA-nucleus accumbens; pcf-pontocerebellar fibers; PN-pontine nuclei; SN-substantia nigra; vn- vestibular nuclei.

**Supplementary Figure 6.**
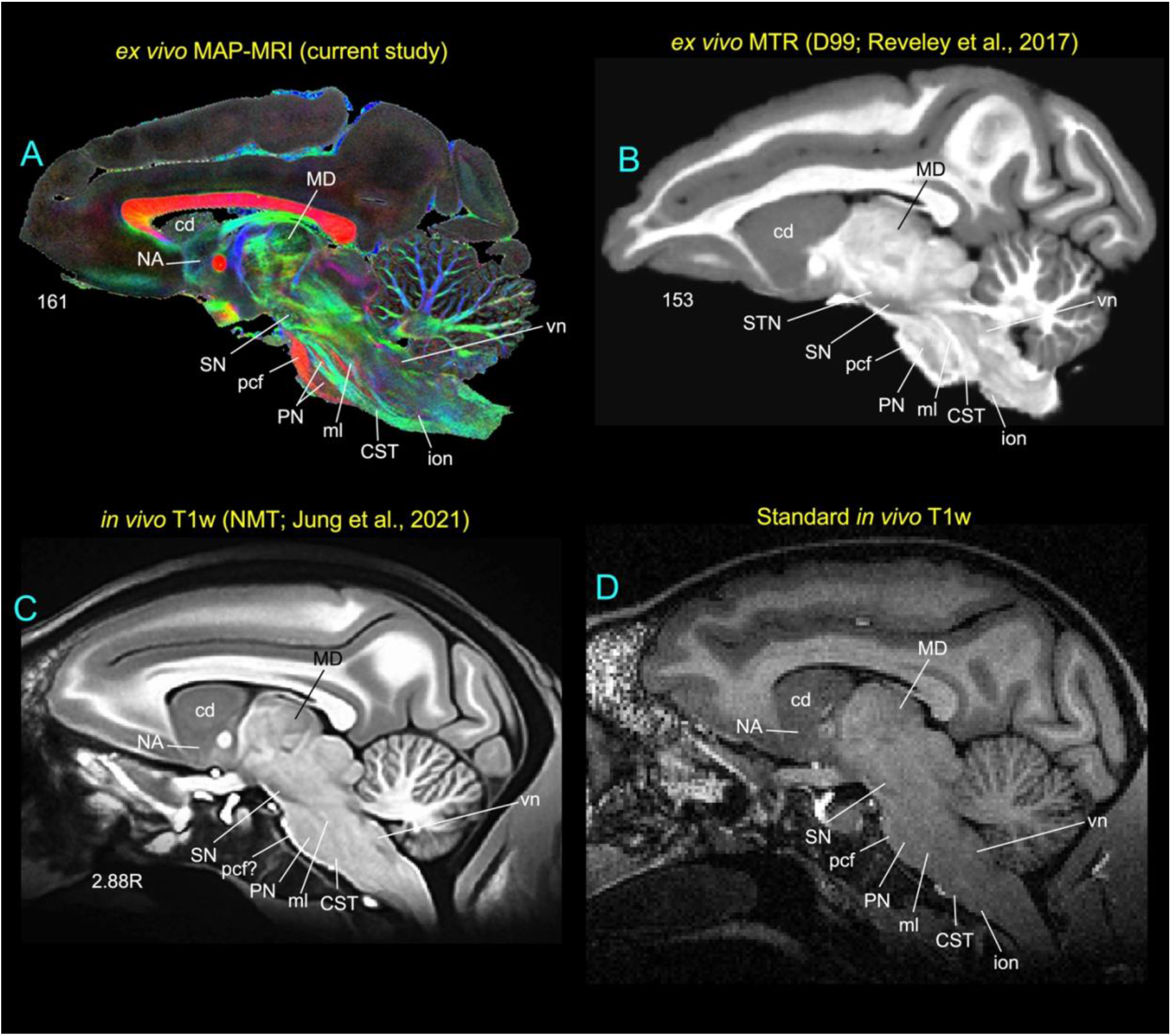
Subcortical regions in in vivo and ex vivo structural MRI.

## Notes

### Competing Interest Statement

The authors have declared no competing interest.

## References

Abosch, A., Yacoub, E., Ugurbil, K., Harel, N., 2010. An assessment of current brain targets for deep brain stimulation surgery with susceptibility-weighted imaging at 7 tesla. Neurosurgery 67, 1745–1756; discussion 1756.

Aggleton, J.P., O’Mara, S.M., Vann, S.D., Wright, N.F., Tsanov, M., Erichsen, J.T., 2010. Hippocampal-anterior thalamic pathways for memory: uncovering a network of direct and indirect actions. Eur J Neurosci 31, 2292–2307.

Amaral, D.G., Bassett, J.L., 1989. Cholinergic innervation of the monkey amygdala: an immunohistochemical analysis with antisera to choline acetyltransferase. J Comp Neurol 281, 337–361.

Arsenault, M.Y., Parent, A., Seguela, P., Descarries, L., 1988. Distribution and morphological characteristics of dopamine-immunoreactive neurons in the midbrain of the squirrel monkey (Saimiri sciureus). J Comp Neurol 267, 489–506.

Asanuma, C., Thach, W.T., Jones, E.G., 1983. Distribution of cerebellar terminations and their relation to other afferent terminations in the ventral lateral thalamic region of the monkey. Brain Res 286, 237–265.

Avants, B.B., Tustison, N., Song, G., 2009. Advanced normalization tools (ANTS). Insight j 2, 1–35.

Avram, A.V., Barnett, A.S., Basser, P.J., 2014a. The variation of MAP-MRI derived parameters along white matter fiber pathways in the human brain. Proceedings of the International Society for Magnetic Resonance in Medicine, Milan, Italy, p. 2587.

Avram, A.V., Bernstein, A.S., Irfanoglu, M.O., Simmons, A., Cota, M., Gai, N., Jikaria, A., Moses, A., Turtzo, C., Latour, L., Pham, D., Butman, J.A., Basser, P.J., 2018a. Anatomical Atlas of 3D MAP MRI-derived 3D diffusion propagators and microstructural parameters. Proceedings of the International Society for Magnetic Resonance in Medicine, Paris, France, p. 1577.

Avram, A.V., Guidon, A., Song, A.W., 2010. Myelin water weighted diffusion tensor imaging. Neuroimage 53, 132–138.

Avram, A.V., Guidon, A., Truong, T.-K., Liu, C., Song, A.W., 2014b. Dynamic and inherent B0 correction for DTI using stimulated echo spiral imaging. Magnetic Resonance in Medicine 71, 1044–1053.

Avram, A.V., Hutchinson, E., Basser, P.J., 2017. Higher-order statistics of 3D spin displacement probability distributions measured with MAP MRI. Proceedings of the International Society for Magnetic Resonance in Medicine, Hawai’i, USA, p. 3367.

Avram, A.V., Saleem, K.S., Ye, F.Q., Chen, C.C., Komlosh, M.E., Basser, P.J., 2020a. Whole-brain mapping of cortical architectonic features with high-resolution MAP-MRI. Proceedings of the International Society for Magnetic Resonance in Medicine, Virtucal Conference, p. 740.

Avram, A.V., Saleem, K.S., Ye, F.Q., Yen, C.C., Komlosh, M.E., Basser, P.J., 2020b. Modeling cortical architectonic features by analyzing diffusion MRI data in the cortical reference frame. Proceedings of the International Society for Magnetic Resonance in Medicine Virtual Conference, Virtual Conference, p. 713.

Avram, A.V., Sarlls, J.E., Barnett, A.S., Ozarslan, E., Thomas, C., Irfanoglu, M.O., Hutchinson, E., Pierpaoli, C., Basser, P.J., 2016. Clinical feasibility of using mean apparent propagator (MAP) MRI to characterize brain tissue microstructure. Neuroimage 127, 422–434.

Avram, A.V., Sarlls, J.E., Basser, P.J., 2019. Measuring non-parametric distributions of intravoxel mean diffusivities using a clinical MRI scanner. Neuroimage 185, 255–262.

Avram, A.V., Sarlls, J.E., Basser, P.J., 2021. Whole-Brain Imaging of Subvoxel T1-Diffusion Correlation Spectra in Human Subjects. Frontiers in Neuroscience 15.

Avram, A.V., Sarlls, J.E., Hutchinson, E., Basser, P.J., 2018b. Efficient experimental designs for isotropic generalized diffusion tensor MRI (IGDTI). Magnetic Resonance in Medicine 79, 180–194.

Baker, J.T., Patel, G.H., Corbetta, M., Snyder, L.H., 2006. Distribution of activity across the monkey cerebral cortical surface, thalamus and midbrain during rapid, visually guided saccades. Cereb Cortex 16, 447–459.

Baron, M.S., Sidibe, M., DeLong, M.R., Smith, Y., 2001. Course of motor and associative pallidothalamic projections in monkeys. J Comp Neurol 429, 490–501.

Basser, P.J., 1995. Inferring microstructural features and the physiological state of tissues from diffusion-weighted images. NMR Biomed 8, 333–344.

Basser, P.J., Mattiello, J., LeBihan, D., 1994. MR diffusion tensor spectroscopy and imaging. Biophys J 66, 259–267.

Boscolo Galazzo, I., Brusini, L., Akinci, M., Cruciani, F., Pitteri, M., Ziccardi, S., Bajrami, A., Castellaro, M., Salih, A.M.A., Pizzini, F.B., Jovicich, J., Calabrese, M., Menegaz, G., Unraveling the MRI-Based Microstructural Signatures Behind Primary Progressive and Relapsing–Remitting Multiple Sclerosis Phenotypes. Journal of Magnetic Resonance Imaging n/a.

Boscolo Galazzo, I., Brusini, L., Obertino, S., Zucchelli, M., Granziera, C., Menegaz, G., 2018. On the viability of diffusion MRI-based microstructural biomarkers in ischemic stroke. Frontiers in Neuroscience 12, 92.

Brimblecombe, K.R., Cragg, S.J., 2017. The Striosome and Matrix Compartments of the Striatum: A Path through the Labyrinth from Neurochemistry toward Function. ACS Chem Neurosci 8, 235–242.

Brodmann, K., 1909. Vergleichende Lokalisationslehre der Grosshirnrinde. Johann Ambrosius Barth, Leipzig.

Calabrese, E., Badea, A., Coe, C.L., Lubach, G.R., Shi, Y., Styner, M.A., Johnson, G.A., 2015a. A diffusion tensor MRI atlas of the postmortem rhesus macaque brain. Neuroimage 117, 408–416.

Calabrese, E., Hickey, P., Hulette, C., Zhang, J., Parente, B., Lad, S.P., Johnson, G.A., 2015b. Postmortem diffusion MRI of the human brainstem and thalamus for deep brain stimulator electrode localization. Hum Brain Mapp 36, 3167–3178.

Carmichael, S.T., Price, J.L., 1994. Architectonic subdivision of the orbital and medial prefrontal cortex in the macaque monkey. J Comp Neurol 346, 366–402.

Coude, D., Parent, A., Parent, M., 2018. Single-axon tracing of the corticosubthalamic hyperdirect pathway in primates. Brain Struct Funct 223, 3959–3973.

Cox, R.W., 1996. AFNI: software for analysis and visualization of functional magnetic resonance neuroimages. Comput Biomed Res 29, 162–173.

Crittenden, J.R., Graybiel, A.M., 2011. Basal Ganglia disorders associated with imbalances in the striatal striosome and matrix compartments. Front Neuroanat 5, 59.

DeArmond, S.J., Fusco, M.M., Dewey, M.M., 1989. Structure of the human brain. A photographic atlas., 3 ed. Oxford University Press, Oxford/New York.

Deistung, A., Schafer, A., Schweser, F., Biedermann, U., Gullmar, D., Trampel, R., Turner, R., Reichenbach, J.R., 2013a. High-Resolution MR Imaging of the Human Brainstem In vivo at 7 Tesla. Front Hum Neurosci 7, 710.

Deistung, A., Schafer, A., Schweser, F., Biedermann, U., Turner, R., Reichenbach, J.R., 2013b. Toward in vivo histology: a comparison of quantitative susceptibility mapping (QSM) with magnitude-, phase-, and R2*-imaging at ultra-high magnetic field strength. Neuroimage 65, 299–314.

Demeter, S., Rosene, D.L., Van Hoesen, G.W., 1985. Interhemispheric pathways of the hippocampal formation, presubiculum, and entorhinal and posterior parahippocampal cortices in the rhesus monkey: the structure and organization of the hippocampal commissures. J Comp Neurol 233, 30–47.

Demeter, S., Rosene, D.L., Van Hoesen, G.W., 1990. Fields of origin and pathways of the interhemispheric commissures in the temporal lobe of macaques. J Comp Neurol 302, 29–53.

Diaz, E., Bravo, D., Rojas, X., Concha, M.L., 2011. Morphologic and immunohistochemical organization of the human habenular complex. J Comp Neurol 519, 3727–3747.

Eblen, F., Graybiel, A.M., 1995. Highly restricted origin of prefrontal cortical inputs to striosomes in the macaque monkey. J Neurosci 15, 5999–6013.

Ewert, S., Plettig, P., Li, N., Chakravarty, M.M., Collins, D.L., Herrington, T.M., Kuhn, A.A., Horn, A., 2018. Toward defining deep brain stimulation targets in MNI space: A subcortical atlas based on multimodal MRI, histology and structural connectivity. Neuroimage 170, 271–282.

Feinberg, D.A., Moeller, S., Smith, S.M., Auerbach, E., Ramanna, S., Glasser, M.F., Miller, K.L., Ugurbil, K., Yacoub, E., 2010. Multiplexed Echo Planar Imaging for Sub-Second Whole Brain FMRI and Fast Diffusion Imaging. PLoS One 5, e15710.

Folloni, D., Verhagen, L., Mars, R.B., Fouragnan, E., Constans, C., Aubry, J.F., Rushworth, M.F.S., Sallet, J., 2019. Manipulation of Subcortical and Deep Cortical Activity in the Primate Brain Using Transcranial Focused Ultrasound Stimulation. Neuron 101, 1109–1116 e1105.

Francois, C., Percheron, G., Yelnik, J., Heyner, S., 1985. A Histological Atlas of the Macaque (Macaca mulatta) Substantia Nigra in Ventricular Coordinate. Brain Res Bulletin 14, 349–367.

Galvan, A., Hu, X., Smith, Y., Wichmann, T., 2012. In vivo optogenetic control of striatal and thalamic neurons in non-human primates. PLoS One 7, e50808.

Gerfen, C.R., 1989. The neostriatal mosaic: striatal patch-matrix organization is related to cortical lamination. Science 246, 385–388.

Goldstein, M.E., Sternberger, L.A., Sternberger, N.H., 1987. Varying degrees of phosphorylation determine microheterogeneity of the heavy neurofilament polypeptide (Nf-H). J Neuroimmunol 14, 135–148.

Graybiel, A.M., Ragsdale, C.W., Jr., 1978. Histochemically distinct compartments in the striatum of human, monkeys, and cat demonstrated by acetylthiocholinesterase staining. Proc Natl Acad Sci U S A 75, 5723–5726.

Haber, S.N., 2003. The primate basal ganglia: parallel and integrative networks. J Chem Neuroanat 26, 317–330.

Halassa, M.M., Kastner, S., 2017. Thalamic functions in distributed cognitive control. Nat Neurosci 20, 1669–1679.

Hartig, R., Glen, D., Jung, B., Logothetis, N.K., Paxinos, G., Garza-Villarreal, E.A., Messinger, A., Evrard, H.C., 2021. The Subcortical Atlas of the Rhesus Macaque (SARM) for neuroimaging. Neuroimage 235, 117996.

Haynes, W.I., Haber, S.N., 2013. The organization of prefrontal-subthalamic inputs in primates provides an anatomical substrate for both functional specificity and integration: implications for Basal Ganglia models and deep brain stimulation. J Neurosci 33, 4804–4814.

Heimer, L., Switzer, R.D., Van Hoesen, G.W., 1982. Ventral striatum and ventral pallidum: Components of the motor system? TINS 5, 83–87.

Hikosaka, O., Sesack, S.R., Lecourtier, L., Shepard, P.D., 2008. Habenula: crossroad between the basal ganglia and the limbic system. J Neurosci 28, 11825–11829.

Hirsch, E.C., Graybiel, A.M., Hersh, L.B., Duyckaerts, C., Agid, Y., 1989. Striosomes and extrastriosomal matrix contain different amounts of immunoreactive choline acetyltransferase in the human striatum. Neurosci Lett 96, 145–150.

Hoch, M.J., Bruno, M.T., Faustin, A., Cruz, N., Crandall, L., Wisniewski, T., Devinsky, O., Shepherd, T.M., 2019a. 3T MRI Whole-Brain Microscopy Discrimination of Subcortical Anatomy, Part 1: Brain Stem. AJNR Am J Neuroradiol 40, 401–407.

Hoch, M.J., Bruno, M.T., Faustin, A., Cruz, N., Mogilner, A.Y., Crandall, L., Wisniewski, T., Devinsky, O., Shepherd, T.M., 2019b. 3T MRI Whole-Brain Microscopy Discrimination of Subcortical Anatomy, Part 2: Basal Forebrain. AJNR Am J Neuroradiol 40, 1095–1105.

Hof, P.R., Cox, K., Morrison, J.H., 1990. Quantitative analysis of a vulnerable subset of pyramidal neurons in Alzheimer’s disease: I. Superior frontal and inferior temporal cortex. J Comp Neurol 301, 44–54.

Hof, P.R., Glezer, II, Revishchin, A.V., Bouras, C., Charnay, Y., Morgane, P.J., 1995. Distribution of dopaminergic fibers and neurons in visual and auditory cortices of the harbor porpoise and pilot whale. Brain Res Bull 36, 275–284.

Hof, P.R., Morrison, J.H., 1990. Quantitative analysis of a vulnerable subset of pyramidal neurons in Alzheimer’s disease: II. Primary and secondary visual cortex. J Comp Neurol 301, 55–64.

Hof, P.R., Morrison, J.H., 1995. Neurofilament protein defines regional patterns of cortical organization in the macaque monkey visual system: a quantitative immunohistochemical analysis. J Comp Neurol 352, 161–186.

Horn, A.K.E., Horng, A., Buresch, N., Messoudi, A., Hartig, W., 2018. Identification of Functional Cell Groups in the Abducens Nucleus of Monkey and Human by Perineuronal Nets and Choline Acetyltransferase Immunolabeling. Front Neuroanat 12, 45.

Hsu, S.M., Raine, L., Fanger, H., 1981. Use of avidin-biotin-peroxidase complex (ABC) in immunoperoxidase techniques: a comparison between ABC and unlabeled antibody (PAP) procedures. J Histochem Cytochem 29, 577–580.

Huang, S.Y., Witzel, T., Keil, B., Scholz, A., Davids, M., Dietz, P., Rummert, E., Ramb, R., Kirsch, J.E., Yendiki, A., Fan, Q., Tian, Q., Ramos-Llorden, G., Lee, H.H., Nummenmaa, A., Bilgic, B., Setsompop, K., Wang, F., Avram, A.V., Komlosh, M., Benjamini, D., Magdoom, K.N., Pathak, S., Schneider, W., Novikov, D.S., Fieremans, E., Tounekti, S., Mekkaoui, C., Augustinack, J., Berger, D., Shapson-Coe, A., Lichtman, J., Basser, P.J., Wald, L.L., Rosen, B.R., 2021. Connectome 2.0: Developing the next-generation ultra-high gradient strength human MRI scanner for bridging studies of the micro-, meso- and macro-connectome. Neuroimage 243, 118530.

Hung, C.C., Yen, C.C., Ciuchta, J.L., Papoti, D., Bock, N.A., Leopold, D.A., Silva, A.C., 2015. Functional MRI of visual responses in the awake, behaving marmoset. Neuroimage 120, 1–11.

Hutchinson, E.B., Schwerin, S.C., Avram, A.V., Juliano, S.L., Pierpaoli, C., 2018. Diffusion MRI and the detection of alterations following traumatic brain injury. Journal of Neuroscience Research 96, 612–625.

Jankowski, M.M., Ronnqvist, K.C., Tsanov, M., Vann, S.D., Wright, N.F., Erichsen, J.T., Aggleton, J.P., O’Mara, S.M., 2013. The anterior thalamus provides a subcortical circuit supporting memory and spatial navigation. Front Syst Neurosci 7, 45.

Jiang, R., Jiang, S., Song, S., Wei, X., Deng, K., Zhang, Z., Xue, Y., 2021. Laplacian-regularized mean apparent propagator-MRI in evaluating corticospinal tract injury in patients with brain glioma. Korean Journal of Radiology 22, 759.

Jimenez-Castellanos, J., Graybiel, A.M., 1987. Subdivisions of the primate substantia nigra pars compacta detected by acetylcholinesterase histochemisty. Brain Res 437, 349–354.

Johnson, V.E., Stewart, W., Weber, M.T., Cullen, D.K., Siman, R., Smith, D.H., 2016. SNTF immunostaining reveals previously undetected axonal pathology in traumatic brain injury. Acta Neuropathol 131, 115–135.

Jones, E.G., 1998. The thalamus of primates. In: Handbook of Chemical Neuroanatomy: The primate nervous system. Part II. Elsevier, New York.

Jones, E.G., Hendry, S.H., 1989. Differential Calcium Binding Protein Immunoreactivity Distinguishes Classes of Relay Neurons in Monkey Thalamic Nuclei. Eur J Neurosci 1, 222–246.

Jung, B., Taylor, P.A., Seidlitz, J., Sponheim, C., Perkins, P., Ungerleider, L.G., Glen, D., Messinger, A., 2021. A comprehensive macaque fMRI pipeline and hierarchical atlas. Neuroimage 235, 117997.

Kaas, J.H., 2012. Somatosensory system. In: Mai, J.K., Paxinos, G. (Eds.), The human nervous system. Academic Press, New York, pp. 1074–1109.

Keil, B., Blau, J.N., Biber, S., Hoecht, P., Tountcheva, V., Setsompop, K., Triantafyllou, C., Wald, L.L., 2013. A 64-channel 3T array coil for accelerated brain MRI. Magnetic Resonance in Medicine 70, 248–258.

Keuken, M.C., Bazin, P.L., Crown, L., Hootsmans, J., Laufer, A., Muller-Axt, C., Sier, R., van der Putten, E.J., Schafer, A., Turner, R., Forstmann, B.U., 2014. Quantifying inter-individual anatomical variability in the subcortex using 7 T structural MRI. Neuroimage 94, 40–46.

Koay, C.G., Özarslan, E., Johnson, K.M., Meyerand, M.E., 2012. Sparse and optimal acquisition design for diffusion MRI and beyond. Medical Physics 39, 2499–2511.

Kondo, H., Lavenex, P., Amaral, D.G., 2008. Intrinsic connections of the macaque monkey hippocampal formation: I. Dentate gyrus. J Comp Neurol 511, 497–520.

Kondo, H., Lavenex, P., Amaral, D.G., 2009. Intrinsic connections of the macaque monkey hippocampal formation: II. CA3 connections. J Comp Neurol 515, 349–377.

Kundu, B., Brock, A.A., Englot, D.J., Butson, C.R., Rolston, J.D., 2018. Deep brain stimulation for the treatment of disorders of consciousness and cognition in traumatic brain injury patients: a review. Neurosurg Focus 45, E14.

Lanciego, J.L., Luquin, N., Obeso, J.A., 2012. Functional neuroanatomy of the basal ganglia. Cold Spring Harb Perspect Med 2, a009621.

Larsell, O., 1953. The cerebellum of the cat and the monkey. J Comp Neurol 99, 135–199.

Le, H., Zeng, W., Zhang, H., Li, J., Wu, X., Xie, M., Yan, X., Zhou, M., Zhang, H., Wang, M., Hong, G., Shen, J., 2020. Mean Apparent Propagator MRI Is Better Than Conventional Diffusion Tensor Imaging for the Evaluation of Parkinson’s Disease: A Prospective Pilot Study. Frontiers in Aging Neuroscience 12.

Lenglet, C., Abosch, A., Yacoub, E., De Martino, F., Sapiro, G., Harel, N., 2012. Comprehensive in vivo mapping of the human basal ganglia and thalamic connectome in individuals using 7T MRI. PLoS One 7, e29153.

Logothetis, N.K., Eschenko, O., Murayama, Y., Augath, M., Steudel, T., Evrard, H.C., Besserve, M., Oeltermann, A., 2012. Hippocampal-cortical interaction during periods of subcortical silence. Nature 491, 547–553.

Ma, K., Zhang, X., Zhang, H., Yan, X., Gao, A., Song, C., Wang, S., Lian, Y., Cheng, J., 2020. Mean apparent propagator-MRI: a new diffusion model which improves temporal lobe epilepsy lateralization. European journal of radiology 126, 108914.

Mallet, L., Schupbach, M., N’Diaye, K., Remy, P., Bardinet, E., Czernecki, V., Welter, M.L., Pelissolo, A., Ruberg, M., Agid, Y., Yelnik, J., 2007. Stimulation of subterritories of the subthalamic nucleus reveals its role in the integration of the emotional and motor aspects of behavior. Proc Natl Acad Sci U S A 104, 10661–10666.

Martin, L.J., Blackstone, C.D., Huganir, R.L., Price, D.L., 1993. The striatal mosaic in primates: striosomes and matrix are differentially enriched in ionotropic glutamate receptor subunits. J Neurosci 13, 782–792.

Massey, L.A., Miranda, M.A., Zrinzo, L., Al-Helli, O., Parkes, H.G., Thornton, J.S., So, P.W., White, M.J., Mancini, L., Strand, C., Holton, J.L., Hariz, M.I., Lees, A.J., Revesz, T., Yousry, T.A., 2012. High resolution MR anatomy of the subthalamic nucleus: imaging at 9.4 T with histological validation. Neuroimage 59, 2035–2044.

Matsui, T., Koyano, K.W., Tamura, K., Osada, T., Adachi, Y., Miyamoto, K., Chikazoe, J., Kamigaki, T., Miyashita, Y., 2012. FMRI activity in the macaque cerebellum evoked by intracortical microstimulation of the primary somatosensory cortex: evidence for polysynaptic propagation. PLoS One 7, e47515.

McNab, J.A., Edlow, B.L., Witzel, T., Huang, S.Y., Bhat, H., Heberlein, K., Feiweier, T., Liu, K., Keil, B., Cohen-Adad, J., 2013. The Human Connectome Project and beyond: initial applications of 300 mT/m gradients. Neuroimage 80, 234–245.

Min, H.K., Ross, E.K., Jo, H.J., Cho, S., Settell, M.L., Jeong, J.H., Duffy, P.S., Chang, S.Y., Bennet, K.E., Blaha, C.D., Lee, K.H., 2016. Dopamine Release in the Nonhuman Primate Caudate and Putamen Depends upon Site of Stimulation in the Subthalamic Nucleus. J Neurosci 36, 6022–6029.

Mitchell, A.S., Sherman, S.M., Sommer, M.A., Mair, R.G., Vertes, R.P., Chudasama, Y., 2014. Advances in understanding mechanisms of thalamic relays in cognition and behavior. J Neurosci 34, 15340–15346.

Murris, S.R., Arsenault, J.T., Vanduffel, W., 2020. Frequency- and State-Dependent Network Effects of Electrical Stimulation Targeting the Ventral Tegmental Area in Macaques. Cereb Cortex 30, 4281–4296.

Naidich, T.P., Duvernoy, H.M., Delman, B.N., Sorensen, A.G., Kollias, S.S., Haacke, E.M., 2009. Duvernoy’s Atlas of the Human Brain Stem and Cerebellum. High-Field MRI: Surface anatomy, internal structure, vascularit. Springer-Wien, New York.

Naik, N.T., 1963. Technical variations in Koelle’s histochemical method for demonstrating cholinesterase activity. Q J MICROSC SCI 104, 89–100.

Nambu, A., Takada, M., Inase, M., Tokuno, H., 1996. Dual somatotopical representations in the primate subthalamic nucleus: evidence for ordered but reversed body-map transformations from the primary motor cortex and the supplementary motor area. J Neurosci 16, 2671–2683.

Neudorfer, C., El Majdoub, F., Hunsche, S., Richter, K., Sturm, V., Maarouf, M., 2017. Deep Brain Stimulation of the H Fields of Forel Alleviates Tics in Tourette Syndrome. Front Hum Neurosci 11, 308.

Neudorfer, C., Maarouf, M., 2018. Neuroanatomical background and functional considerations for stereotactic interventions in the H fields of Forel. Brain Struct Funct 223, 17–30.

Oishi, K., Mori, S., Troncoso, J.C., Lenz, F.A., 2020. Mapping tracts in the human subthalamic area by 11.7T ex vivo diffusion tensor imaging. Brain Struct Funct 225, 1293–1312.

Olszewski, J., 1952. The thalamus of the Macaca mulatta: An atlas for use with the stereotaxic instrument. S. Karger, Basel., New York.

Ongur, D., An, X., Price, J.L., 1998. Prefrontal cortical projections to the hypothalamus in macaque monkeys. J Comp Neurol 401, 480–505.

Ortiz-Rios, M., Kusmierek, P., DeWitt, I., Archakov, D., Azevedo, F.A., Sams, M., Jaaskelainen, I.P., Keliris, G.A., Rauschecker, J.P., 2015. Functional MRI of the vocalization-processing network in the macaque brain. Front Neurosci 9, 113.

Ouhaz, Z., Fleming, H., Mitchell, A.S., 2018. Cognitive Functions and Neurodevelopmental Disorders Involving the Prefrontal Cortex and Mediodorsal Thalamus. Front Neurosci 12, 33.

Ozarslan, E., Koay, C.G., Shepherd, T.M., Komlosh, M.E., Irfanoglu, M.O., Pierpaoli, C., Basser, P.J., 2013. Mean apparent propagator (MAP) MRI: a novel diffusion imaging method for mapping tissue microstructure. Neuroimage 78, 16–32.

Pajevic, S., Pierpaoli, C., 1999. Color schemes to represent the orientation of anisotropic tissues from diffusion tensor data: application to white matter fiber tract mapping in the human brain. Magn Reson Med 42, 526–540.

Palomero-Gallagher, N., Kedo, O., Mohlberg, H., Zilles, K., Amunts, K., 2020. Multimodal mapping and analysis of the cyto- and receptorarchitecture of the human hippocampus. Brain Struct Funct 225, 881–907.

Parent, A., 1990. Extrinsic connections of the basal ganglia. Trends Neurosci 13, 254–258.

Parent, A., Hazrati, L.N., 1995. Functional anatomy of the basal ganglia. II. The place of subthalamic nucleus and external pallidum in basal ganglia circuitry. Brain Res Brain Res Rev 20, 128–154.

Parent, M., Levesque, M., Parent, A., 2001. Two types of projection neurons in the internal pallidum of primates: single-axon tracing and three-dimensional reconstruction. J Comp Neurol 439, 162–175.

Parent, M., Parent, A., 2004. The pallidofugal motor fiber system in primates. Parkinsonism Relat Disord 10, 203–211.

Pauli, W.M., Nili, A.N., Tyszka, J.M., 2018. A high-resolution probabilistic in vivo atlas of human subcortical brain nuclei. Sci Data 5, 180063.

Paxinos, G., Huang, X.F., Petrides, M., Toga, A.W., 2009. The Rhesus Monkey Brain in Stereotaxic Coordinates, 2 ed. Elsevier/Academic press, San Diego.

Pergola, G., Danet, L., Pitel, A.L., Carlesimo, G.A., Segobin, S., Pariente, J., Suchan, B., Mitchell, A.S., Barbeau, E.J., 2018. The Regulatory Role of the Human Mediodorsal Thalamus. Trends Cogn Sci 22, 1011–1025.

Pierpaoli, C., Basser, P.J., 1996. Toward a quantitative assessment of diffusion anisotropy. Magn Reson Med 36, 893–906.

Pierpaoli, C., Jezzard, P., Basser, P.J., Barnett, A., Di Chiro, G., 1996. Diffusion tensor MR imaging of the human brain. Radiology 201, 637–648.

Pierpaoli, C., Walker, L., Irfanoglu, M.O., Barnett, A., Basser, P.J., Chang, L.-C., Koay, C.G., Pajevic, S., Rohde, G., Sarlls, J.E., Wu, M., 2010. TORTOISE: an integrated software package for processing of diffusion MRI data., International Society for Magnetic Resonance in Medicine (ISMRM).

Pitkanen, A., Amaral, D.G., 1998. Organization of the intrinsic connections of the monkey amygdaloid complex: projections originating in the lateral nucleus. J Comp Neurol 398, 431–458.

Plantinga, B.R., Temel, Y., Roebroeck, A., Uludag, K., Ivanov, D., Kuijf, M.L., Ter Haar Romenij, B.M., 2014. Ultra-high field magnetic resonance imaging of the basal ganglia and related structures. Front Hum Neurosci 8, 876.

Price, J.L., Russchen, F.T., Amaral, D.G., 1987. The limbic region. II. The amygdaloid complex. In: Bjorkland, A., Hokfelt, T., Swanson, L. (Eds.), Handbook of Chemical Neuroanatomy. Elsevier, Amsterdam.

Rempel-Clower, N.L., Barbas, H., 1998. Topographic organization of connections between the hypothalamus and prefrontal cortex in the rhesus monkey. J Comp Neurol 398, 393–419.

Reveley, C., Gruslys, A., Ye, F.Q., Glen, D., Samaha, J., B, E.R., Saad, Z., A, K.S., Leopold, D.A., Saleem, K.S., 2017. Three-Dimensional Digital Template Atlas of the Macaque Brain. Cereb Cortex 27, 4463–4477.

Rijkers, K., Temel, Y., Visser-Vandewalle, V., Vanormelingen, L., Vandersteen, M., Adriaensens, P., Gelan, J., Beuls, E.A., 2007. The microanatomical environment of the subthalamic nucleus. Technical note. J Neurosurg 107, 198–201.

Rohlfing, T., Kroenke, C.D., Sullivan, E.V., Dubach, M.F., Bowden, D.M., Grant, K.A., Pfefferbaum, A., 2012. The INIA19 Template and NeuroMaps Atlas for Primate Brain Image Parcellation and Spatial Normalization. Front Neuroinform 6, 27.

Roman, E., Weininger, J., Lim, B., Roman, M., Barry, D., Tierney, P., O’Hanlon, E., Levins, K., O’Keane, V., Roddy, D., 2020. Untangling the dorsal diencephalic conduction system: a review of structure and function of the stria medullaris, habenula and fasciculus retroflexus. Brain Struct Funct 225, 1437–1458.

Rosene DL, Van-Hoesen GW, 1987. The hippocampal formation of the primate brain; A review of some comparative aspects of cytoarchitecture and connections. In: Cerebral Cortex: Further aspects of cortical function, including hippocampus (Jones EG, Peters A, eds). Vol. 6, pp. 345–456., New York: Plenum Press.

Rouiller, E.M., Liang, F., Babalian, A., Moret, V., Wiesendanger, M., 1994. Cerebellothalamocortical and pallidothalamocortical projections to the primary and supplementary motor cortical areas: a multiple tracing study in macaque monkeys. J Comp Neurol 345, 185–213.

Saad, Z.S., Reynolds, R.C., 2012. Suma. Neuroimage 62, 768–773.

Sakai, S.T., Inase, M., Tanji, J., 1996. Comparison of cerebellothalamic and pallidothalamic projections in the monkey (Macaca fuscata): a double anterograde labeling study. J Comp Neurol 368, 215–228.

Saleem, K.S., Avram, A.V., Ye, F.Q., Yen, C.C., Komlosh, M., Basser, P.J., 2020. Multimodal high-resolution mapping of subcortical regions with MAP-MRI and histology., Organization for Human Brain Mapping, Virtual conference, p. 1768.

Saleem, K.S., Logothetis, N.K., 2012. A combined MRI and histology atlas of the rhesus monkey brain in stereotaxic coordinates., 2 ed. Elsevier/Academic press., San Diego.

Saleem, K.S., Price, J.L., Hashikawa, T., 2007. Cytoarchitectonic and chemoarchitectonic subdivisions of the perirhinal and parahippocampal cortices in macaque monkeys. J Comp Neurol 500, 973–1006.

Sato, F., Parent, M., Levesque, M., Parent, A., 2000. Axonal branching pattern of neurons of the subthalamic nucleus in primates. J Comp Neurol 424, 142–152.

Schaeffer, D.J., Selvanayagam, J., Johnston, K.D., Menon, R.S., Freiwald, W.A., Everling, S., 2020. Face selective patches in marmoset frontal cortex. Nat Commun 11, 4856.

Schaltenbrand, G., Wahren, W., 1977. Atlas for Stereotaxy of the Human Brain., 2 ed. Thieme;, Stuttgart, Germany.

Schmahmann J, Pandya D, 2009. Fiber Pathways of the Brain. New York: Oxford University Press.

Seidlitz, J., Sponheim, C., Glen, D., Ye, F.Q., Saleem, K.S., Leopold, D.A., Ungerleider, L., Messinger, A., 2018. A population MRI brain template and analysis tools for the macaque. Neuroimage 170, 121–131.

Setsompop, K., Fan, Q., Stockmann, J., Bilgic, B., Huang, S., Cauley, S.F., Nummenmaa, A., Wang, F., Rathi, Y., Witzel, T., Wald, L.L., 2018. High-resolution in vivo diffusion imaging of the human brain with generalized slice dithered enhanced resolution: Simultaneous multislice (gSlider-SMS). Magnetic Resonance in Medicine 79, 141–151.

Shah, A., Jhawar, S.S., Goel, A., 2012. Analysis of the anatomy of the Papez circuit and adjoining limbic system by fiber dissection techniques. J Clin Neurosci 19, 289–298.

Sidibe, M., Bevan, M.D., Bolam, J.P., Smith, Y., 1997. Efferent connections of the internal globus pallidus in the squirrel monkey: I. Topography and synaptic organization of the pallidothalamic projection. J Comp Neurol 382, 323–347.

Smith, J.B., Klug, J.R., Ross, D.L., Howard, C.D., Hollon, N.G., Ko, V.I., Hoffman, H., Callaway, E.M., Gerfen, C.R., Jin, X., 2016. Genetic-Based Dissection Unveils the Inputs and Outputs of Striatal Patch and Matrix Compartments. Neuron 91, 1069–1084.

Stauffer, W.R., Lak, A., Yang, A., Borel, M., Paulsen, O., Boyden, E.S., Schultz, W., 2016. Dopamine Neuron-Specific Optogenetic Stimulation in Rhesus Macaques. Cell 166, 1564–1571 e1566.

Sternberger, L.A., Sternberger, N.H., 1983. Monoclonal antibodies distinguish phosphorylated and nonphosphorylated forms of neurofilaments in situ. Proc Natl Acad Sci U S A 80, 6126–6130.

Straub, S., Knowles, B.R., Flassbeck, S., Steiger, R., Ladd, M.E., Gizewski, E.R., 2019. Mapping the human brainstem: Brain nuclei and fiber tracts at 3 T and 7 T. NMR Biomed 32, e4118.

Sullivan, C.R.P., Olsen, S., Widge, A.S., 2021. Deep brain stimulation for psychiatric disorders: From focal brain targets to cognitive networks. Neuroimage 225, 117515.

Tang, Y., Sun, W., Toga, A.W., Ringman, J.M., Shi, Y., 2018. A probabilistic atlas of human brainstem pathways based on connectome imaging data. Neuroimage 169, 227–239.

Tellmann, S., Bludau, S., Eickhoff, S., Mohlberg, H., Minnerop, M., Amunts, K., 2015. Cytoarchitectonic mapping of the human brain cerebellar nuclei in stereotaxic space and delineation of their co-activation patterns. Front Neuroanat 9, 54.

Thangavel, R., Sahu, S.K., Van Hoesen, G.W., Zaheer, A., 2009. Loss of nonphosphorylated neurofilament immunoreactivity in temporal cortical areas in Alzheimer’s disease. Neuroscience 160, 427–433.

Tournier, J.D., Calamante, F., Connelly, A., 2012. MRtrix: diffusion tractography in crossing fiber regions. Int J Imaging Syst Technol 22, 53–66.

Truong, T.-K., Darnell, D., Song, A.W., 2014. Integrated RF/shim coil array for parallel reception and localized B0 shimming in the human brain. Neuroimage 103, 235–240.

Turchi, J., Chang, C., Ye, F.Q., Russ, B.E., Yu, D.K., Cortes, C.R., Monosov, I.E., Duyn, J.H., Leopold, D.A., 2018. The Basal Forebrain Regulates Global Resting-State fMRI Fluctuations. Neuron 97, 940–952 e944.

Vitek, J.L., Johnson, L.A., 2019. Understanding Parkinson’s disease and deep brain stimulation: Role of monkey models. Proc Natl Acad Sci U S A.

Wells, A.M., Garcia-Cabezas, M.A., Barbas, H., 2020. Topological atlas of the hypothalamus in adult rhesus monkey. Brain Struct Funct 225, 1777–1803.

Westin, C.F., Maier, S.E., Mamata, H., Nabavi, A., Jolesz, F.A., Kikinis, R., 2002. Processing and visualization for diffusion tensor MRI. Med Image Anal 6, 93–108.

Wichmann, T., Delong, M.R., 2011. Deep-Brain Stimulation for Basal Ganglia Disorders. Basal Ganglia 1, 65–77.

Xiao, Y., Zitella, L.M., Duchin, Y., Teplitzky, B.A., Kastl, D., Adriany, G., Yacoub, E., Harel, N., Johnson, M.D., 2016. Multimodal 7T Imaging of Thalamic Nuclei for Preclinical Deep Brain Stimulation Applications. Front Neurosci 10, 264.

Yoshida, A., Ye, F.Q., Yu, D.K., Leopold, D.A., Hikosaka, O., 2021. Visualization of iron-rich subcortical structures in non-human primates in vivo by quantitative susceptibility mapping at 3T MRI. Neuroimage 241, 118429.

Yushkevich, P.A., Piven, J., Hazlett, H.C., Smith, R.G., Ho, S., Gee, J.C., Gerig, G., 2006. User-guided 3D active contour segmentation of anatomical structures: significantly improved efficiency and reliability. Neuroimage 31, 1116–1128.

